# Rapid and accurate genotype imputation from low coverage short read, long read, and cell free DNA sequence

**DOI:** 10.1101/2024.07.18.604149

**Authors:** Zilong Li, Anders Albrechtsen, Robert W Davies

## Abstract

Inexpensive and accurate genotyping methods are essential to modern genomics and health risk prediction. Here we introduce QUILT2, a scalable read-aware imputation method that can efficiently use biobank scale haplotype reference panels. This allows for fast and accurate imputation using short reads, as well as long reads (e.g. ONT 1X r^2^ = 0.937 at common SNPs), linked-reads and ancient DNA. In addition, QUILT2 contains a methodological innovation that enables imputation of the maternal and fetal genome using cell free non-invasive prenatal testing (NIPT) data. Using a UK Biobank reference panel, we see accurate imputation of both mother (r^2^ = 0.966) and fetus (r^2^ = 0.465) at 0.25X (fetal fraction of 10%, common SNPs). Imputation gets increasingly accurate as coverage increases, with r^2^ of around 0.90 or above for both mother and fetus at 4.0X (mother r^2^ = 0.996, fetal r2 = 0.894). We show that this imputation enables powerful GWAS and accurate PRS for both mother and fetus, which creates both clinical opportunities, and if phenotypes can be collected alongside clinical NIPT, the potential for large GWAS.

## Introduction

Low-coverage whole genome sequencing (lc-WGS) was originally demonstrated to be a cost-effective alternative to DNA genotyping arrays for genotyping in 2012^1^. However, early use of lc-WGS used statistical methods for phasing and imputation originally designed for microarrays^2^. Subsequently, efficient dedicated methods for imputation from lc-WGS were designed, which could operate both with or without reference panels, which were generally shown to be more powerful than approaches using arrays^3–6^.

Previously, we developed QUILT for rapid genotype imputation of lc-WGS data with a reference panel^3^. For both human and animal studies, QUILT has been shown to be the most accurate and robust approach when a reference panel is available ^6,7^. In addition, QUILT, by natively operating on individual sequencing reads, is able to accurately impute using long reads, or using reads with linkage information^4,8^. However, QUILT uses an approach to haplotype selection that has linear computational complexity in the number of haplotypes in the reference panel. As panels have grown in size, from roughly sixty thousand in the haplotype reference consortium (HRC), to several hundred thousand with the UK Biobank, QUILT becomes increasingly slow. Further, as haplotype reference panels are more consistently being generated through high coverage whole genome sequencing (WGS), with far greater SNP density, QUILT, with linear computational complexity in the number of SNPs, has been further slowed down.

For human GWAS, studies have estimated sample sizes in the millions are required to robustly detect signals that explain most of the heritability underlying common traits and diseases ^9^. This problem is only exacerbated when we consider the need for GWAS in many different populations, to enable population specific effect size estimation, and to help realize the promise of precision medicine for all^10^. One source of data that could help facilitate these large GWAS is using data from cell-free DNA (cfDNA) from non-invasive prenatal testing (NIPT). NIPT is a sensitive and specific screening test which tests for the common fetal aneuploidies of trisomies 13, 18 and 21, and that, according to the American Society of Obstetricians and Gynecologists (ACOG, 2020), should be offered for all pregnancies regardless of risk. In some countries, NIPT is routinely offered to all pregnant women, for instance in the Netherlands, with an average uptake rate of 46%^11^. In addition, in the first study to leverage lc-WGS NIPT data for GWAS, they suggested that over seven million pregnant women had already undergone NIPT with lc-WGS (0.1×-0.3×) in China by 2018 ^12^. Separately, Brand et al. recently proposed high-resolution noninvasive prenatal screening to investigate the entire fetal exome from circulating cfDNA by whole exome sequencing (WES) ^13^, which usually offers genome-wide off-target reads for non-coding regions for 1×-2× coverage ^14^. The increased benefit of such a test over screening only for aneuploidies should increase the rate of adoption of such tests worldwide. However, state-of-the-art imputation methods for lc-WGS are designed for diploid samples, i.e. where the sequencing reads come from two haplotypes ^3–6^, and there is no dedicated method designed for utilizing the triploid nature of NIPT data, i.e. where the sequencing reads come unequally from the maternal transmitted, maternal untransmitted, and paternal transmitted haplotypes. Given different timepoints of sequencing cfDNA in the blood of pregnant women, the fetal fraction (FF) varies between 4% and 50%^13^, and at the most commonly assayed gestational ages of 10 to 20 weeks, is about 10∼15%^15^. As such, the underlying data suggests that imputation from NIPT can work, and the scale of NIPT suggests it would be very valuable. Furthermore, imputing fetal genomes before childbirth may facilitate health risk management through polygenic risk scores (PRS) for perinatal traits in clinical practice.

In this study, we present QUILT2, a novel scalable method for rapid phasing and imputation from lc-WGS and cfDNA using very large haplotype reference panels. QUILT2 contains three key innovations, two technical and one methodological, compared to QUILT (or QUILT1). First, we introduce a memory efficient version of the positional burrows wheeler transform (PBWT) ^16^, which we call the multi-symbol PBWT (msPBWT). QUILT2 uses msPBWT in the imputation process to find haplotypes in the haplotype reference panel that share long matches to imputed haplotypes with constant computational complexity, and with a very low memory footprint. Second, we introduce a two stage imputation process, where we first sample read labels and find an optimal subset of the haplotype reference panel using information at common SNPs, and then use these to initialize a final imputation at all SNPs. This both speeds up imputation, as well as decreases the amount of RAM required. Finally, we introduce a novel methodological innovation to QUILT2, which we call the QUILT2-nipt method, which assumes the observed sequencing reads are from the three haplotypes present in NIPT data, and which imputes both the maternal and fetal genomes using DNA from pregnant women. In what follows, we evaluated the accuracy, performance and scalability of QUILT2 for various data types on diploid samples. In addition, we evaluated the accuracy of imputing the maternal and fetal genomes from simulated NIPT data. Finally, we show how this data can enable powerful GWAS and accurate PRS, in both the mother and the fetus.

## Results

### Model overview

QUILT2 is based upon, and improves, our previous model QUILT1 for diploid genotype imputation, which operates, which uses an iterative approach, as follows. First, Gibbs sampling is performed to generate a posterior estimate of the read labels (an estimate of what haplotype each read comes from) given the sequencing reads and the haplotypes in a small haplotype reference panel using the Li and Stephens algorithm^17^. Second, the small haplotype reference panel is updated based on the current estimate of the haploid dosages, again using Li and Stephens. Because of the second part of the iterative process, QUILT1 has linear time complexity in the number of haplotypes and variants in the full reference panel. For RAM and speed reasons, in QUILT1, we disable transitions except between every 32nd pair of SNPs, and store haplotypes as 32 bit integers (or a further representation of this). We refer to 32 consecutive SNPs as a grid.

With QUILT2, we first addressed the scaling of QUILT1. As illustrated in **Figure 1**, we implemented an analogue of the regular binary PBWT, which we call the multi-symbol PBWT (msPBWT), which naturally operates on the symbols in the grids used in QUILT1 and QUILT2 (**Extended Online Figure 1**). With msPBWT, and using pre-computed indices of the reference panel, we use an algorithm to identify haplotypes in the full reference panel that have long identical matches against a target imputed haplotype with computational time independent to the panel size. Next, we note with WGS derived reference panels, most SNPs are very rare, and as such, will slow down methods that are linear in the number of SNPs. In addition, these rare SNPs are of marginal importance in determining which reference haplotype to copy from when there are centimorgan or longer haplotype matches between target and reference haplotype. As such, we introduce a further two step process, where we iteratively perform imputation using only common SNPs and the reads that intersect them, to get an optimal subset of the full reference panel (**Extended Online Figure 2**). We then load all reads at all sites, initialize their phase according to the current best estimate of the underlying haplotypes, and perform a final round of Gibbs sampling, to get final read labels, and from them haplotype phase and genotype dosage. Finally, methodologically, QUILT2 introduces a novel method for jointly imputing the mother and fetus utilizing the lc-WGS cfDNA data from NIPT. This novel method includes changes to the mathematics as compared to the original “diploid” model, for instance the probability of the reads given the read labels and the haplotypes, as well as the probability of the read labels given the fetal fraction, to accommodate the three haplotypes present in different frequencies in cfDNA. Further, the change in the prior on read labels, which goes from uniform to unequal and dictated by the fetal fraction, was incorporated throughout the modeling, which necessitated substantial changes to the heuristics to minimize the Gibbs sampling getting stuck in local extrema. Further details, including for msPBWT, common variant initialized imputation, NIPT, as well heuristics, are described in **Methods** and **Supplementary Note**.

**Figure 1.**
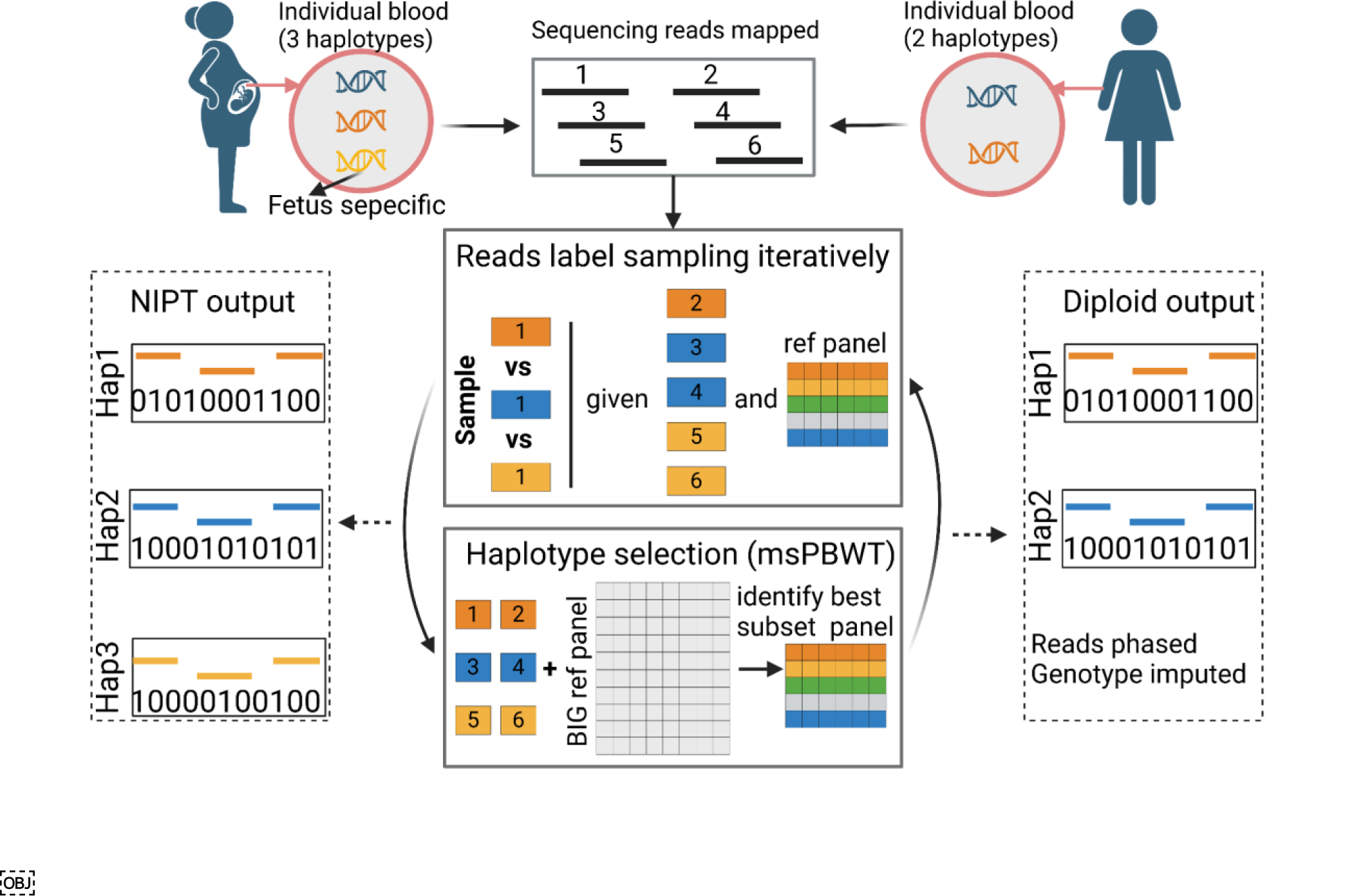
Schematic of the QUILT2 model. The model has two modes, called diploid and nipt mode, where the obvious difference is the number of underlying haplotypes modeled (2 vs 3). For both modes, QUILT2 first partitions the reads into different sets using Gibbs sampling and performs haploid imputation using a subset of reference haplotypes, and then obtains a new subset of the reference panel using msPBWT and the full reference panel. These two procedures are done iteratively several times to reach the final optimal phasing and imputation. Through this process, the input sequencing reads (unlabeled, black) are phased into different haplotypes (labeled, orange, blue, yellow), and genotypes are imputed and phased. More details about the msPBWT algorithm are shown in **Extended Online Figure 1**. Details about the iterative approach using common and all SNPs, which is not shown in this schematic, is shown in **Extended Online Figure 2**.

**Figure 2.**
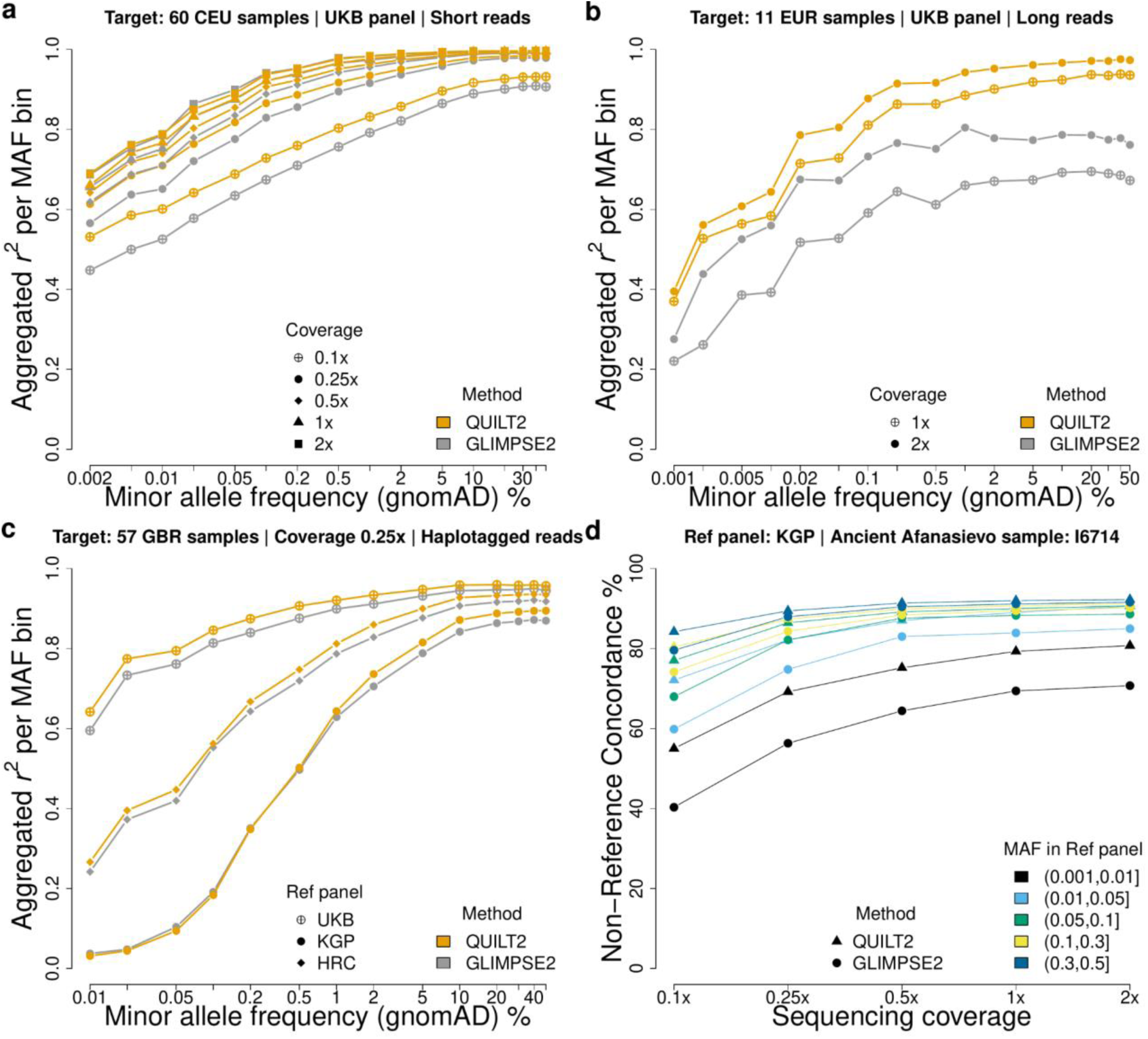
**Diploid imputation on various data types. a, b, c**, Imputation performance for short a), long b) and linked c) short read data on European samples across different reference panels and sequencing coverages for QUILT2 (orange) and GLIMPSE2 (gray). Accuracy, measured as the Pearson correlation coefficient (*r*^2^) between imputed dosage and true genotypes, is stratified by MAF. **d,** Imputation performance, measured as non-reference concordance rate, on aDNA sample from Afanasievo (∼4.6kya) across various sequencing coverages and MAF in the KGP panel.

### Diploid imputation performance

In the benchmarking of diploid imputation performance, we predominantly analyzed European samples in the 1000 Genomes and used chromosome 20 for convenience, and unless noted, focus on the analysis with three different reference panels: the 1000 Genome Project (KGP, *N (haps)*=5,008, *M (SNPs)*=2,115,074), the Haplotype Reference Consortium (HRC, *N*=54,330, *M*=884,932), and the UK Biobank 200K WGS genomes (UKB, *N*=400,022, *M*=14,075,021). We first investigated the imputation performance of QUILT2 and GLIMPSE2 for diploid genomes using various data types with multiple sequencing coverages. Throughout the results, unless noted, we stratified accuracy of European samples by allele frequencies from the separate Genome Aggregation Database (gnomAD v3.1.2 Non-Finish European). As exemplars, we used ‘very rare’ to refer to SNPs with MAF of 0.01-0.02%, ‘rare’ to refer to SNPs with MAF of 0.1-0.2%, and ‘common’ to refer to SNPs with MAF of 10-20%. We first considered imputation using short read Illumina data from the CEU population of the 1000 Genomes. As shown in Figure 2A and **Supplementary Figure 1**, QUILT2 was more accurate than GLIMPSE2 at sequencing coverage ≤0.5×, regardless of the reference panel, and particularly at 0.1×-0.25×. For higher coverage samples ≥1×, at common SNPs, accuracy was similar between the panels and between the methods, while at rarer SNP, far more rare SNPs were imputed, and more accurately imputed with the larger panels (**Supplementary Figure 1, Supplementary Table 1, 2**). No method was uniformly superior for these rare SNPs at higher coverage samples, with QUILT2 being slightly more accurate than GLIMPSE2 for the larger panel (UKB) but not for the smaller panels (KGP and HRC). For QUILT2, we also evaluated the effect of the msPBWT haplotype selection versus the Li and Stephens approach of QUILT1, and note the msPBWT approach generates slightly more accurate results (**Supplementary Figure 2**). In terms of phasing, the larger UKB panel substantially improved QUILT2 phasing accuracy compared to the smaller panels (**Extended Online Figure 3, Supplementary Figure 3**), and mirroring the imputation results, phasing with QUILT2 was more accurate than GLIMPSE2 particularly for lower coverages.

**Figure 3.**
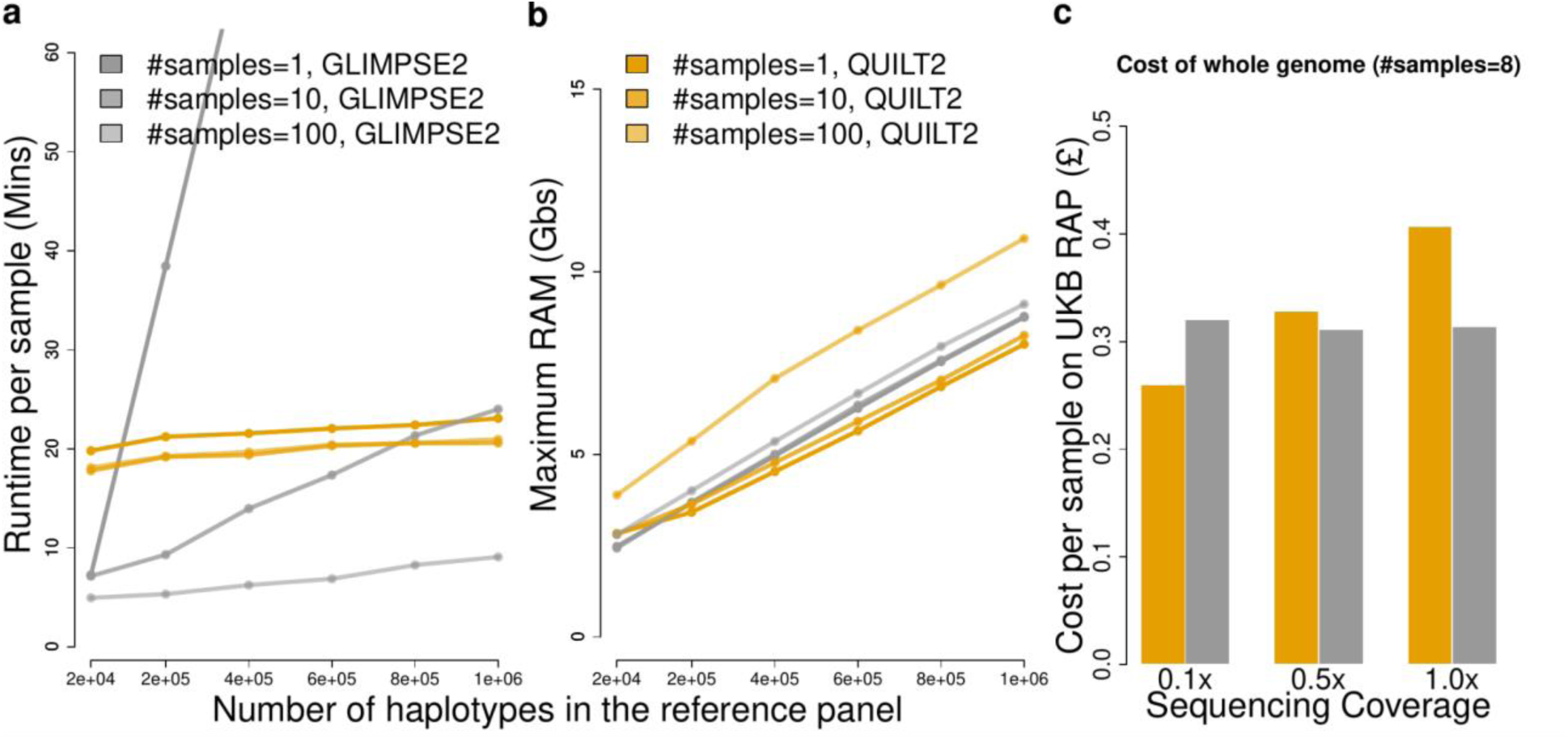
Scalability and computational cost. The benchmarking for QUILT2 (orange) and GLIMPSE2 (gray) was performed on virtual machines of type mem2_ssd1_v2_x4 with 4 cores and 16 GBs memory on the RAP. Regarding the scalability **a,b**, six different sizes of reference panel and three different sizes of input samples were tested. The number of SNPs in the reference panels varies, reflecting the removal of SNPs that become monomorphic in a subset of the panel. Time and RAM were measured for each chunk on chr20 by “/usr/bin/time -v” and one thread was used here. **c**, for estimating the real cost on RAP, both QUILT2 and GLIMPSE2 run parallelly with 4 cores to maximize the usage of available resources, but there were two big chunks (>10Mb) that fell back to 2 cores for QUILT2 due to out-of-memory. We note the chunks (>5Mb) determined by the *GLIMPSE2_chunk* were used for comparison, but QUILT2 works best with smaller chunks (e.g. 3Mb) determined by the *quilt_chunk_map* function. The cost of the whole genome was estimated based on the bill of imputing chr20 by multiplying a factor of 34.48.

We also evaluated the imputation accuracy of QUILT2 using other data sources. We evaluated the imputation accuracy using long sequencing reads from Oxford Nanopore Technologies (ONT) by calculating both the *r*^2^ at variant level and F1-score at sample level. Across all coverages and reference panels, QUILT2 was substantially more accurate compared to GLIMPSE2 for long reads (**Figure 2B, Supplementary Table 3**), e.g common *r*^2^=0.937 vs *r*^2^*=*0.695 at 1× coverage. Similarly, we saw a consistent benefit of QUILT2 versus GLIMPSE2 when using haplotagged data, a low cost method for generating linked read information for short reads ^18^ (**Figure 2C**). As with the regular short read data, imputation was most accurate when using the UKB panel, followed by HRC and KGP (**Figure 2C**). In particular, the UKB panel can boost the accuracy for very rare variants (*r*^2^=0.83, coverage=1×). Lastly, we assessed the imputation performance on ancient DNA (aDNA) with three reference panels. Interestingly, QUILT2 outperformed GLIMPSE2 across all frequency variants using KGP and HRC panel, while both performed similarly using the UKB panel (**Figure 2D, Supplementary Figure 4**). Similar to results for modern DNA, with QUILT2, the imputation of aDNA showed similar accuracy for common variants across reference panels, while the UKB panel again improved the performance of rare variants.

### Scalability and computational cost

We next evaluated the computational cost and scalability of QUILT2 along with GLIMPSE2 for both speed and memory usage, varying both the size of the haplotype reference panel, as well as the number of target samples to run. As shown in **Figure 3A**, thanks to msPBWT, QUILT2 has approximately constant runtime with respect to haplotype reference panel size, and the per-sample runtime is not affected appreciably by the input sample size. By contrast, the per-sample runtime of GLIMPSE2 is affected by the number of samples run, and only shows approximately constant computational complexity with respect to haplotype reference panel size for larger number of input samples (n>100) but not for smaller size, as has been previously noted by the authors of GLIMPSE2 ^6^. In the extreme case, when only imputing one sample at a time with around 1 million haplotypes in the panel, QUILT2 would be an order of magnitude faster than GLIMPSE2. In terms of memory usage, we note that when using msPBWT, QUILT2 used a maximum of 20 GBs of RAM for *N*=400,000 haplotypes in the reference panel with chunk size over 5 Mb on average, while QUILT1 used around 160 GBs. Further, QUILT2 with both msPBWT and the two-stage imputation enabled was ∼3× faster and used ∼4× less RAM, than QUILT2 with only msPBWT (**Supplementary Figure 5**). Comparing QUILT2 and GLIMPSE2 for RAM, we see that both approaches show linear increases in RAM with reference panel size (**Figure 3B**). Given around 1 million reference haplotypes and an average chunk size of around 5Mb and 500Kb buffer to impute on chromosome 20, the maximum RAM for both is less than 16 GBs. We note that QUILT2 uses slightly higher RAM given 100 samples to run, and QUILT2 also runs faster with lower RAM for lower coverage data (**Figure 3C, Supplementary Figure 5, 6**). Since the UKB panel is only accessible on the UK Biobank research platform (RAP), we also assessed the cost of imputing the whole genome on the RAP. As shown in **Figure 3C**, given 8 samples as input, QUILT2 costs £0.260, £0.328 and £0.407 to impute the whole genome for coverage 0.1×, 0.5× and 1.0× respectively, while GLIMPSE2 costs around £0.315 for all coverages equally.

### NIPT cfDNA imputation performance

We next investigated the QUILT2-nipt method for the maternal and fetal genome imputation using cell-free DNA sequencing data from NIPT. We simulated 30 NIPT samples by mixing the sequencing reads of the mother and child in the 30 CEU trios from KGP (**Methods**). Here we focus on NIPT imputation using the UKB panel across different sequencing coverages and various fetal fractions (FF). Firstly, we compared the QUILT2-nipt method with the existing diploid method (i.e. QUILT2-diploid) for maternal genotype imputation. As shown in **Figure 4A and Supplementary Table 4**, given FF=0.1, there is a clear gain of accuracy in QUILT2-nipt over QUILT2-diploid (0.25×, very rare *r*^2^=0.724 vs 0.695; 1.0×, very rare *r*^2^=0.831 vs 0.799; 2.0×, very rare *r*^2^=0.851 vs 0.839). At higher sequencing coverage, for higher fetal fractions, QUILT2-diploid performed increasingly poorly, while QUILT2-nipt still maintained high accuracy for common variants as expected (e.g. 4.0×, FF=0.2, common *r*^2^=0.994, 0.853 for QUILT2-nipt and QUILT2-diploid respectively).

**Figure 4.**
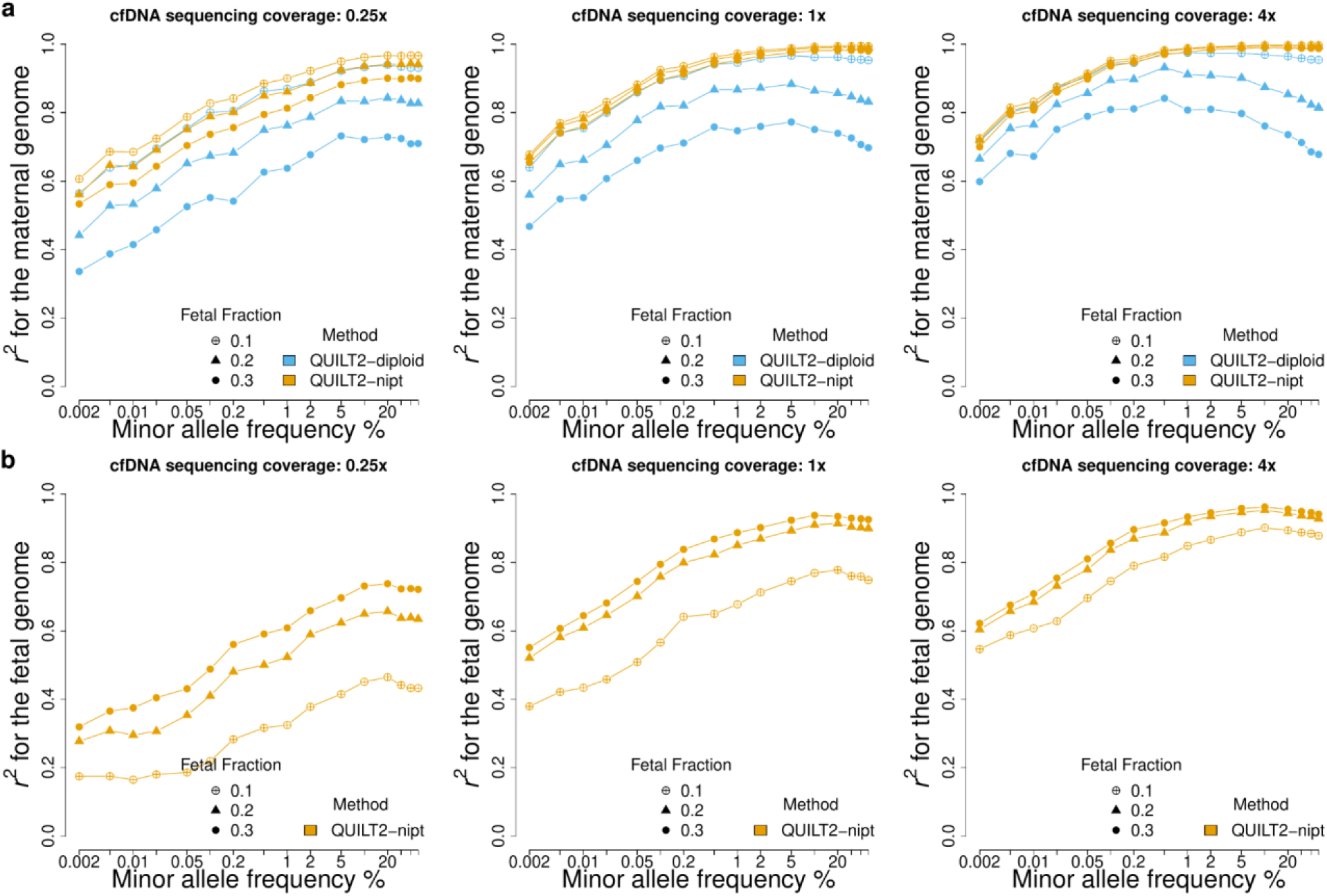
Maternal and fetal genome imputation from cfDNA. The simulated 30 NIPT samples from CEU trios were analyzed. **a**, imputation performance for the maternal genome across different NIPT sequencing coverages and various fetal fractions. Both QUILT2-diploid and QUILT2-nipt are able to generate the imputed genotypes of the mothers. **b**, imputation performance for the fetal genome, where only QUILT2-nipt can impute the fetal genome.

Next we investigated the performance of QUILT2-nipt on fetal genotype imputation where there are no other approaches for comparison. As shown in **Figure 4B** and **Supplementary Table 5**, both increasing fetal fraction, as well as sequencing coverage, contribute to improved accuracy, at both rare and common SNPs. For instance, at 0.25× coverage, accuracies at common SNPs improve with higher FF (*r*^2^ = 0.465, 0.657, 0.783 for FF = 0.1, 0.2, 0.3 respectively), and they improve further at higher coverage with 4.0x (*r*^2^ = 0.894, 0.944 and 0.955, same FFs). However, we do not observe a gain in accuracy when the sequencing coverage exceeds 4.0×, assuming FF=0.3 (**Supplementary Table 5**). We note that *r*^2^*=0.955* appears to be the upper limit for fetal genotype imputation using the UKB panel, versus 0.92 and 0.90 for the HRC and KGP panels, respectively (**Supplementary Figure 7)**. This contrasts with the performance for the mother, where the upper limit remains 1.0 regardless of the reference panel. This improved performance for the larger panel is driven by the improved phasing capabilities offered by the larger reference panel (**Extended Online Figure 3**), enabled by msPBWT within QUILT2, which is especially important for phasing the reads from the paternally transmitted haplotype. In terms of the run time, where mother and fetus are imputed together, we note that the QUILT2-nipt method is about 30% slower than the QUILT2-diploid method, and imputing only common variants can reduce both the runtime and memory substantially (**Supplementary Figure 6**).

### Power in association testing and prediction with cfDNA

Finally, we explored the useability of imputed results from QUILT2-nipt for both GWAS and PRS. To create realistic NIPT sequencing samples, we used UK Biobank samples, for which we have real phenotypic data, and used parent offspring pairs to represent the mother and the fetus (**Methods**). We used 11,028 parent-offspring pairs and simulated mean sequencing coverages of 0.25×, 1.0×, and 2.0×, with fetal fractions being simulated with mean 10% and following a normal distribution, with range from 5% to 15% (**Supplementary Figure 8**). We conducted association tests on chromosome 1 for 25 quantitative traits using a linear mixed model on these 11,028 individuals (**Methods**).

We compared the GWAS association findings using QUILT2-nipt imputed maternal and fetal genotype dosages to a baseline that used imputed genotype dosages from arrays. With the QUILT2-nipt imputed data, we found that at common SNPs the *r*^2^ for maternal genotypes were 0.973, 0.994, 1.000 at 0.25×, 1.0×, 2.0× coverage respectively, and the common *r*^2^ for the fetal genotypes were 0.480, 0.75 and 0.82 (**Figure 5A**). GWAS summary statistics based on different imputed datasets did not show signs of inflation of test statistics as noted by Q-Q plots (**Supplementary Figure 9**). Next, we assessed the power of the different coverages and approaches, by measuring how many of 615 independent GWAS signals on chromosome 1 found using the entire UK Biobank participants could be identified (*p* < 0.05) using the imputed datasets of 11,028 simulated NIPT samples (**Methods**). Using the array based imputation, 190 associations were found, while for QUILT2-nipt maternal imputation, 190 (100% estimated relative power), 188 (99%) and 181 (95%) were found at 2.0×, 1.0×, and 0.25× respectively. For QUILT2-nipt fetal imputation, 163 (86%), 159 (84%) and 106 (56%) of loci were validated for 2.0×, 1.0×, and 0.25× data respectively (**Figure 5B**). Notably, even though the common *r*^2^ for fetal genotypes was only 0.82 at 2.0× with 10% FF, we still obtained a relative power of 86% in the GWAS. This is further validated in an error-prone setting where the estimates of fetal fraction are measured with uncertainty and thus the prior for QUILT2-nipt method is mis-specified (**Supplementary Figure 10**).

**Figure 5.**
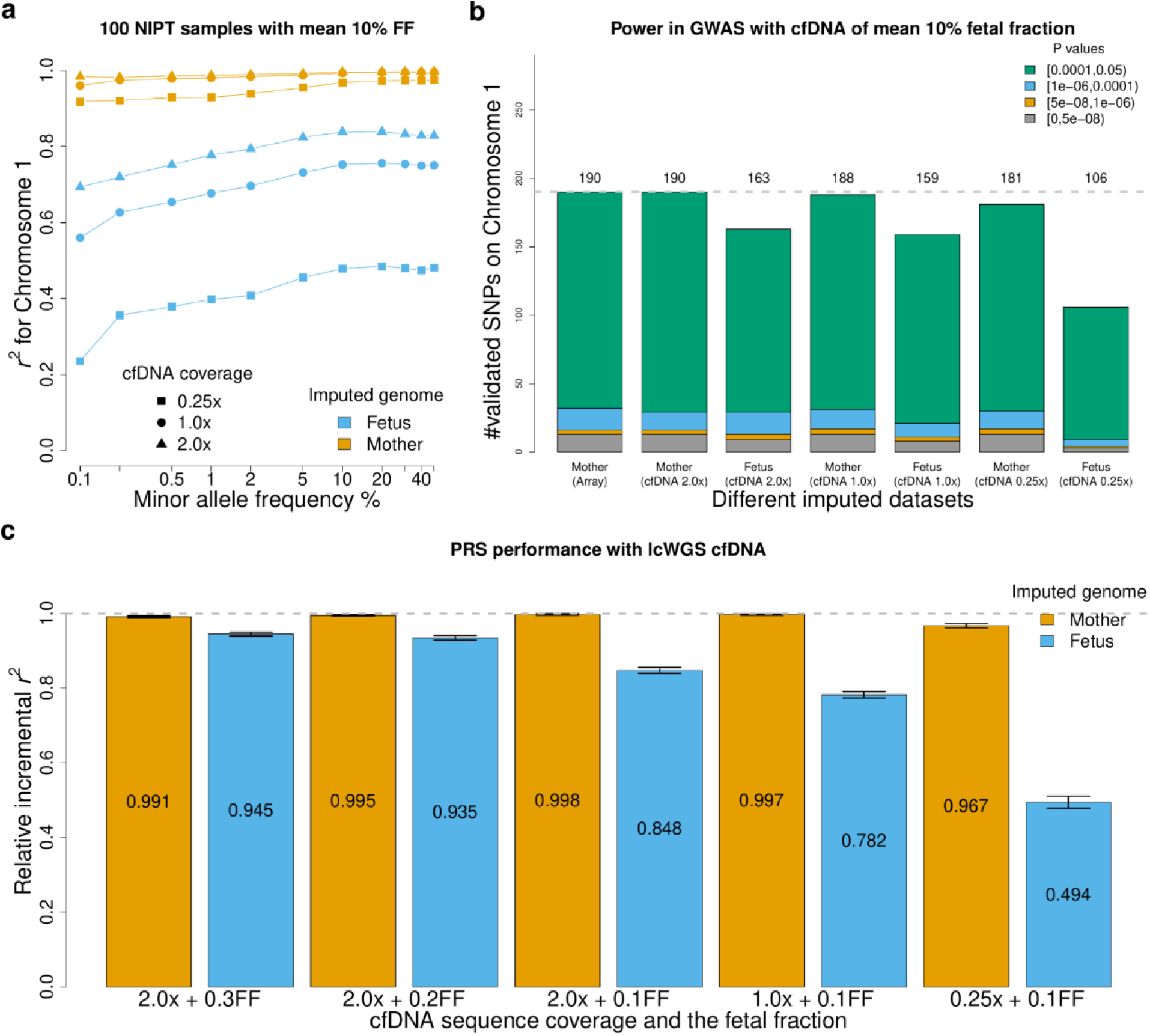
Power in GWAS and PRS with cfDNA. a, imputation accuracy of both maternal and fetal genotypes on chromosome 1 using the UKB panel for different cfDNA sequencing coverage with mean 10% fetal fraction. **b,** Power in GWAS with both imputed mother and fetus genotype, compared the array-imputed data. Power is measured as the number of independent GWAS hits that could be validated (*p* values < 0.05) using the target imputed datasets of 11,028 samples. **c,** PRS performance with the whole imputed fetal and maternal genome from cfDNA with various sequencing coverage and fetal fractions. The mean value of relative incremental *r*^2^ is shown at the middle of each bar.

In addition to GWAS, the fetal imputed genotypes from QUILT2-nipt can be used to generate fetal polygenic risk scores, which can have potential clinical value for perinatal traits. We therefore evaluated the performance of PRS using QUILT2-nipt imputed fetal genotype dosages for 1000 individuals (**Methods**). We examined 21 traits with incremental *r*^2^ > 0.1 (except for the waist-hip ratio) in unrelated white British Europeans^19^, which is the difference in *r*^2^ between the full prediction model (including PRS) versus the model with covariates alone (age, sex, age×sex, assessment center, genotyping array, 10 PCs). Using the reported PRS weights of each SNP, we calculated the PRS for each individual given different imputed datasets, that is the array-imputed, as well as NIPT imputed from 2.0×, 1.0×, and 0.25× coverage. We compared the incremental *r*^2^ of the NIPT imputed datasets to the array imputed datasets, referring to this comparison as the relative incremental *r*^2^ (**Methods**). As shown in **Figure 5C** and **Supplementary Figure 11**, with effective sequencing coverage for the mother at least 0.2× (e.g. 0.25×, 10% FF), we observed a relative incremental *r*^2^ > 0.967. For the fetal PRS, while at low effective sequencing coverage, relative incremental *r*^2^ is modest (e.g. 0.25×, 10% FF, giving 0.025× coverage, *r*^2^=0.494), at high effective sequencing coverage (e.g. 2.0×, 20% FF, giving 0.4×), we observed a high relative incremental *r*^2^ > 0.935. As expected, the PRS performance is aligned with the imputation accuracy.

## Discussion

Here, we introduced QUILT2, a novel method for imputation from lc-WGS. QUILT2 includes both technical improvements, which increase the speed and scalability versus our previously released method QUILT1, as well as a methodological improvement to allow it to impute both mother and fetus from NIPT data. For diploid samples, we used the WGS derived UKB panel to show that QUILT2 is approximately as accurate as QUILT1, but substantially faster, and uses much less memory, and as such, should be much less expensive in cloud based applications. We further showed that QUILT2 is about as accurate as GLIMPSE2, being more accurate in certain situations, like lower coverages, and with long or linked read sequencing data. The latter point suggests QUILT2 should retain greater accuracy over GLIMPSE2 in various non-human settings, as the effective population size, and hence SNP density, is often much higher than in humans. In addition, while QUILT2 is slower than GLIMPSE2 when processing many samples, it is faster, by an order of magnitude on very large reference panels, when processing only a very small number of samples. As such, QUILT2 offers a benefit over GLIMPSE2 in analysis pipelines geared towards processing samples individually or in small batches, as might happen if imputation is performed immediately after data acquisition. Importantly, we also showed the extent to which large haplotype reference panels can improve both phasing and imputation performance, showing a substantial improvement in the overall accuracy and the number of well imputed rare variants.

In addition, we showed with QUILT2-nipt that QUILT2 can impute both mother and fetus using large reference panels. We showed that QUILT2-nipt can achieve very high accuracy at common variants in the mother at moderate effective sequencing coverage (e.g. 1×, 0.1 FF). For the fetus, we showed that PRS and GWAS performance are reasonable at moderate effective sequencing coverage (e.g. 1×, 0.1 FF), and they become strong with small increases (e.g. 2×, 0.1FF). Given the decrease in sequencing cost in recent years, it is not much more expensive to generate lc-WGS from NIPT at 2× versus the more modest levels common today. Even so, the accuracy of the QUILT2-nipt method at the more modest coverage should allow researchers to leverage existing large NIPT data to conduct GWAS both generally, as well as in specific contexts, such as to help unravel the complicated mechanisms underlying the regulation of human parturition, where both the maternal and fetal genome are involved ^20–23^. From a GWAS perspective, the value of maternal and fetal genotypes can be enhanced through phenotypes acquired through linkage with electronic health records, or through surveys of affected family members, using approaches like GWAS-by-proxy ^24^ and LT-FH ^25^. From a clinical perspective, as we move toward leveraging PRS in health risk management, the imputation of the fetal genome before childbirth should afford the opportunity to change clinical practice for relevant traits, such as gestational hypertension and preeclampsia ^26,27^.

We note that QUILT2 contains limitations and areas for future improvements. First, it should be possible to model recombination in the maternal haplotypes in NIPT. Although we expect not modeling this to have a small effect for low fetal fractions, and can be mediated by imputing in small genomic windows, future methodological innovations could directly account for this by modeling both maternal haplotypes, and then using a hidden variable reflecting which of the two haplotypes is the maternally inherited one in the fetus. Second, there is clinical value in the detection of *de novo* CNVs before birth. For example, detection of *de novo* CNVs at 22q11.2, which occurs in around 1/1500 - 1/2000 live births^28^, allows for delivery to take place at specialized medical centers, and for timely treatment of conditions like for neonatal hypocalcemia and immunodeficiency, which improves outcomes ^29,30^. Current methods to detect *de novo* copy number variation using NIPT focus on either read counting approaches in windows^31^, or machine learning approaches using massively multiplexed PCR^32^. By binning reads into sets reflecting their origin, and effectively using long range and external information, it is reasonable to believe an approach based on an extension of QUILT2 could have greater sensitivity and specificity compared to current approaches, and improve clinical utility. Third, QUILT2 inherently phases reads to their haplotypic background, and as such, with ONT long reads, a simple structural variants (SV) caller (e.g. large deletions, insertions) could be implemented easily. A more complicated, and likely more accurate, caller that modeled a copy number at each genomic location, as would an even more complicated approach that involved iterative re-mapping, would likely improve accuracy further. Finally, as we noted in QUILT1, large lc-WGS datasets offer the possibility to augment reference panels. While reference panels from commonly studied populations, particularly European ones, have large reference panels available, there would be tremendous value in leveraging large NIPT efforts to generate and use novel haplotype reference panels.

In conclusion, we expect QUILT2 to improve imputation of diploid samples in many settings. In addition, the imputation of cfDNA NIPT samples using QUILT2-nipt can enable the creation of large GWAS through recovery of both maternal as well as fetal genotypes, as well as offer healthcare opportunities, enabling prenatal PRS.

## Methods

### QUILT2

Details about the QUILT model have been previously published^4^. In brief, QUILT is a Gibbs sampler that samples read labels based on the observed data (i.e. sequencing reads) and parameters of the model (e.g. haplotype reference panel), and from this, allows for phasing and genotype imputation. Here, we provide information about the mathematical changes to QUILT2 that form the basis of the QUILT2-nipt method. We also give a brief introduction to msPBWT, as well as the iterative process that uses common SNPS then all SNPs. More details about msPBWT, as well as the heuristics used in Gibbs sampling for NIPT are given in the **Supplementary Note**.

In the NIPT mode, we used Gibbs sampling to generate draws from *H∼P(H|O, □, FF)*, where *H* is a vector of read (haplotype) membership labels, *O* is our reads (i.e. observed bases at SNPs), *FF* is the fetal fraction and □ is the parameters of our model. In NIPT, sequencing reads come from three sources, and let us label them as 1 = maternal transmitted, 2 = maternal untransmitted and 3 = paternal transmitted. As such, for each read *v,* we have the prior probability of haplotype membership as

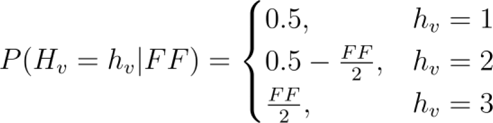

Let *o* be the realized observations for the random variable *O* (sequencing read bases and base qualities). Consider in the Gibbs sampling that we want to sample a new value for read indexed by *v* with read label *h_v_,* conditional on all other read labels. Let *H_v_* be this random variable at this point in the Gibbs sampler and let *H_-v_* be a random variable representing the remaining read labels. We therefore need to calculate:

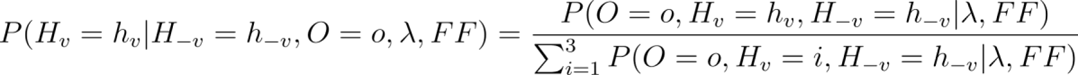

for 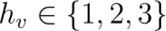 and sample *h_v_* using this probability. To do this we use 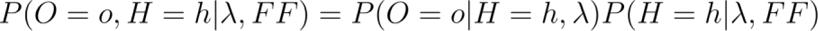 which uses the prior probability of all reads, which is the product of the prior for one read described above. This prior probability does not cancel like it did for QUILT1 (diploid) where 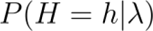 was constant regardless of *h_v_*. For 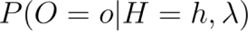 we note that given the vector of read labels, we split the reads into three sets reflecting their haplotypic origin, call that 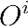, and can generate probabilities in the normal ways for HMMs, in an obvious extension to what is presented in the QUILT1 paper. For 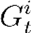 the haploid (genotype) for haplotype *i* for some SNP *t*, we can calculate the posterior probability that this is the alternate allele (encoded by a 1) as follows

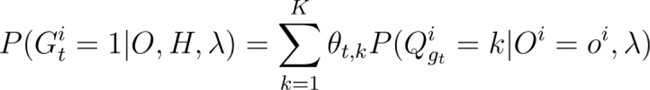

where, □*_t,k_* is the probability that reference haplotype *k* carries the alternate allele at SNP *t*, 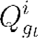 is the reference haplotype carried at grid *g_t_* for haplotype *i*, noting that 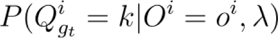 is a standard output from an HMM. From these probabilities, a haplotype dosage can be taken as the sum across several Gibbs samplings, and from this, genotype dosages can be taken for the mother by summing up the value for *i = 1* and *i = 2*, while for the fetus we choose *i = 1* and *i = 3*.

We have introduced the msPBWT algorithm in QUILT2 to find haplotype matches for target haplotypes against in the full reference panel, with computational complexity independent of the size of the full reference panel. An overview of msPBWT is given in **Extended Online Figure 1**. First, the haplotype reference panel, containing binary information about whether haplotypes carry reference or alternate alleles, is encoded, and then ranked. This ranked matrix contains columns with 255 symbols representing the most commonly seen 32 SNP haplotype in this window (grid), and a final symbol (0) representing a haplotype not amongst the 255 most commonly seen, for which information (the haplotypes carried and their positions) are stored in helper matrices. Next, two indices are built: *A*, which allows for indices to be looked up at any grid to put the original haplotype reference panel in reverse sorted prefix order; and *U*, which allows for determination between grids about how the ordering of indices changes. Both *A* and *U* can be used with a two part algorithm that allows for rapid determination of which haplotypes in the haplotype reference panel contain long exact matches to the target haplotype. First, a vector *f* is found, representing at each grid, where the target haplotype would be inserted into the haplotype reference panel to place the haplotype reference panel in reverse sorted prefix order. Second, scanning up and down is performed, to maintain a list of the closest *L* haplotypes to the target haplotypes. Finally, filtering is done with respect to a minimum length of match, and when more matches are identified than requested haplotypes, further filtering is done for the haplotypes identified for the two haplotypes, which optimizes both the length of matches, as well as their uniqueness along the target region. Full details of msPBWT are given in the **Supplementary Note**.

Finally, we note that with high coverage WGS based reference panels, the vast majority of variants are very rare, and thus are not informative early in the iterative process, as they offer little discriminatory information about which reads come from which haplotypes. As such, we introduce a further two-step process, where we first impute using a set of common variants and the reads that intersect them, to get an initial estimate of the carried haplotypes. We then load all reads at all sites, initialize their phase according to the current best estimate of the underlying haplotypes, and perform a final round of read-sampling (**Extended Online Figure 2**). Together, these changes made QUILT2 substantially faster and more memory efficient than QUILT1. Finally, we upgraded a heuristic, to perform efficient block Gibbs sampling between all sequential pairs of grids, to identify and solve phase switch errors with read labellings.

### Datasets

We created various test and truth datasets to benchmark all methods. We used three reference panels for testing genotype imputation and phasing accuracy, which are the 200K WGS derived UK biobank reference panel, the HRC and the 1000 Genomes Project. Additionally, for benchmarking the computational performance, we used the very big reference panel of *N*=976,630 haplotypes (488,315 individuals), that is the UK biobank data imputed by the Genomics England haplotype reference panel^33^, which we referred to as the UKB-GEL. We used liftover to convert the HRC reference panel from the GRCh37 to the GRCh38 build using the Genome Analysis Toolkit^34^ (GATK) Picard LiftoverVCF v2.22.2.

#### CEU

We downloaded the high coverage (>30x) CRAM files of 30 CEU trios in the 1000 Genome project, from which we called the true genotypes at sites in the UKB-GEL panel using bcftools^35^ v1.18 with option ‘call -Aim -C alleles -T ref.sites’. We filtered the true genotypes at sites with read depth lower than 10. Leveraging trios information, phasing was done first assuming Mendilian inheritance, excluding triple-heterozygous sites using bespoke R v.4.2.2 code; then, the excluded sites were phased with this scaffold using shapeit4 v4.2.2 ^36^.

#### GBR

We downloaded the low-coverage haplotagged data (∼0.25x) for 59 GBR samples in the 1000 Genome Project, which were generated in the QUILT1 paper^4^. When evaluating the imputation performance on haplotagged reads, we used the called genotypes from high coverage short Illumina reads as truth, which was done using the same bcftools command.

#### ONT

We downloaded high-coverage ONT alignment files of GRCh38 build (https://s3.amazonaws.com/1000g-ont/index.html?prefix=ALIGNMENT_AND_ASSEMBLY_DATA/FIRST_100/) for 11 samples in the EUR super-population from the first 100 ONT samples in the 1000 Genomes Project^37^. When evaluating the imputation accuracy with ONT reads, We used the called genotypes from high coverage short Illumina reads as truth, which was done using the same bcftools command.

#### Ancient DNA

We downloaded four high-coverage ancient DNA samples from the Afanasievo family (mother I3388, father I3950, son I3949, son I6714), who lived ∼4.6 thousand years ago (ka) in the Altai Mountains of Russia, with average coverages of 10.8×, 25.8×, 21.2×, and 25.3×, respectively, which have reliable genotype calls and phasing ^38^. Since both the public BAM and VCF files are of GRCh37 coordinates, we first liftovered the VCF with true genotypes to GRCh38 and converted the downloaded BAM files back to raw FASTQ using ‘samtools fastq’. Then we used the mapache^39^ snakemake workflow to map the FASTQ files against the human reference genome GRCh38 to generate the BAM files for imputation benchmarking.

#### NIPT cfDNA

We simulated the NIPT cfDNA using the real sequencing reads from trios or duos individuals. The details are described in the next section

### NIPT cfDNA simulations from real reads

We simulated 3 sets of NIPT cfDNA reads for different analyses using the real sequencing reads. First, to evaluate the QUILT2-nipt and QUILT2-diploid method on maternal genotype imputation, we simulated NIPT cfDNA with various fetal fractions by mixing the maternal reads and offspring reads in the 30 CEU trios from the 1000 Genome Project. And we only simulated BAM files for chromosome 20 for benchmarking purposes. Second, to evaluate the GWAS performance on NIPT samples and obtain as many samples as possible, we first identified 1060 trios and 4114 duos in the UK Biobank given the released kinship statistics^40^. Specifically, we defined the first degree relatives as the paris with kinship value >0.1767 and IBS0 value <0.0012. To further identify the trios and duos, we defined that the difference between the parents and offspring in age must be greater than 15. Therefore, given a list of trios and duos, we simulated 11,028 NIPT samples by treating the offspring as the parent regardless of sex and vice versa. And we only simulated the BAM files of 11,028 samples for chromosome 1 due to the limited budget and computational resources. Third, to evaluate the PRS performance that requires genotypes of the whole genome, we sampled only 1000 NIPT samples from the above identified duos and trios, which includes 500 unique parent-offspring pairs. For those 1000 NIPT samples, we simulated the BAM files for the whole genome.

### Imputation benchmarking workflow

To access the performance of QUILT2 and GLIMPSE2 on various datasets, we developed a reproducible and configurable snakemake workflow for imputation benchmarking (https://github.com/Zilong-Li/lcWGS-imputation-workflow), which includes downsampling BAM/CRAM files to low coverage; creating reference panels by removing the target samples from the input panel; parallelly running by chunks and ligating results; automated reproducible pipeline with support to High-Performance Computing clusters.

### Assessing imputation accuracy

To assess the imputation and phasing accuracy, we developed an efficient function *vcfcomp* in the vcfppR v0.4.6 package^41^, which can calculate various concordance metrics between two VCF/BCF files at either samples level or variants level. In the output VCF files, homozygous reference, heterozygous and homozygous alternative genotypes are coded as integer value 0, 1 and 2 respectively, while the genotype dosage are float values between 0 and 2. There are three concordance metrics defined here. First, we defined *r*^2^ as the squared Pearson correlation between the impute dosages (test) and high-coverage genotype calls (truth). Second, we defined the non-reference concordance (NRC) rate as NRC = 1 - (e0 + e1 + e2) / (e0 + e1 + e2 + m1 + m2), where e0, e1, e2 are the counts of mismatches for genotype 0, 1 and 2 respectively, while m1 and m2 are the counts of matches between the imputed and truth for genotype 1 and 2. Third, we defined the F1-score as F1 = 2 * TP / (2 * TP + FP + FN), where TP is the counts of matches between the imputed and truth for genotype 1 and 2, FP is the counts of mismatches for genotype 1 and 2, and FN is the counts of mismatches for genotype 0 and 1.

To assess genotype imputation accuracy at variant level, we first grouped SNP by its frequency intervals and aggregated the concordance metrics (e.g. *r*^2^) for all samples across all SNPs in that frequency interval. On the other hand, to assess the genotype imputation accuracy at the sample level, we only aggregated the concordance metrics (e.g. NRC, F1) for genotypes of a single sample at a given allele frequency interval. The allele frequencies were taken from either the gnomAD v.3.1.2 database (for modern European samples, https://gnomad.broadinstitute.org/downloads#v3) or from the reference panel itself (for ancient DNA).

### Assessing phasing accuracy

To assess phasing accuracy, we used phasing switch error (PSE) rates as follows, which is implemented in *vcfcomp* with option *stats=‘pse’*. First, consider that we have both true haplotypes (truth) and the imputed haplotype (test) at sites where the truth genotypes are heterozygous. Next, define as discordant any test sites that are also not heterozygous. On the remaining sites, define a phase switch error when either the true haplotypes record a change where the haplotype carries the alternate allele between adjacent heterozygous sites when the test haplotypes do not or vice versa. We removed from consideration sites that were flipped, that is, yielding consecutive phase switch errors. The PSE rate is the number of phase switch errors divided by the total number of pairs of consecutive heterozygous sites examined and can be combined across discrete imputed windows.

### Association testing and validations

Since the data consists of the related individuals from families, we used the linear mixed model implemented in gemma^42^ v0.98.5 for association tests accounting for the genetic relatedness. First, we computed the genetic relatedness matrix (GRM) using *‘gemma -gk 1’* based on the genotypes from the array with the default setting for SNP QC ( *-maf 0.01 -miss 0.05*). Then we ran association tests on multiple imputed datasets in BIMBAM format (for dosages) using *‘gemma −lmm 3’* with the same GRM and sex, age and 10 PCs as covariants. The PCs were calculated based on the imputed array data for the target samples using PCAone^43^ v0.4.4. To prepare the BIMBAM format for gemma, we used *bcftools query -f ‘%CHROM:%POS, %REF, %ALT[, %MDS]\n’* to extract the dosages of the mother, and used *bcftools query -f ‘%CHROM:%POS, %REF, %ALT[,%FDS]\n’* to extract the dosages of the fetus from the VCF output of QUILT2.

We used the publicly available summary statistics of GWAS on more than 330 thousand individuals to find a list of the significant association hits in Europeans, which is downloaded from https://www.nealelab.is/uk-biobank (GWAS round 2). Then, to define the independent association signals that we need to validate using our target samples, we used the *extract_instruments* function with default parameters from the TwoSampleMR^44^ v.0.5.10 package to perform the clumping and curate a list of independent association signals for each trait.

### PRS and incremental *r*^2^

To calculate the PRS, we downloaded the polygenic score of each trait for Europeans^19^ from the PGS catalog https://www.pgscatalog.org/publication/PGP000332/. We calculated the PRS for each individual using the imputed genotype dosages and effect size of each trait, which is given by

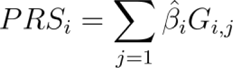

Where *PRS*_*i*_ is the PRS for each individual, 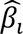 is the effect size associated with each SNP, *G_i,j_* is the imputed genotype dosage (float value between 0 and 2) for each SNP and for each individual. Since the array-imputed data used genome builds GRCh37 while the lc-WGS imputed data used GRCh38, we used the harmonized score files that shared the same variants for both genome builds, which resulted in 1,095,616 out 1,109,311 reported SNPs (98.8% matches). We used the nextflow workflow (https://github.com/PGScatalog/pgsc_calc) to automate our analyses^45^.

Then, we estimated the prediction accuracy using the incremental R2 (*IR2*), which is the difference in *r*^2^ for two regression models for each trait, one with the PRS as the predictor and one without it, which are:

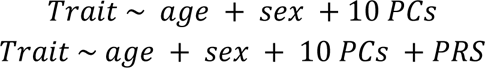

We ran the regression in R using the *lm* function. Therefore, we can obtain the *IR2* for each trait and each dataset. To estimate the accuracy of PRS using the lc-WGS imputed data compared with the array-imputed data, we defined the relative incremental R2 (*RIR2*) as follows,

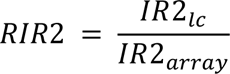

Where *IR*2_*lc*_ and *IR*2_*array*_ are the *IR2* for each trait calculated with the lc-WGS imputed data and the array-imputed data respectively. Further, to estimate the standard error (*se*) of *RIR2*, we ran the *t.test* in R on a vector of *RIR2* (21 traits).

### Benchmarking and computational cost

To benchmarking the performance of QUILT2 with respect to the size of reference panel, we used the real big reference panel, that is the UK biobank data imputed by the Genomics England haplotype reference panel, which contains N=976,630 haplotypes (488,315 individuals) and M=7,179,683 SNPs for chromosome 20. Then we randomly sampled N=2e^4^, 2e^5^, 4e^5^,6e^5^,8e^5^ haplotypes to create 6 more smaller panels. To compare QUILT2 against GLIMPSE2, we split the imputation task into multiple chunks using the same chunksize as outputted by the GLIMPSE2_chunk program. Further to fairly measure the average RAM and time for each sample, we ran both software using only one thread, although both support multithreading. To estimate the realistic cost using the WGS-derived UKB-200K panel on the RAP, we maximized the usage of CPUs threads available as possible as we can. The benchmarking was performed on virtual machines of type mem2_ssd1_v2_x4 with 4 cores and 16 Gbs memory on the RAP using normal priority. The cost was based on the instance rate card v2 (https://platform.dnanexus.com/resources/UKB_Rate_Card-Current.pdf).

## Data availability

The 1,000 Genomes Project phase 3 dataset sequenced at high coverage by the New York Genome Center is available on the European Nucleotide Archive under accession no. PRJEB31736, the International Genome Sample Resource (IGSR) data portal and the University of Michigan school of public health ftp site (ftp://share.sph.umich.edu/1000g-high-coverage/freeze9/phased/). The publicly available HRC reference panel is available from the European Genome-phenome Archive at the European Bioinformatics Institute under accession no. EGAS00001001710. The UKB-200K panel, the UKB-GEL panel, and individuals’ WGS data can be accessed via the UKB RAP (https://ukbiobank.dnanexus.com/landing). The ancient DNA from Afanasievo culture can be accessed via European Nucleotide Archive, accession number PRJEB43093, and the phased VCF file for the family are available from the European Variation Archive, accession number PRJEB46983.

## Code availability

QUILT2 is available from https://github.com/rwdavies/QUILT under a General Public License 3 (GPL3). msPBWT is available from https://github.com/rwdavies/mspbwt. vcfppR is available from https://github.com/Zilong-Li/vcfppR. Snakemake workflow for lc-WGS imputation is available from https://github.com/Zilong-Li/lcWGS-imputation-workflow.

## Acknowledgements

We thank Simon Myers for helpful discussions, and we thank Frederik Filip Stæger for assistance with downloading and preprocessing the GWAS summary statistics. We thank all participants in the UK Biobank. ZL and AA was supported by the Novo Nordisk Foundation (NNF20OC0061343). This research has been conducted using the UK Biobank Resource under Application No. 32683.

## Contributions

Z.L and R.W.D developed and implemented the method as well as conceived the study. Z.L. and R.W.D. performed the analyses. A.A. contributed to simulation, GWAS and PRS analyses for NIPT data. Z.L., A.A., and R.W.D. wrote the paper. All authors reviewed and approved the final manuscript.

## Extended online Figures

**Extended Online Figure 1.**
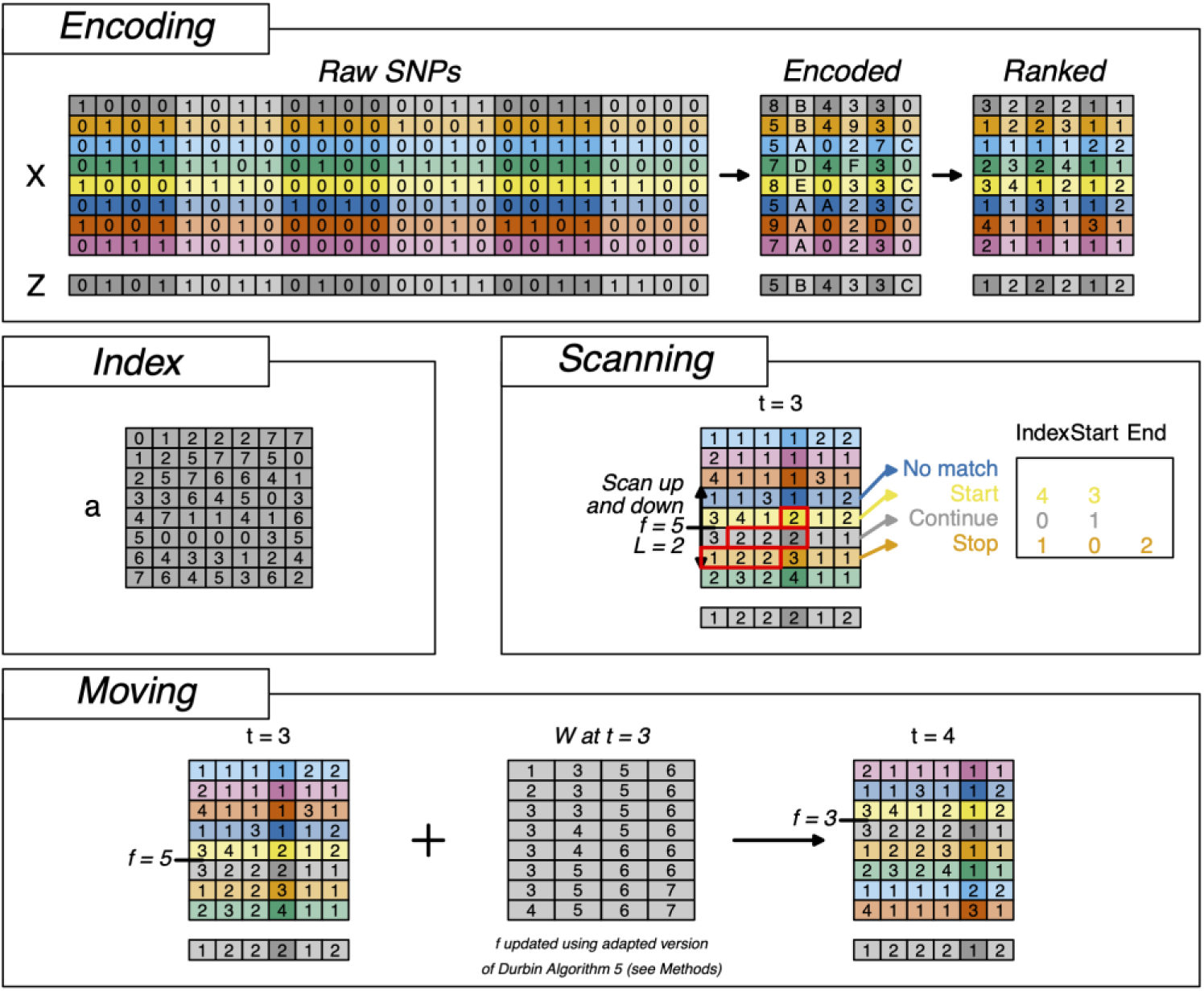
The *Encoding* panel gives an example of how a haplotype reference panel (*X*) and a target haplotype (*Z*) can be encoded and then ranked. Note that here encoding is done for 4 bits while in msPBWT it is done for 32 bits. The *Index* panel gives the index A built from the above ranked haplotype reference panel. The *Scanning* panel describes how at any grid, considering the indices that allow for the haplotype reference panel to be considered in reverse prefix sorted order, and knowing where the target haplotype would insert at that grid, allows for considering the best L matches up and down from the target haplotype. Finally the *Moving* panel shows how to use the current knowledge of where the target haplotype would insert. Here *W* is akin to FM-index for identifying where the target haplotype would match at the subsequent grid, based on the *f’=W(f, Z[t])* mapping. In practice, though shown here is the expanded form of *A* and *W*, the encoded version *U* is predominantly used for memory-efficient purposes (see Supplementary Notes).

**Extended Online Figure 2.**
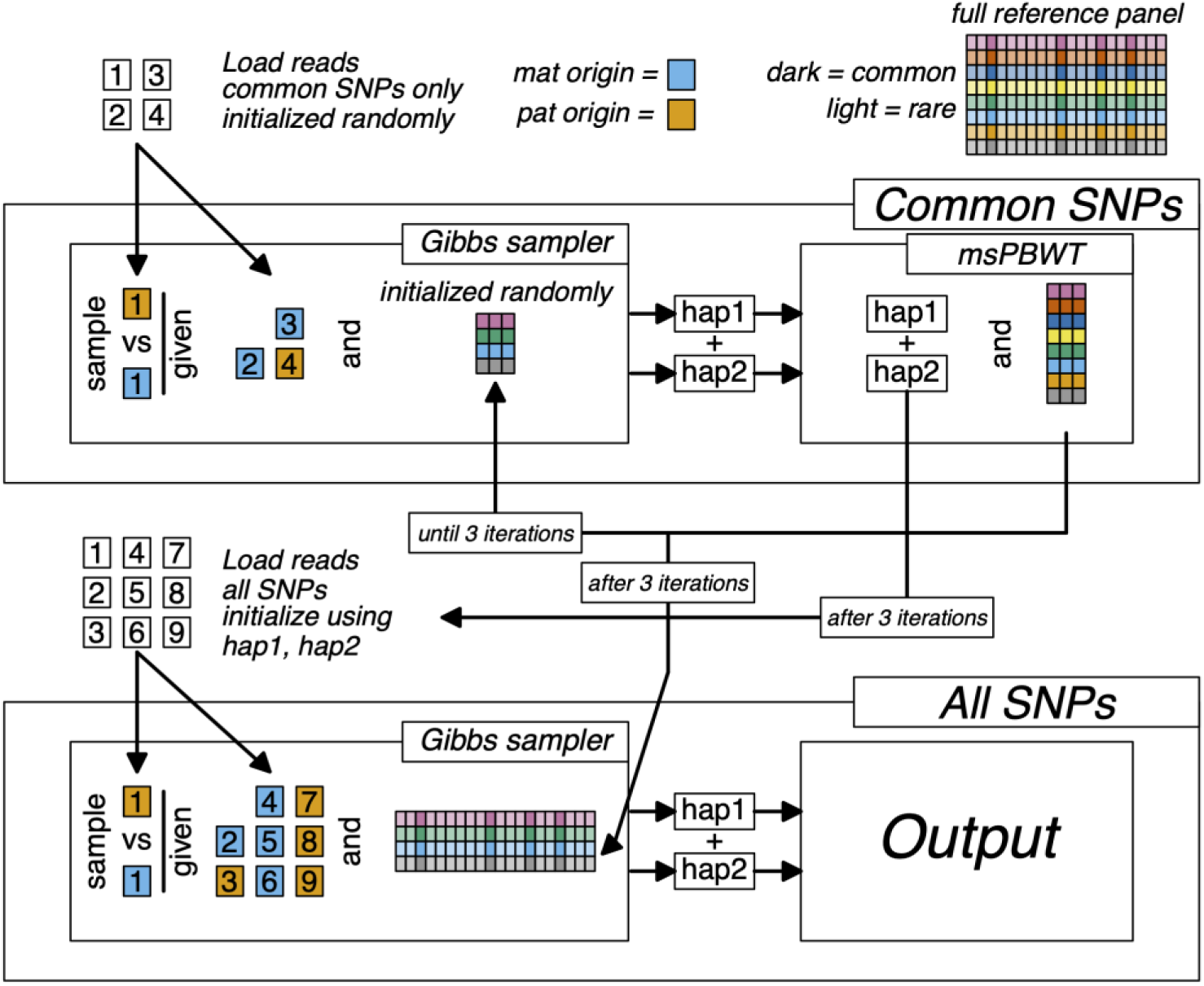
In the two step process, QUILT2 initializes by considering only common sites, by loading reads and using the haplotype reference panel at those SNPs. A two step iterative procedure then re-samples reads based on the subset of the haplotype reference panel, and updates the subset of the haplotype reference panel based on the reads and current target haplotypes from the HMM. After 3 iterations, initial haplotypes at all sites are made using the current target haplotypes at common sites and equal values otherwise, and this is used to initialize the sequencing reads, by sampling read labels based on the probability of those reads coming from the two haplotypes. Finally after Gibbs sampling of those read labels using the small haplotype reference panel at all sites, final genotype posterior probabilities and haplotypes are generated for that draw from the Gibbs sampler

**Extended Online Figure 3.**
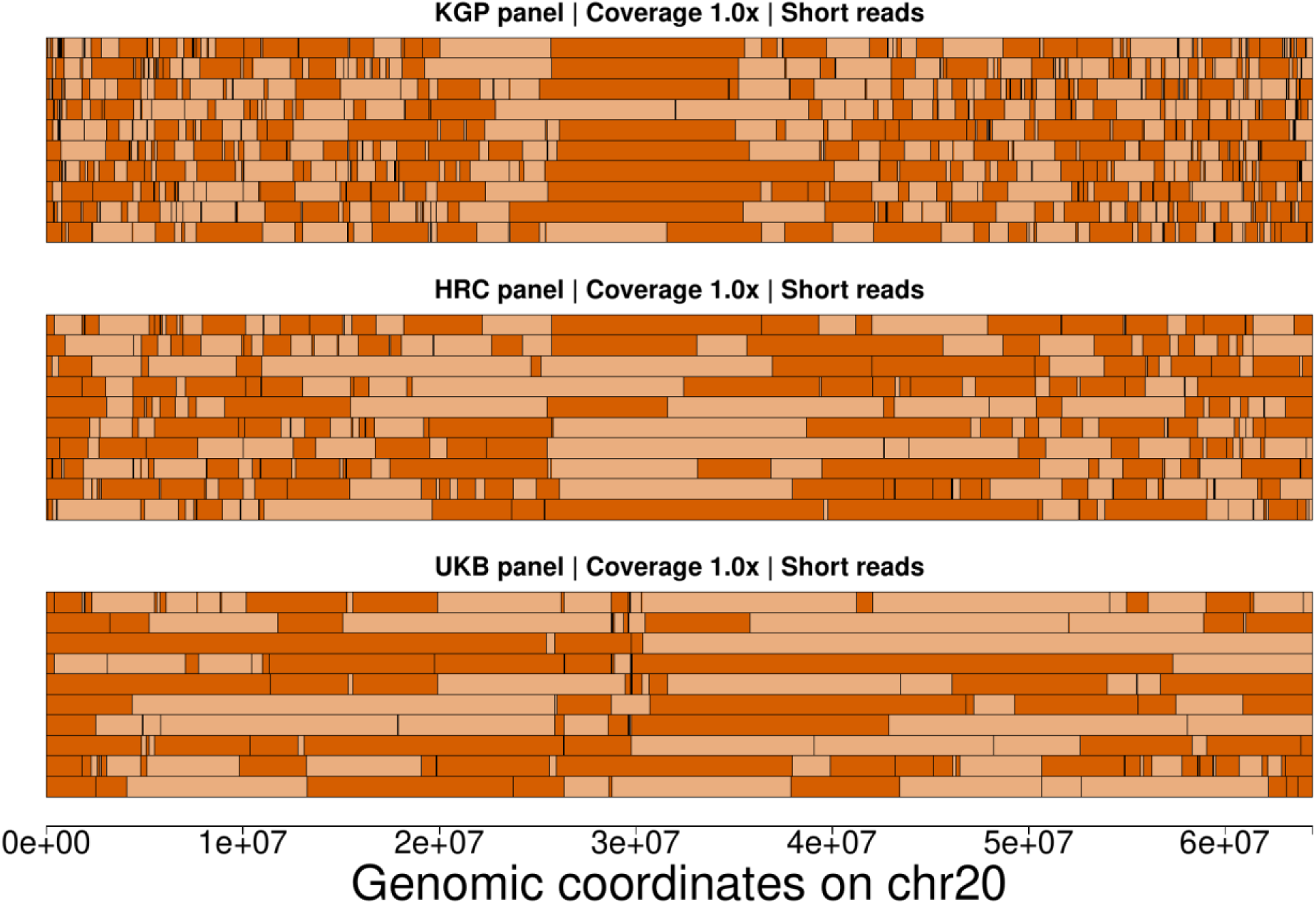
Phasing switch errors of 10 random individuals (parents) in the CEU trios at 1.0× sequencing coverage with QUILT2 using different reference panels. A switch between dark and light orange represents a phasing switch error. Results with the KGP and the HRC panel exhibited many small segments, while with the UKB panel more long segments can be seen.

