## Supplemental_Material for "Rapid and accurate genotype imputation from low coverage short read, long read, and cell free DNA sequence"

Li et al.

July 18, 2024

**2 Supplementary Figures** **14**

**3 Supplementary Tables** **24**

### 1 Supplementary Note

#### 1.1 msPBWT

##### 1.1.1 Definitions and notation

Unless otherwise indicated, assume indexing throughout is 0-based. Note this extends to when we say row or column in the text. For example, column 1 means the column indexed by 1 which is 2nd column (after the first column which is column 0). Here we will use the colon notation without any other indices to mean all entries from that row or column. For example $X_{k,:}$ means the entries from $X$ with row $k$ across all columns (again note $k$ is 0-based). We use the semi-colon notation with indices to indicate a range. For example $X_{k,a:b}$ means the entries of $X$ with row $k$ from columns $a$ through $b$ inclusive. Moreover, we use the parentheses to indicate the term is a function with parameters.

Table 1: Summary of notation related to matrices, indices and constants

| Symbol | Definition |
|---|---|
| $N$ | Number of haplotypes |
| $k$ | Choice of haplotype, $k \in \{0, ..., N-1\}$ |
| $M$ | Number of SNPs |
| $m$ | Choice of SNP, $m \in \{0, ..., M-1\}$ |
| $G$ | Number of grids, with $G = \lceil \frac{M}{B} \rceil$ |
| $g$ | Choice of grid, $g \in \{0, ..., G-1\}$ |
| $S$ | Number of unique symbols at a grid, with $S \leq 256$ |
| $s$ | Choice of symbol, $s \in \{0, ..., S-1\}$ |
| $B$ | Matrix contains binary haplotype reference panel (HRP), where each row contains a haplotype, with $N$ rows and $M$ columns, where each entry contains either a 0 or a 1, representing the reference or alternate allele |
| $R$ | Matrix with 32-bit encoded HRP, with $N$ rows and $G$ columns |
| $X$ | Matrix with encoded and re-labelled HRP, with $N$ rows and $G$ columns |
| $Y$ | $X$ in the reverse sorted prefix order (RSPO) at grid $g$ |
| $A$ | Positional prefix matrix for $X$, with $N$ rows and $G+1$ columns |
| $W$ | Matrix with symbol counts at each grid, with $N$ rows and $S$ columns (symbols), with entry $W^g_{k,s}$ being the number of instances of the symbol $s$ in the first $k$ rows of the matrix $Y$ at $g$. |
| $U$ | Encoded version of $W$, which takes only $O(N)$ space instead of $O(NS)$ |
| $z$ | Vector with encoded and re-labelled target haplotype, with $G$ entries |

##### 1.1.2 Derivation of msPBWT

Positional Burrows-Wheeler transform (PBWT) is a generic way to transform a matrix with various benefits[1]. However, the original algorithms are designed to work with binary matrix, which we refereed to as binary PBWT (b-PBWT). In contrast to b-PBWT where all indices are of $N \times M$ size and there are only 0s and 1s in the panel, here we derive the multiple symbols PBWT (msPBWT) where every e.g. 32 SNPs as a grid in the binary panel is encoded as a symbol (integer value). Therefore, all the indices arrays will have the reduced size by a factor of 32.

Here we start with the haplotype reference panel $B$, where for a single haplotype $k$ and a SNP $m$, we have $B_{k,m} \in \{0, 1\}$. Considering a particular grid $g$ in the encoded panel $R$ we have $m \in \{m_0 = 32 \times (g-1), 32 \times (g-1) + 1, ..., m_1 = \min(32 \times (g-1) + 31, M-1)\}$. We encode $B_{k,m_0:m_1}$ as a 32-bit integer and assign $R_{k,g}$ to that value using Algorithm 1.

**Algorithm 1** How R encodes 32 bit integers

**Given:** x is a binary vector of 32 length
  **if** $x_{31} = 1$ and the rest of $x$ is 0 **then**
    Return NA ▷ Integer version of NA
  **else if** $x_{31} = 1$ **then**
    Return $-1 - \sum_{i=0}^{31}(1 - x_i) \times 2^i$
  **else**
    Return $\sum_{i=0}^{31} x_i \times 2^i$
  **end if**
**Return:** a 32-bit integer

We then find the 255 most common entries of $R_{:,g}$, call then $v^g$, and re-code the entries of $R_{:,g}$ into $X_{:,g}$ using Equation 1. For any symbols in $v^g$, we rank them by frequency and recode $X_{k,g}$ to its rank in $v^g$ (0-based). We assign all symbols that do not exist in $v^g$ to 0.

$$X_{k,g} = \begin{cases} 1 + \text{argmin}_i \left(R_{k,g} = v_i^g\right) & \text{if } R_{k,g} \in v^g \\ 0 & \text{if } R_{k,g} \notin v^g \end{cases} \tag{1}$$

Now we define the transform of $X$ in the **reserve sorted prefix order** (**RSPO**). Formally, given a matrix $X = (X_{:,0}, X_{:,1}, ..., X_{:,G-1})$, we say $Y = (Y_{:,0}, Y_{:,1}, ..., Y_{:,G-1})$ that is in the reverse sorted prefix order at specific postion $g$, which means for any two rows $i < j$, with $g^* \leq g$ and $Y_{i,g^*} \neq Y_{j,g^*}$, there is $Y_{i,g^*} < Y_{j,g^*}$, as illustrated in Figure 1. This process can be seen as reordering the rows of $X$ given the values in $X_{:,g}$ and its nature order in $X_{:,g}$, which can be done with the so-called **positional prefix matrix** $A$ as the following:

$$Y = X_{A_{:,g},:} \tag{2}$$

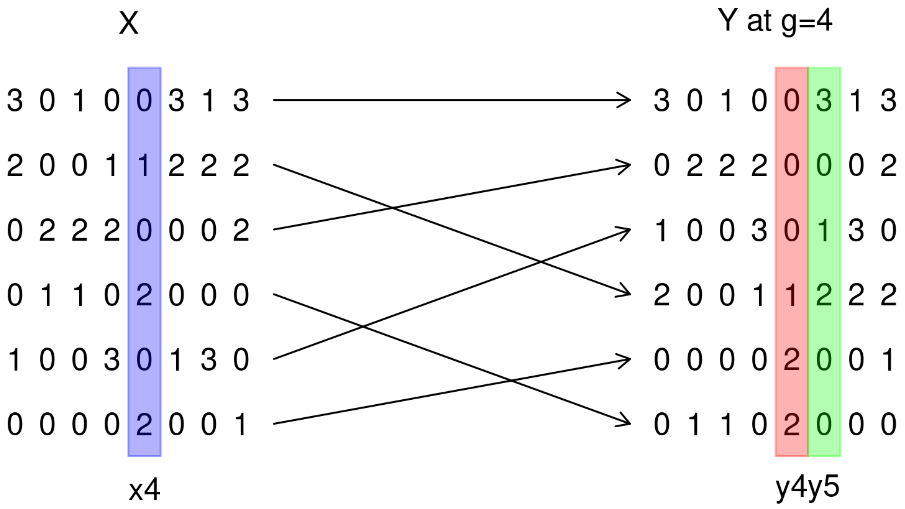

Figure 1: Example of $X$ in RSPO at $g = 4$. The entries of blue column in $X$ are re-ordered so that they are sorted in reversed prefix order before position $g = 4$ (red column) of $Y$.

We see that $A_{:,g}$ is a permutation of numbers in $\{0, 1, ..., N-1\}$ that reorders the rows of $X$ so that $Y$ is in reverse sorted prefix order at $g$. And for each postion $g$, we keep trasnforming $X$ into $Y$ so that we have such $Y$ at any postion $g$, which can help find neighbouring haplotypes at any given end point $g$. For simplicity, we will add $A_{i,-1} = i$ as the sentinal to indicate that the haplotypes in reference panel $X$ are in sequential order as they present in original source.

##### 1.1.3 Build indices for msPBWT

**Algorithm 2** Build $A_{:,g+1}$ from $A_{:,g}$

**Given:** $X$, $A_{:,g}$

$Y = X_{A_{:,g},:}$ ▷ $Y$ is the RSPO form of $X$ at $g$
$c = C(s, Y_{:,g+1})$ ▷ Number of symbols in $Y_{:,g+1}$ smaller than $s$
$u_{:} = 0$ ▷ a vector, the number of times $s$ being observed
**for** $i$ in $0 : (N-1)$ **do**
  $s = Y_{i,g+1}$ ▷ Given $Y$, get $i^{th}$ symbol at $g+1$
  $A_{c+O_s,g+1} = A_{i,g}$ ▷ Given $c$ and $O_s$, update $A_{:,g+1}$
  $u_s = u_s + 1$ ▷ Increment the number of times observing $s$
**end for**

**Return:** $A_{:,g+1}$

Now we describe how to build $A$ as we sweep through the columns of $X$, which means we build current $A_{:,g+1}$ using previous $A_{:,g}$ and current column $X_{:,g+1}$. If we know $A_{:,g}$ the sort order at position $g$, Algorithm 2 shows how to derive $A_{:,g+1}$. More specifically, given current haplotype $k = A_{i,g}$, we need to locate the position of symbol $s = Y_{i,g+1}$ in the new sorted version of $Y_{:,g+1}$ when we move to grid $g + 1$, which requires knowing *i)* the number of symbols in $Y_{:,g+1}$ smaller than $s$ and *ii)* the number of occurrences of symbol $s$ in $Y_{:,g+1}$ before $k$. These two data structures is akin to what's called FM-index[2], which is given by:

$$\begin{aligned} W_{k,s} &= C(s,x) + O(k,s,x) \\ &= \sum_{k=0}^{N-1} \mathbb{I}\{X_{k,g+1} < s\} + \sum_{i=0}^{k} \mathbb{I}\{X_{A_{i,g},g+1} = s\} \\ &= \sum_{k=0}^{N-1} \mathbb{I}\{Y_{k,g+1} < s\} + \sum_{i=0}^{k} \mathbb{I}\{Y_{i,g+1} = s\} \end{aligned} \tag{3}$$

where, $C(s, x)$ is a function that gives the number of lexically smaller characters than $s$ in $x$, and $O(i, s, x)$ gives the number of occurrences of symbol $s$ in $x$ before the position $i$. Therefore, we can locate the position of $s$ in the sorted $x$ by just looking up the table. We see that Equation 3 forms a $N \times S$ matrix with the row being haplotype index $k$ and the column being the ranked symbol $s$, which we called $W$. Note $W$ stored as described above would be very large. Here we use this definition of $W$ in what follows as it is easy for understanding the algorithm. In practice we use an encoded version of $W$ described in Section 1.1.5. With $W$ stored in memory, we can have the bijection mapping from $A_{:,g+1}$ to $A_{:,g}$ , as illustrated in Figure 2, which is given by

$$A_{W_{i,Y_{i,g+1}}-1,g+1} = A_{i,g} \tag{4}$$

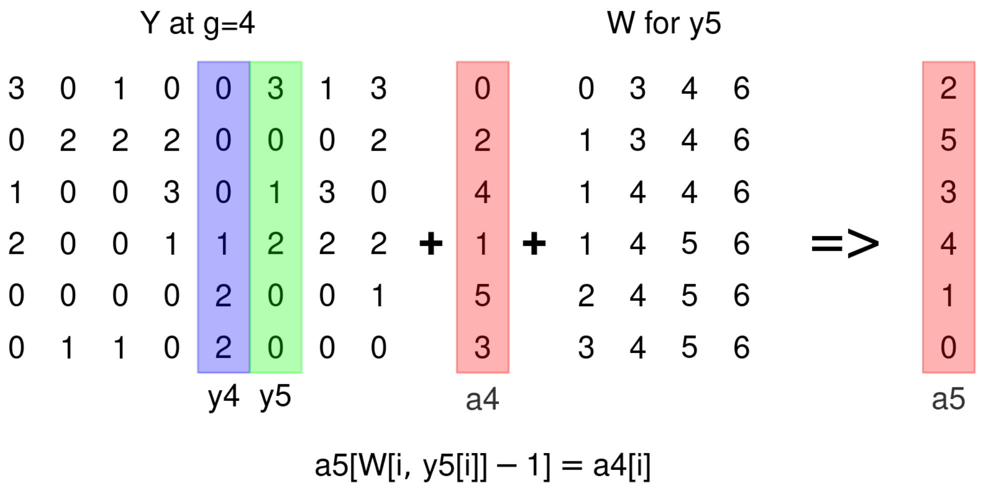

Figure 2: Illustration of deriving $A_{:,g}$ from $A_{:,g+1}$ with FM-index $W$. For example, let $i = 1$, we have $y5[i] = 0$ and $W[i, y5[i]] - 1 = W[1, 0] - 1 = 0$. Thus, $a4[1] = a5[0] = 2$.

##### 1.1.4 Haplotype matching with msPBWT

In the contexts of compression and matching, we only consider $Y$ instead of $X$ because $Y$ is ordered while $X$ is not. Furthermore, $Y$ is expected to be strongly run-length compressible due to linkage disequilibrium (LD). Here we focus on the case that given a new haplotype $z$, and we want to find the haplotypes in $Y$ that share long segments ending at each grid $g$. As proposed in b-PBWT[1] and Syllable-PBWT[3], one can search for set-maximal matches using additional data structure, such as the divergence array or rolling hashing. In contrast, for memory reasons, we use the idea of scanning up and down only $L$ haplotypes at each grid.

Here we define the vector $f$ of length $G$, where at each grid $g$ there is $z_g = Y_{f_g,g}$. This is useful because if we know $f_g$, then we can easily find the nearest neighbors in $Y$ to $z$ at grid $g$. We can build $f_{g+1}$ from $f_g$ easily by using FM-index structure as noting in Equation 4, which is detailed in Algorithm 3. Once we have $f$, as illustrated in Figure 3, we can easily insert $z$ back into $Y$ at each grid $g$ and find haplotype matches by scanning up and down.

**Algorithm 3** Determine $f$ at each column

**Given:** $z$, $X$, $W$

- $Y = X_{A_{:,0},:}$ ▷ $Y$ is the RSPO form of $X$ at $g = 0$
- $f_0 = \text{argmax}_i(Y_{i,0} = z_0)$ ▷ Initial can be any $i$ such that $Y_{i,0} = z_0$
- **for** $g$ in $1 : (G - 1)$ **do**
  - $i = \min(f_{g-1}, N - 1)$ ▷ Protect boundary
  - $s = z_g$ ▷ Current symbol
  - $f_g = W_{i,s}$ ▷ $W$ is decoded from $U$
- **end for**

**Return:** $f$

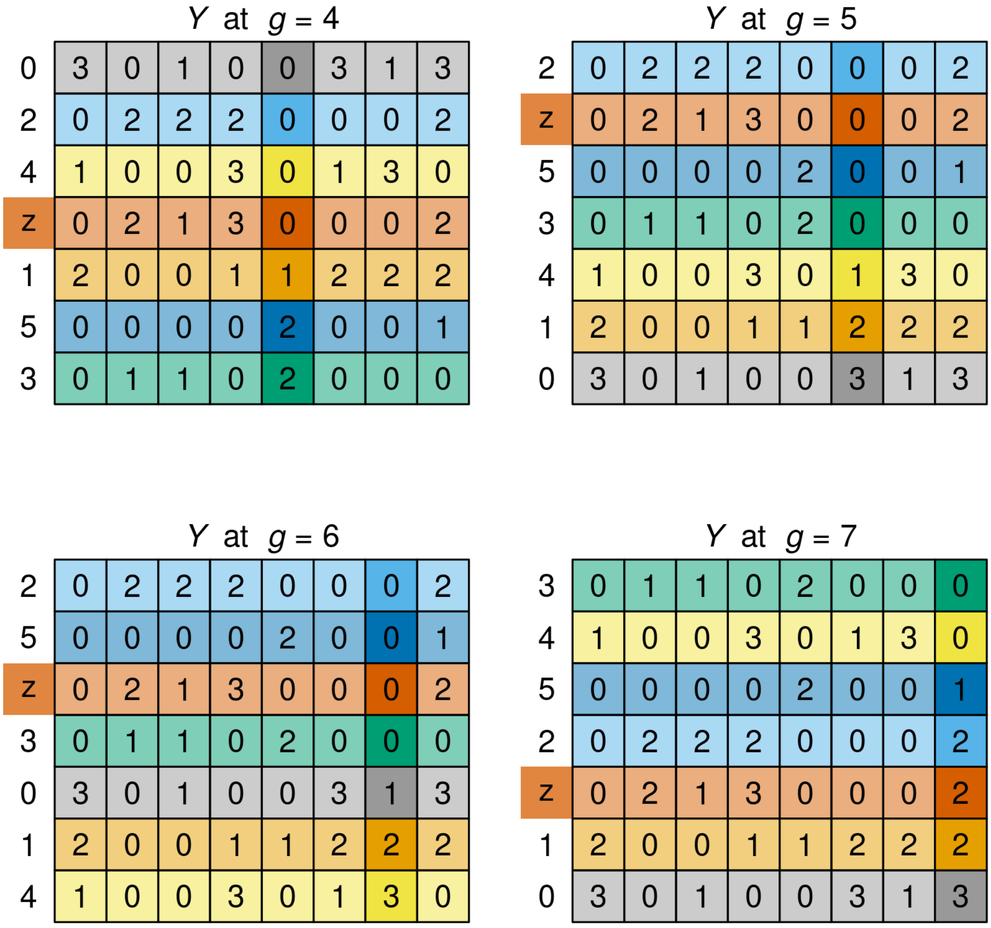

Figure 3: Illustration of inserting $z$ back into $Y$ given $f = (3, 6, 5, 6, 3, 1, 2, 4)$. We can see that haplotype 2 and 5 are the neighbors of $z$ at grid 5,6 and 7.

Now given $f$ at every grid and $L$ number of neighboring haplotypes to be scanned (up and down), we perform Algorithm 4 to find a set of matches from $Y$ to $z$ with a user-defined minimum length. Note here we only present the algorithm for scanning up and the same principle applies to the scanning down. Throughout this section, we assume we have $A$ stored at all grids. However, for being memory-efficient, we can actually store a subset of the columns of $A$ and then use $W$ to rebuild other entries in $A$ as needed using Equation 4.

Now, on to the algorithm. Determining the indices of the closest $L$ matches from $Y$ to $z$ at $g$ is straightforward, if we look them up using $A_{:,g}$, with the $l^{th}$ closest match up being $A_{f_g-l,g}$. However, this doesn't give us the start and end of the matches, or the length. Instead of calculating these for each grid, we keep track of matches by walking backwards through $f$, and noting which of the $L$ indices at each grid change, and only once an index is lost, indicating a match ends. Then we can record the matching haplotype, its start and end. This is detailed in Algorithm 4.

More specifically, suppose for some $g+1$ we consider the matrix $p$ and we set $p_{l,g+1} = f_{g+1} - l$, as well as $p_{:,g}$. Consider briefly the reverse, that is if we consider the top $L$ matches of $z$ to $X$ for some $g$, and consider moving forward one grid, then for some $l^*$ at $g$ and some potential $l$ at $g+1$: the match can be extended, in which case $\exists l^* \leq L$ and $A_{p_{l^*,g},g} = A_{p_{l,g+1},g+1}$ (and we must have $l^* \geq l$); or, the match ends, and no such match exists. As such, we can do the reverse, and scan up through $l$ at $g+1$ to see if $l^*$ exists, and if not, we record that the match has ended.

However, while this gives us an end point (the match must end at $g$), this process doesn't give a start point. The matches can enter into the top $L$ at $p_{:,g}$ either because they start matching, or, more likely, because other matches that started earlier were lost. As such, by keeping track of $p_{:,g}$ and $p_{:,g+1}$, we know when a match ends, a minimal value at which it must start, and can then work out the haplotype index, its start, and hence the length. As such, throughout this process, we can find any match of at least certain length that is one of the $L$ closest matches of $X$ to $z$ at any point.

**Algorithm 4** Find long matches from $X$ to $z$ by scanning up $L$ neighbors

**Given:** $z$, $f$, $A$, $X$

$g = G - 1$ ▷ Last grid, walking backwards
**for** $l$ in $0 : (L-1)$ **do**
  $v = f_g - l$, infer $k$ such that $A_{k,g-1} = A_{v,g}$ ▷ Use FM-index $W$
  $p_{l,g} = k$ ▷ Keep track of the neighboring haplotypes
**end for**
**for** $g$ in $(G-2) : 0$ **do**
  $c = 0$ ▷ Counter
  **for** $l$ in $0 : (L-1)$ **do**
    $v = f_g - l$, infer $k$ such that $A_{k,g-1} = A_{v,g}$
    $s = X_{A_{v,g},g}$ ▷ Current symbol
    **if** $z_g == s$ and $p_{c,g+1} == v$ **then**
      $c = c + 1$
    **end if**
    $p_{l,g} = k$ ▷ Store $L$ up neighbours at $g-1$
  **end for**
  **while** $c < L - 1$ **do**
    $c = c + 1$
    $e = g$ ▷ End of matching
    $i = A_{p_{c,g+1},e}$ ▷ Index of haplotype
    $t = \text{argmin}_{g^* < e}(X_{i,g^*} \neq z_{g^*})$ ▷ Start of matching
    Report $i, t, e$
  **end while**
**end for**

**Return:** a set of haplotype matches with the starts and ends

##### 1.1.5 Encode and decode msPBWT data structure

In the above we assume $W$ is stored for each grid, which requires $O(GNS)$ space for the entire panel. Given $S = 256$ in the limit at each grid in a big reference panel, the total space would be up to $O(256GN)$, which is very inefficient. However, $W$ is highly structured (see Figure 2) and hence we can store $W$ much efficiently. Here we describe how to encode $W$ into $U$ using as few as $O(GN)$ space while having fast access to any value in $W$.

First, we consider encoding $W$ at each grid separately. We further choose to compress each column (each symbol, $W_{:,s}$) independently. Intuitively, for some symbols, call these minimal symbols, we will observe very few instances, and it is easy to store this as a vector of those increases. For example, if $W_{:,s}$ is a vector of length 100 (so $N = 100$) with entries in positions 1 through 21 of 0, 22 through 51 of 1, and 52 through 101 as 2, we can store a vector $v^s = (21, 51)$, and return $W_{k,s} = \sum_{i=1}^{|v^s|} \mathbb{I}\{v_i^s < k\}$. Note this operationally is implemented as a while loop as it is faster.

Alternatively, for other symbols, we observe very many instances - call these maximal instances - however, most of those tend to be clustered, or in other words, we see long runs that increase by

1, and long runs that are constant. This makes sense due to local LD between grids, as there is high correlation between symbols at neighboring grids. As the above storage is inefficient in such a case, we consider the following.

We define an integer, call it $N_e$ (100 in default), and seek to store the true value of $W_{:,s}$ every $N_e$ entries. Intuitively, we can split $W_{:s}$ into multiple bins, and encode each bin by storing the alternating lengths of runs that increase by one. For example, $0, 1, 2, 2, 2, 2, 3$ would be stored as 2,3,1 as there are two +1 increments, three +0 increments, and one +1 increments. We then ultimately store $W_{:,s}$ for maximal columns in a two dimensional array $U^b_{rle}$, with the first being a bin and the second being the run length encoding of that bin as well as the start and end of the bin. Now we can efficiently retrieve any value in $W_{k,s}$ from $U$ as follows. First, we need to locate the bin $W_{k,s}$ falls into by $b = \lfloor \frac{k}{N_e} \rfloor$. Then let $o = k - bN_e$ be the offset, we can decode the run-length encoding $U^b_{rle}$ given $o$. As we would expect long runs that increase by 0 or 1 in each bin, the decoding procedure has $O(1)$ time in most cases with the worst case being $O(N_e)$.

As such, we can encode and decode $U$ efficiently. We could further encode $U$ to use less space for instance gzipping the resulting objects above, however given it takes similar space to $A$, it is of marginal benefit.

##### 1.1.6 Comparison of PBWT variants

We note that there is Syllable-PBWT[3] published at the same time, which also aims to be memory-efficient. Both Syllable-PBWT and msPBWT adopt the same idea of encoding the binary haplotype into integer symbols first, while they differs in the data structure and algorithm designed for querying a new haplotyp $z$ to the reference panel $X$. Here we summarize the space usage of data structures for each PBWT variant in Table 2, which shows that msPBWT uses 2× less space than Syllable-PBWT in storage. However, we note that the haplotype matching algorithm of msPBWT does not guarantee finding the set-maximal matches, and the set-maximal matches may not be the optimal ones for imputation[4]. In our tests, we found the Algorithm 4 of msPBWT can find long-enough matches locally that are informative in genotype imputation.

Table 2: Space comparisons of PBWT variants in haplotype matching. Values are in units of $N \times M$ 32-bit integers. Note $p \geq 1, \alpha \geq 1$ and $B \in \{32, 64, 128\}$.

| Space usage | Bit-PBWT | Syllable-PBWT | msPBWT |
|---|---|---|---|
| Panel of sequences (X) | $\frac{1}{32}$ | $\frac{1}{B}$ | $\frac{1}{B}$ |
| Positional prefix arrays (A) | 1 | $\frac{1}{B}$ | $\frac{1}{\alpha B}$ |
| Syllable dictionaries | - | $\frac{1}{32p}$ | - |
| FM-index arrays (U) | 2 | - | $\frac{1}{B}$ |
| Prefix hash arrays | - | $\frac{2}{B}$ | - |
| Divergence matrix | 1 | - | - |
| Total space | $4 + \frac{1}{32}$ | $\frac{4}{B} + \frac{1}{32p}$ | $\frac{2}{B} + \frac{1}{\alpha B}$ |

W with 40 haplotypes and 6 symbols

| | | | | | |
|---|---|---|---|---|---|
| 0 | 1 | 2 | 4 | 6 | 12 |
| 0 | 1 | 2 | 4 | 6 | 13 |
| 0 | 1 | 2 | 4 | 6 | 14 |
| 0 | 1 | 2 | 5 | 6 | 14 |
| 0 | 1 | 2 | 5 | 6 | 15 |
| 0 | 1 | 2 | 5 | 6 | 16 |
| 0 | 1 | 2 | 5 | 6 | 17 |
| 0 | 1 | 2 | 5 | 6 | 18 |
| 0 | 1 | 2 | 5 | 6 | 19 |
| 0 | 1 | 2 | 5 | 7 | 19 |
| 0 | 1 | 2 | 5 | 7 | 20 |
| 0 | 1 | 2 | 5 | 7 | 21 |
| 0 | 1 | 2 | 5 | 8 | 21 |
| 0 | 1 | 3 | 5 | 8 | 21 |
| 0 | 1 | 3 | 5 | 8 | 22 |
| 0 | 1 | 3 | 5 | 8 | 23 |
| 0 | 2 | 3 | 5 | 8 | 23 |
| 0 | 2 | 3 | 5 | 9 | 23 |
| 0 | 2 | 3 | 5 | 9 | 24 |
| 0 | 2 | 3 | 5 | 9 | 25 |
| 0 | 2 | 3 | 5 | 9 | 26 |
| 0 | 2 | 3 | 5 | 9 | 27 |
| 0 | 2 | 3 | 5 | 9 | 28 |
| 0 | 2 | 3 | 5 | 9 | 29 |
| 0 | 2 | 3 | 5 | 9 | 30 |
| 0 | 2 | 3 | 5 | 9 | 31 |
| 1 | 2 | 3 | 5 | 9 | 31 |
| 1 | 2 | 3 | 5 | 9 | 32 |
| 1 | 2 | 3 | 6 | 9 | 32 |
| 1 | 2 | 3 | 6 | 10 | 32 |
| 1 | 2 | 3 | 6 | 10 | 33 |
| 1 | 2 | 3 | 6 | 10 | 34 |
| 1 | 2 | 3 | 6 | 10 | 35 |
| 1 | 2 | 3 | 6 | 10 | 36 |
| 1 | 2 | 3 | 6 | 10 | 37 |
| 1 | 2 | 4 | 6 | 10 | 37 |
| 1 | 2 | 4 | 6 | 10 | 38 |
| 1 | 2 | 4 | 6 | 11 | 38 |
| 1 | 2 | 4 | 6 | 11 | 39 |
| 1 | 2 | 4 | 6 | 11 | 40 |

Binning the red column of W per 5 rows

| | | | | |
|---|---|---|---|---|
| 12 | 13 | 14 | 14 | 15 |
| 16 | 17 | 18 | 19 | 19 |
| 20 | 21 | 21 | 21 | 22 |
| 23 | 23 | 23 | 24 | 25 |
| 26 | 27 | 28 | 29 | 30 |
| 31 | 31 | 32 | 32 | 32 |
| 33 | 34 | 35 | 36 | 37 |
| 37 | 38 | 38 | 39 | 40 |

Run length encoding of each bin above

| | | | | |
|---|---|---|---|---|
| 12 | 2 | 1 | 1 | |
| 16 | 3 | 1 | | |
| 20 | 1 | 2 | 1 | |
| 23 | 0 | 2 | 2 | |
| 26 | | | | |
| 31 | 0 | 1 | 1 | 2 |
| 33 | 4 | | | |
| 37 | 1 | 1 | 2 | |

Figure 4: Illustration of encoding $W$ into $U$. The first symbol (blue column) uses the mininal encoding ($U = (0, 26)$), while the 6-th symbol (red column) uses the maximal encoding. For maximal encoding, we first split the column into bins per $N_e = 5$ rows and store the run length encoding of each bin and some references to the start position. Note we only store the start of the 5-th bin (yellow row) because it has sequential values from 26 to 30. We expect long runs that increase by 0 or 1 in a real haplotype reference panel.

##### 1.2 NIPT imputation

Table 3: Summary of notation related to counts, observed data and haplotype labels

| Symbol | Definition |
|---|---|
| $N_r$ | Number of reads in total. |
| $R_r$ | Read with index $r$ which spans $j_r$ SNPs, with SNP indices $u_r$, sequenced bases $s_r$ and base qualities $b_r$ or $R_r = \{u_r, s_r, b_r\}$. |
| $J_r$ | Number of SNPs spanned by read $R_r$ |
| $c_r$ | Central SNP for read $R_r$ |
| $O_t$ | Set of reads with central SNP $t$, $O_t = \{R_r \| c_r = t\}$ |
| $O$ | Set of observations for each SNP $t$ on the chromosome, $O = \{O_t \| t = 1, ..., T\}$ |
| $O_t^i$ | Set of reads from haplotype $i \in \{1, 2, 3\}$, $O_t^i = \{R_r \| c_r = t, H_r = i\}$ |
| $O^i$ | Observations for each SNP on the chromosome from haplotype $i$, $O^i = \{O_t^i \| t = 1, ..., T\}$ |
| $u_{r,j}$ | For SNP $j$ in read $R_r$, its index with respect to the chromosomal listing of SNPs (*e.g.* If the physical position of SNP $t$ in the region is $L_t$ for $t = 1, ..., T$, then SNP $j$ in read $R_r$ has physical position $L_{u_{r,j}}$) |
| $s_{r,j}$ | Sequencing base for SNP $j$ in read $R_r$, with $s_{r,j} = 1$ for the alternate base and 0 for the reference base |
| $b_{r,j}$ | Base quality for SNP $j$ in read $R_r$ |
| $R_{r,j}$ | Subset of read $R_r$ for SNP $j$, or $R_{r,j} = \{u_{r,j}, s_{r,j}, b_{r,j}\}$ |
| $\phi_{r,j}^i$ | Probability of SNP $j$ from read $R_r$ coming from an underlying genotype $i$, or $P(s_{r,j} \| g = i)$ |
| $H$ | Vector of haplotype membership labels |
| $H_r$ | Variable that takes value 1 or 2 if read $R_r$ comes from the maternal haplotypes and 3 if it comes from the paternal transmitted haplotype in the fetus |
| $H_{-r}$ | Vector $H$ with member $r$ removed |
| $\mathcal{H}$ | Space on which $H$ is defined. Hamming cube with entries 1 and 2 (versus usual 0 and 1) for diploid, and 1, 2, 3 for triploid |

###### 1.2.1 Gibbs sampling setup

For some arbitrary SNP indexed by $t$ that we are interested in estimating, let $Hap$ be the haploid genotype, with arbitrary labels $I \in \{1, 2, 3\}$ indicating the maternal transmitted, the maternal untransmitted and the paternal transmitted, respectively. Consider $Gen$ as the diploid genotype for the mother or fetus, and that we want the posterior genotype probabilities $P(Gen|O, \lambda)$, which can be used to give us the diploid genotype dosage in the normal way as

$$E(Gen|O, \lambda, FF) = \sum_{g=0}^{2} g \times P(Gen = g|O, \lambda, FF) \tag{5}$$

We can calculate $P(Gen = g|O, \lambda, FF)$ using Monte Carlo as follows (with $\approx$ becoming $=$ in the limit)

$$\begin{aligned} P(Gen = g|O, \lambda, FF) &= \sum_{H \in \mathcal{H}} P(Gen = g|H, O, \lambda) P(H|O, \lambda, FF) \\ &\approx \sum_{H \sim P(H|O,\lambda,FF)} P(Gen = g|H, O, \lambda) \end{aligned} \tag{6}$$

We can rapidly calculate $P(Gen = g|H, O, \lambda)$, e.g. for the fetus under the model as

$$P(Gen = g|H, O, \lambda) = \begin{cases} P(Hap^1 = 0|H, O, \lambda, FF) \times P(Hap^3 = 0|H, O, \lambda, FF) & \text{if g = 0} \\ P(Hap^1 = 0|H, O, \lambda, FF) \times P(Hap^3 = 1|H, O, \lambda, FF)+ & \\ P(Hap^1 = 1|H, O, \lambda, FF) \times P(Hap^3 = 0|H, O, \lambda, FF) & \text{if g = 1} \\ P(Hap^3 = 1|H, O, \lambda, FF) \times P(Hap^3 = 1|H, O, \lambda, FF) & \text{if g = 2} \end{cases} \quad (7)$$

and further through the HMM model in imputation process, we can rapidly calculate $P(Hap^i = 1|H, O, \lambda, FF)$ for some SNP $t$ as

$$P(Hap^i = 1|H, O, \lambda, FF) = \sum_{k=1}^{K} \theta_{t,k} P(Q^i_{gt} = k|O, \lambda, FF) \quad (8)$$

We use the same notation as in the QUILT paper, and have that $\theta_{t,k}$ gives the probability that reference haplotype $k$ carries the alternate allele at SNP $t$, and $P(Q^i_{gt} = k|O, \lambda, FF)$ is the probability the the sample copies from haplotype $k$ at grid point $g_t$.

We therefore need rapid draws of the read labels conditional on the observed data, $H \sim P(H|O, \lambda, FF)$, which can be done using Gibbs sampling in way described in the following subsection.

##### 1.2.2 Gibbs sampling practical

Now, for $H$ a vector random variable of read labels, with realization $h$, suppose in the Gibbs sampling, for some initial $H$, we start with $h$, initialized at random, given the fetal fraction $FF$.

That is

$$P(H_v = h_v|FF) = \begin{cases} 0.5, & h_v = 1 \\ 0.5 - \frac{FF}{2}, & h_v = 2 \\ \frac{FF}{2}, & h_v = 3 \end{cases} \quad (9)$$

where arbitrary 1 = maternal transmitted, 2 = maternal untransmitted, 3 = paternal transmitted.

Now, let $h_v$ be the current label for read $v$ and $h_v^{o1}$, $h_v^{o2}$ be the alternate read labels, i.e $h_v + h_v^{o1} + h_v^{o2} = 6$ (for example, if $h_v = 2$, then $h_v^{o1} = 1$, $h_v^{o2} = 3$). With Gibbs sampling, we want to efficiently calculate

$$P(H_v = i|H_{-v}, O, \lambda, FF) = \frac{P(O, H_v = i, H_{-v} = h_{-v}|\lambda, FF)}{\sum_{j=1}^{3} P(O, H_v = j, H_{-v} = h_{-v}|\lambda, FF)} \quad (10)$$

where $i \in \{1, 2, 3\}$. This can be done if we can efficiently calculate $P(O, H_v = i, H_{-v} = h_{-v}|\lambda, FF)$, noting that we already have $P(O, H_v = h_v, H_{-v} = h_{-v}|\lambda, FF)$ under the HMM. We can write down the NIPT version of the joint probability as

$$P(O, H_v = i, H_{-v} = h_{-v}|\lambda, FF) = P(O^1|\lambda)P(O^2|\lambda)P(O^3|\lambda) \prod_{N_r} P(H = h|FF) \quad (11)$$

Where here $P(O^i|\lambda)$ is the probability of the reads belonging to haplotype $i$ (including read indexed by $v$). The rapid calculation of these probabilities is not exhaustively detailed here but follows in the obvious way from what was described in the supplementary of QUILT paper (`https://www.nature.com/articles/s41588-021-00877-0#Sec26`). Note, however, as compared to the QUILT model, for the NIPT model, as opposed to the diploid model, we need to take the probability of read labels (9) into account, as these do not cancel.

##### 1.2.3 Gibbs sampling heuristics

Heuristics are essential to ensure that read label vectors do not get stuck in local extrema. The heuristics in QUILT2 for NIPT are broadly similar to QUILT; however, they must become substantially more complicated, to deal with the unequal prior probabilities based on the different frequencies of reads coming from the different underlying haplotypes, as will be explained below.

Recall that the standard Gibbs sampler works through

$$P(H_v = h_v | H_{-v} = h_{-v}, O = o, \lambda, FF) = \frac{P(O = o, H_v = h_v, H_{-v} = h_{-v} | \lambda, FF)}{\sum_{j=1}^{3} P(O = o, H_v = j, H_{-v} = h_{-v} | \lambda, FF)} \tag{12}$$

Suppose we have identified a partition of the region to be imputed, such that we define blocks based on grid points, such that each SNP, each grid, and each read, uniquely falls into this partioning. Without loss of generality, consider one such block. Here we will use 1-based indices. Let $t_1$ and $t_2$ be the SNP starts and ends of that block, $g_1$ and $g_2$ be the grid start and end, and let $r_1$ and $r_2$ be the first and last read in that region. Let $V = [r_1, r_1 + 1, ..., r_2]$ be an increasing vector of the read indices. Then, given the current vector of labels $h_V$, we want to efficiently sample, for some function $f$, $f(h_V, i)$, the following

$$P(H_V = f(h_V, i) | H_{-V} = h_{-V}, O = o, \lambda, FF) = \frac{P(O = o, H_v = f(h_V, i), H_{-V} = h_{-V} | \lambda, FF)}{\sum_{j=1}^{3} P(O = o, H_V = f(h_V, j), H_{-V} = h_{-V} | \lambda, FF)} \tag{13}$$

Now, consider the matrix $rr$ , defined as

$$rr = \begin{bmatrix} 1 & 2 & 3 \\ 1 & 3 & 2 \\ 2 & 1 & 3 \\ 2 & 3 & 1 \\ 3 & 1 & 2 \\ 3 & 2 & 1 \end{bmatrix} \tag{14}$$

Consider $f(h_V, i)$. In the simplest form, we want a block Gibbs heuristics to consider switching entire label groups, e.g,

$$f(h_r, i) = rr_{i,h_r} \tag{15}$$

Note that we are using notation that assumes that we can let $f$ act on a vector in the obvious way i.e $[f(h_V, i)]_j = rr_{i,h_{V_j}}$. However, this definition of $f$ is not ideal, as it will make it difficult for the majority of proposed block Gibbs swaps to occur, as reads that are equally likely to come from sets of haplotypes will assort to those haplotypes with frequencies proportional to their relative prior probabilities based on the fetal fraction, and then trying to flip based on those probabilities will be unlikely. For example, suppose a region of the maternal genome was homozygous, and the maternal transmitted, and untransmitted haplotypes, were identical. We would want a block Gibbs sampler in this region to choose to swap the haplotypes with equal probability. However, as the reads that map to those two haplotypes (but not the third) will be distributed not at random, but proportional to their prior probabilities of 0.5 and $0.5 - \frac{FF}{2}$, as such, such a swap would be less likely than remaining stationary.

To avoid this, we consider a version of $f$ that re-samples read labels according to a class based on which sets of haplotypes they could have come from. Let $p_1 = 0.5$, $p_2 = \frac{1-FF}{2}$, $p_3 = \frac{FF}{2}$. Then consider the matrix $rlc$ as below

$$rlc = \begin{bmatrix} 1 & 0 & 0 \\ 0 & 1 & 0 \\ 0 & 0 & 1 \\ \frac{p_1}{p_1+p_2} & \frac{p_2}{p_1+p_2} & 0 \\ \frac{p_1}{p_1+p_3} & 0 & \frac{p_3}{p_1+p_3} \\ 0 & \frac{p_2}{p_2+p_3} & \frac{p_3}{p_2+p_3} \\ p_1 & p_2 & p_3 \end{bmatrix} \tag{16}$$

Let's suppose then we have, from the previous iteration, $P(H_r = i|O, \lambda, FF) \forall i \in \{1, 2, 3\}$. Define a read class as the set of haplotypes that read likely came from. More formally, let the read class of read $r$ be $H_r^* \in \{0, ..., 7\}$ where 0 is no match and 1 through 7 represent matching to one of the rows above suitably close. Specifically, consider $d(j) = \sum_{i=1}^{3} |rlc_{j,i} - P(H_r = i|O, \lambda, FF)|$, and some threshold $y$. Then

$$H_r^* = \begin{cases} \text{argmin}_{j \in \{1,...,7\}} d(j), & \min_{j \in \{1,...,7\}} d(j) < y \\ 0, & \text{otherwise} \end{cases} \tag{17}$$

Therefore, for sampling, we care about the read class $H_r^*$, and not just the read label $H_r$, and we define $f(h_r, i)$ instead as $f(h_r^*, i)$ as sampling from a multinomial probability with

$$P(f(h_r^*, i) = j) = \begin{cases} rlc_{h_r^*, rr_{i,j}}, & h_r^* \neq 0 \\ rr_{i,j}, & h_r^* = 0 \end{cases} \tag{18}$$

Therefore, we are not stricly performing a Gibbs sampling, though in practice, this substantially improves the quality of the Gibbs sampling, and the accuracy of imputation.

Alternatively, one can consider a two stage approach, which is equivalent, but easier to implement computationally. First, for a given $ir$ (*i.e.* row of $rr$, corresponding to what swap to perform), the read class memberships are re-sampled, and then second, read labels are re-sampled based on this according to $rr$.

#### 2 Supplementary Figures

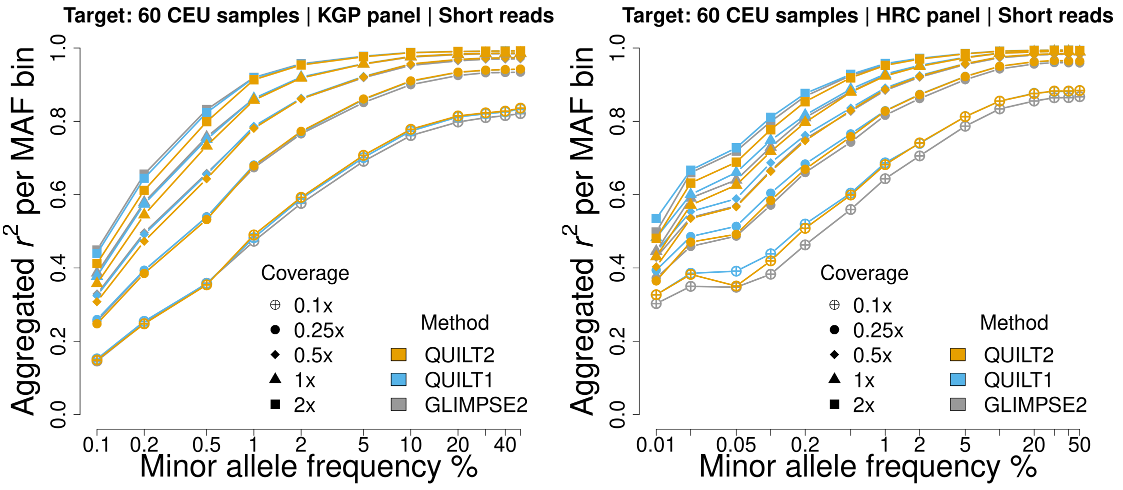

Supplementary Figure 1: Compare imputation accuracy of QUILT2 against QUILT1 and GLIMPSE2 for small reference panels, which are the HRC and KGP panel. QUILT1 is the most accurate and robust across all tests, while QUILT2 only has slightly reduced accuracy using small reference panels for rare variants at higher coverage compared to GLIMPSE2.

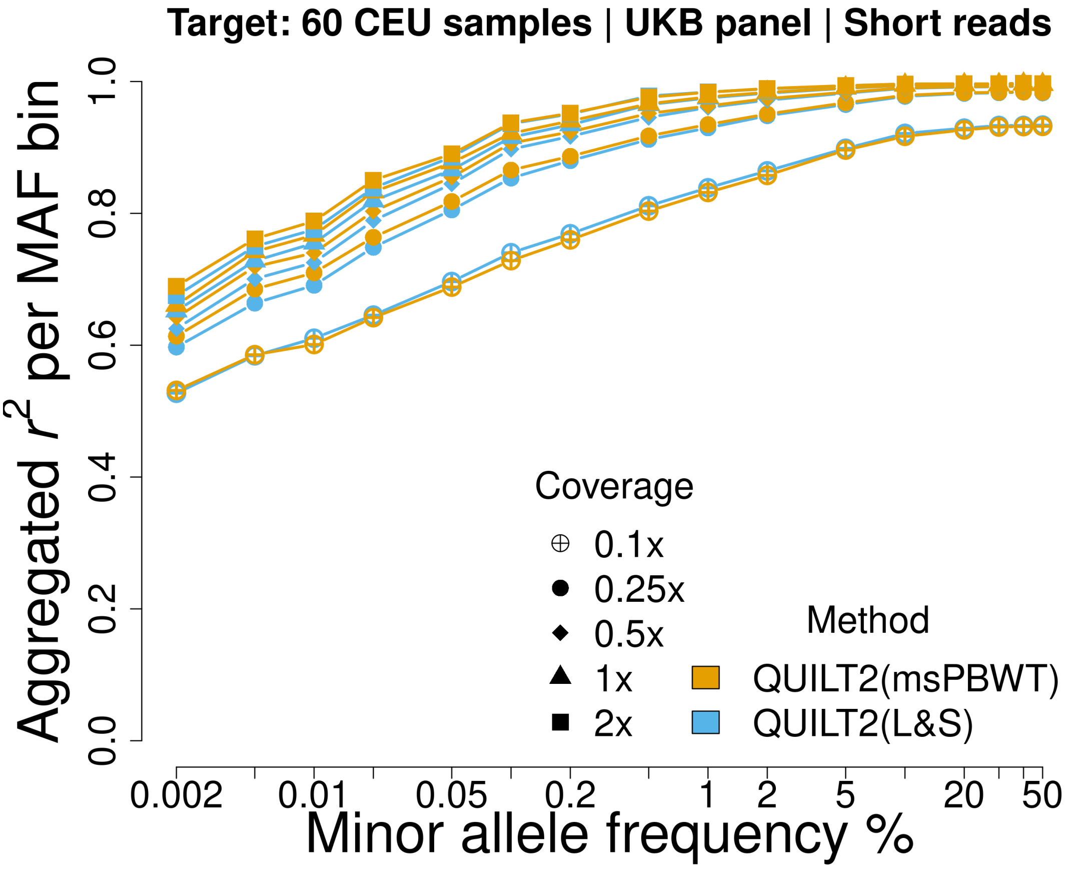

Supplementary Figure 2: Compare imputation accuracy of QUILT2(msPBWT) against QUILT2(L&S) with "rare_af_threshold=0.001" (default) using the big UKB reference panel. We note that QUILT2 involves two computational innovations, i.e. the msPBWT and "impute_rare_common", which we referred to as QUILT2(msPBWT) explicitly here. Further, we note that the original QUILT1 is computationally expensive to run for very big reference panel. Therefore, we added the "impute_rare_common" strategy and retained the haplotype selection algorithm in QUILT1 based on Li and Stephens, which we referred to as QUILT2(L&S).

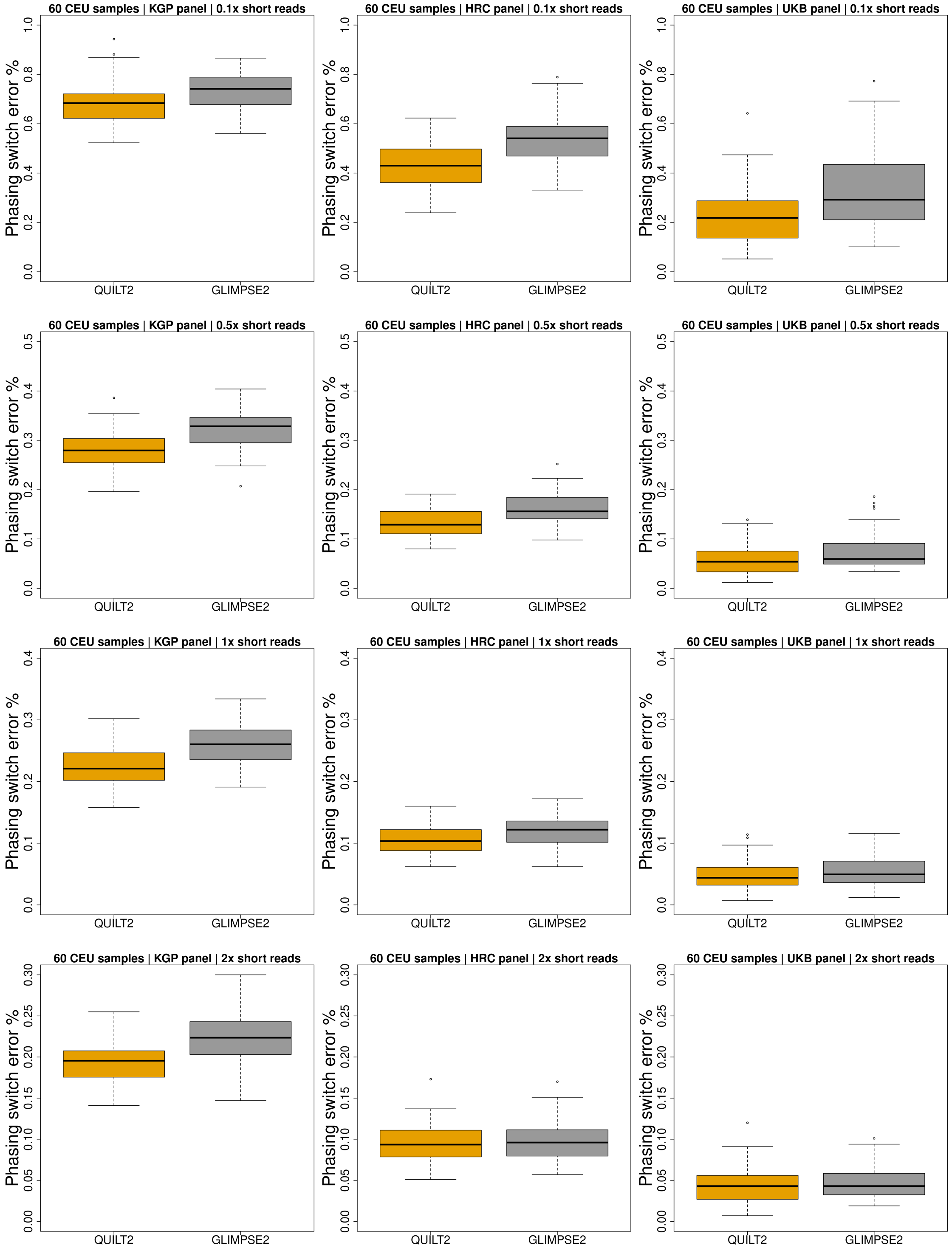

Supplementary Figure 3: Phasing performance of QUILT2 and GLIMPSE2 for 60 individuals (parents) in 30 CEU trios.

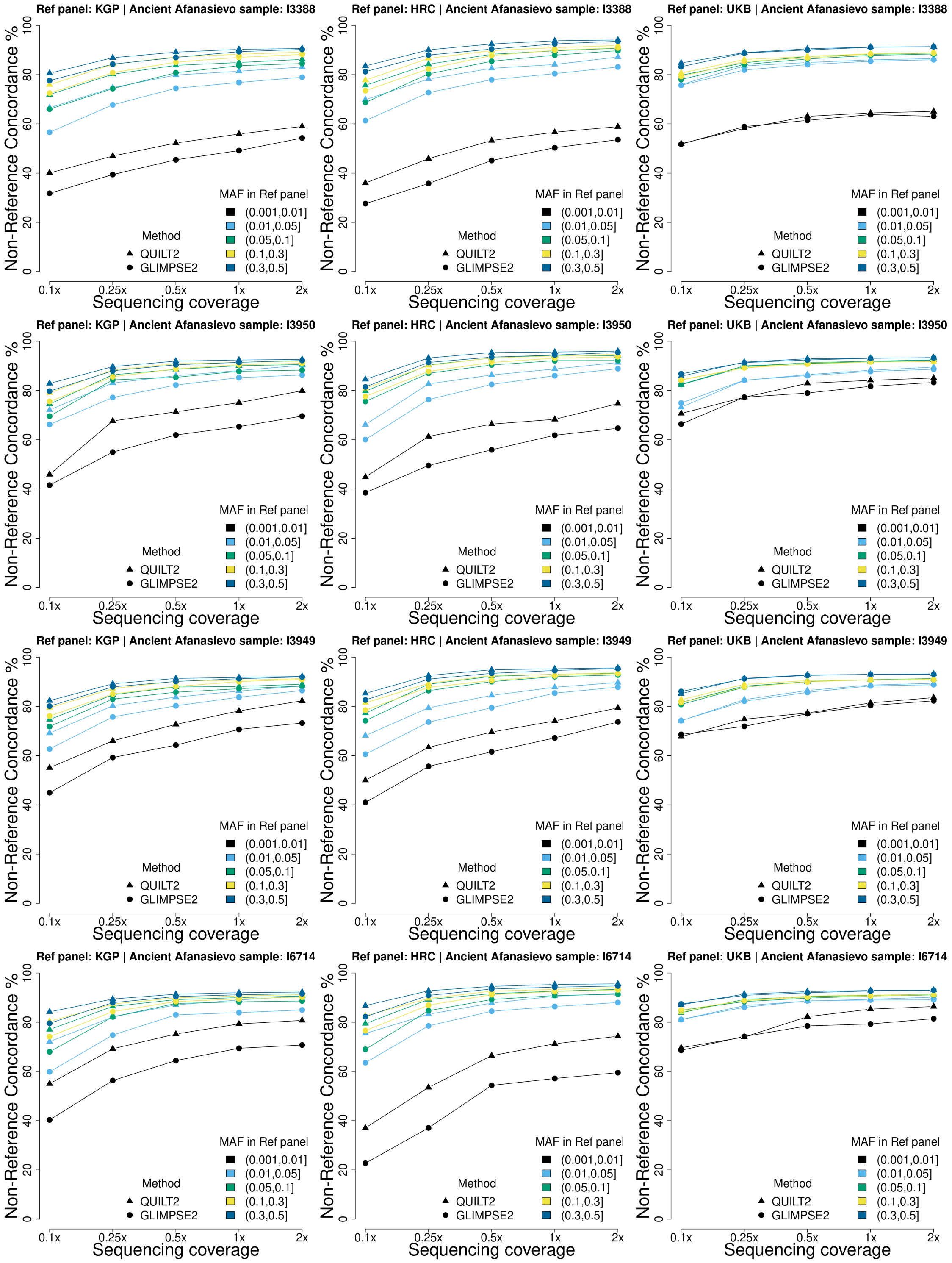

Supplementary Figure 4: Imputation accuracy for ancient DNA using different reference panels.

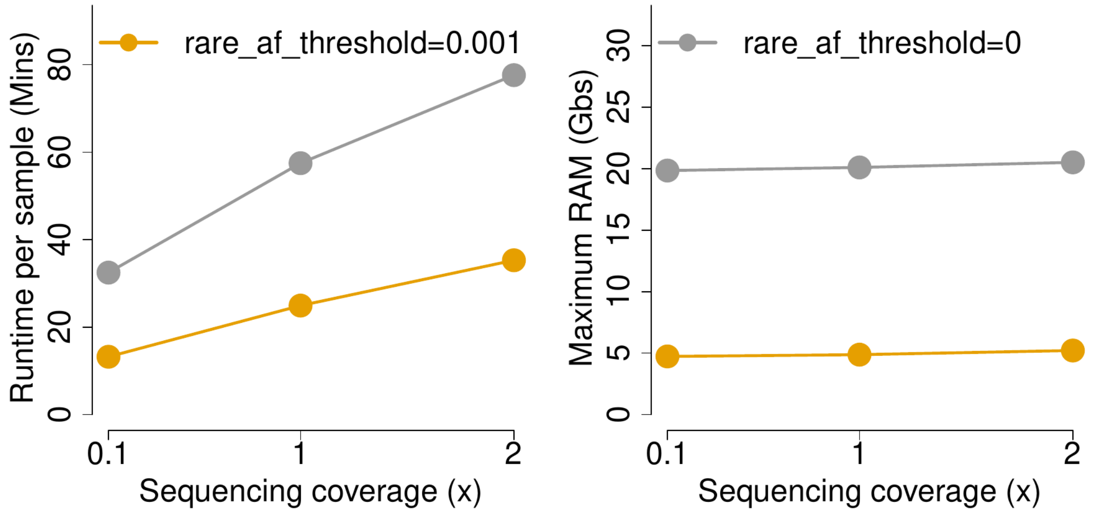

Supplementary Figure 5: Effect of rare_af_threshold on computational performance when imputing all variants. A subset of the UKB-GEL imputed reference panel was used (N=400,000, M=7,179,683), and QUILT2 run with 10 samples as input. QUILT2 with option rare_af_threshold=0 will build indices of msPBWT for all variants, while with rare_af_threshold=0.001 QUILT2 only builds indices of msPBWT for variants with MAF > rare_af_threshold.

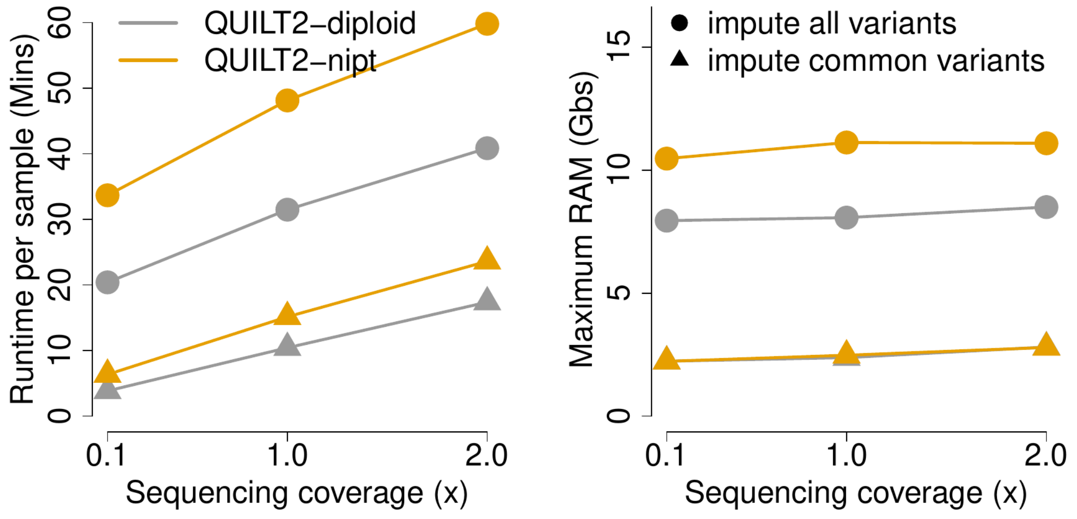

Supplementary Figure 6: Effect of impute_rare_common on computational performance when rare_af_threshold=0.001. The UKB-200K WGS reference panel was used(N=400,000, M=14,075,021), and QUILT2 run with 30 samples as input. Note that QUILT2 can prepare the reference panel for all variants, while users can choose whether to impute only common variants or not. For most GWAS on common variants, run QUILT2 with either diploid or nipt mode and impute_rare_common=FALSE can save a lot of computations.

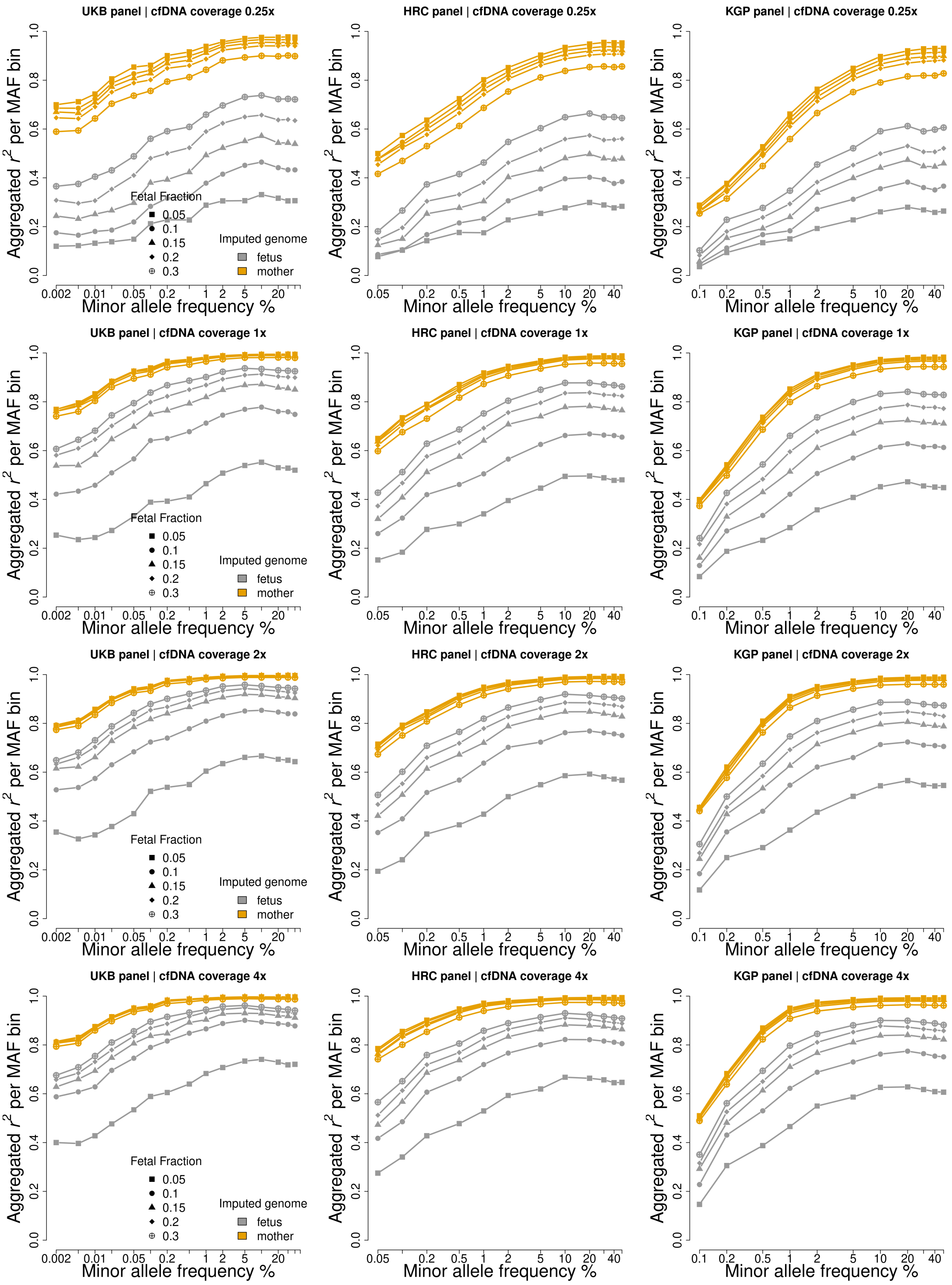

Supplementary Figure 7: Imputation accuracy of both maternal and fetal genomes across different NIPT coverage and fetal fraction using QUILT2-nipt method for 30 NIPT (CEU) samples.

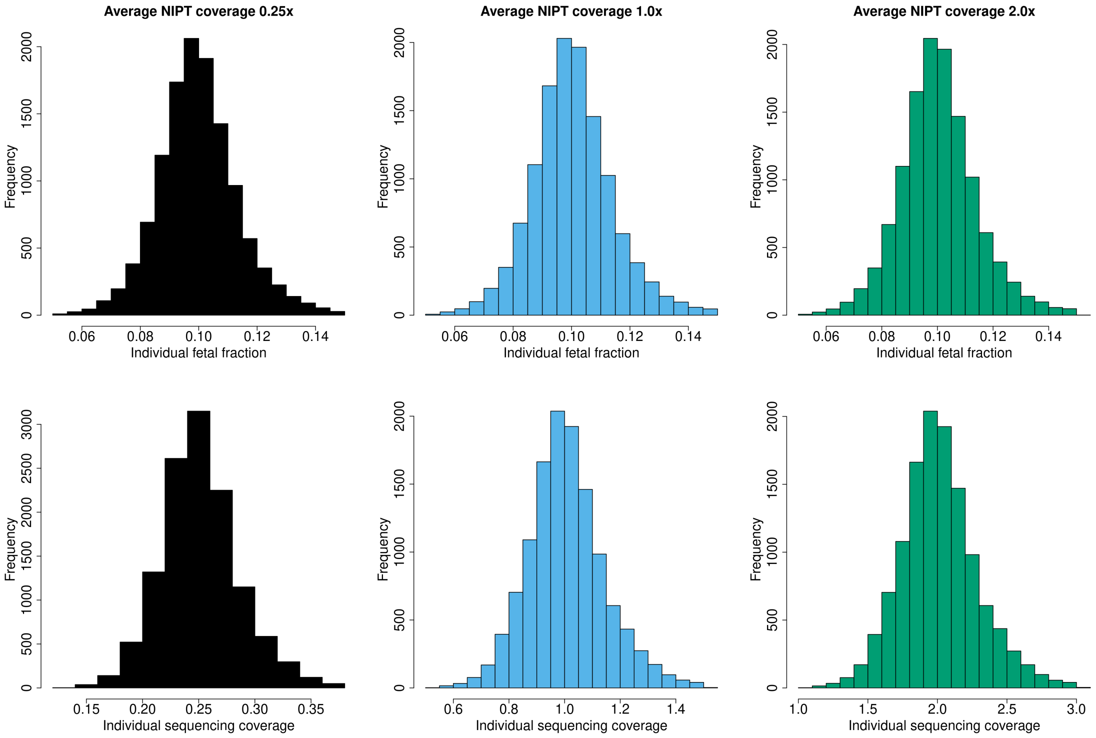

Supplementary Figure 8: Distribution of sequencing coverage and fetal fraction for 11,028 NIPT samples

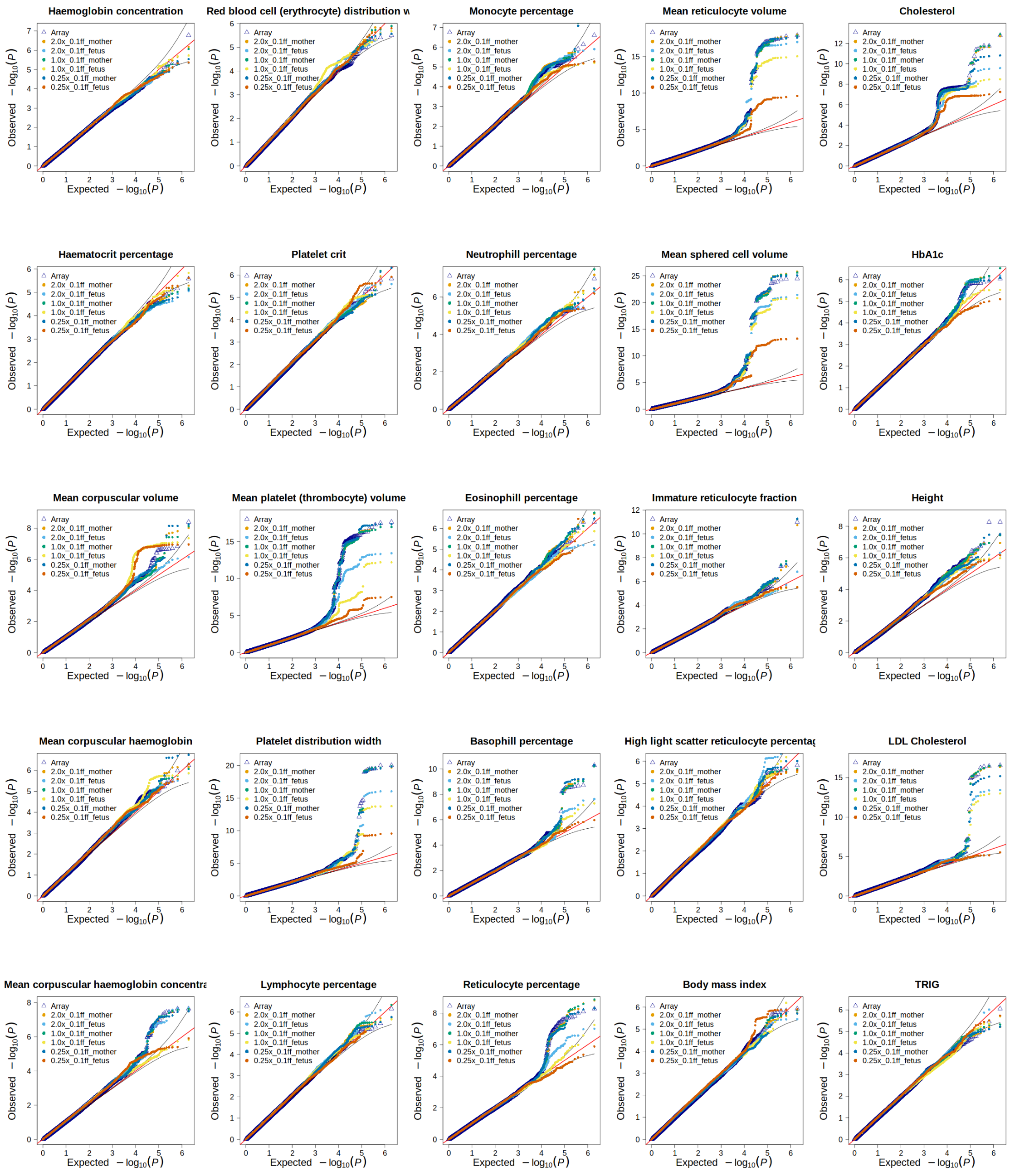

Supplementary Figure 9: Q-Q plots with around 1 million SNPs on chrmosome 1 for GWAS on imputed cfDNA with mean 10% fetal fraction.

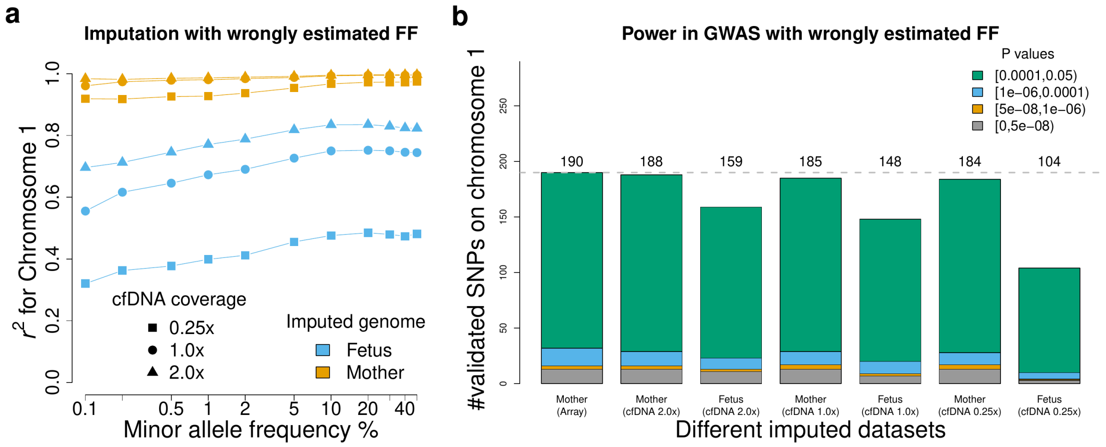

Supplementary Figure 10: Imputation performance and power in GWAS when there is bias in estimating fetal fraction. The real fetal fraction of each sample is shown in Supplementary Figure 8, while we run QUILT2-nipt with $FF = 0.1$ for all samples.

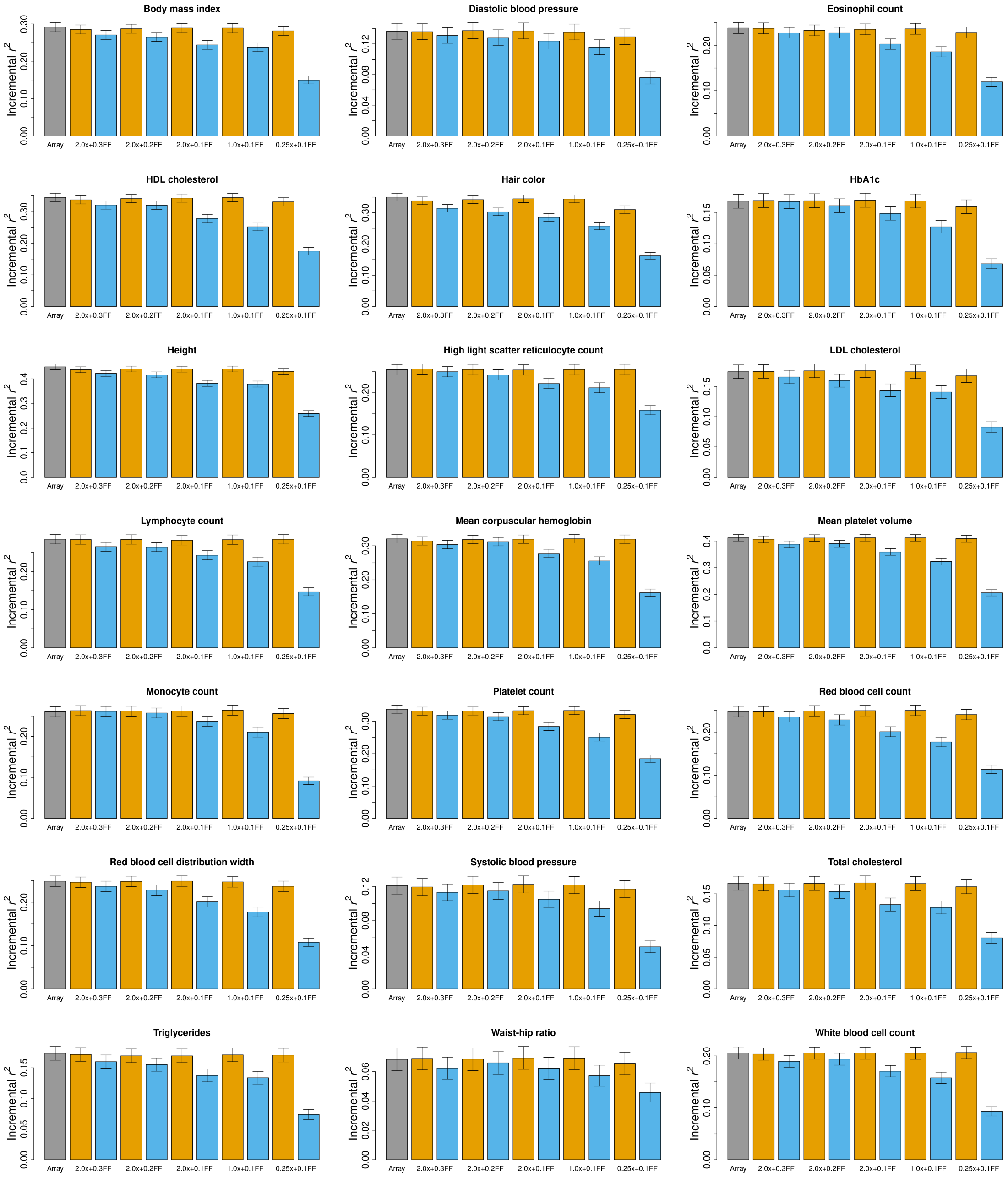

Supplementary Figure 11: PRS accuracy of each trait with different imputed datasets. The array-imputed data (gray) is the baseline. Then given different low coverage NIPT sequencing data and fetal fraction, there are imputed mother (yellow) and fetus (blue) genomes using QUILT2-nipt method.

### 3 Supplementary Tables

Supplementary Table 1: Number of SNPs for each reference panel, stratified by the minor allele frequency in gnomAD v3.1.2.

| MAF(gnomAD) | UKB | HRC | KGP |
|---|---|---|---|
| [0,0.0001] | 2625187 | 256957 | 853000 |
| (0.0001,0.0002] | 243964 | 103047 | 100898 |
| (0.0002,0.0005] | 158021 | 131209 | 92359 |
| (0.0005,0.001] | 62682 | 63353 | 47194 |
| (0.001,0.002] | 44870 | 43550 | 38083 |
| (0.002,0.005] | 44852 | 42032 | 42450 |
| (0.005,0.01] | 27163 | 25584 | 26551 |
| (0.01,0.02] | 23822 | 22487 | 23383 |
| (0.02,0.05] | 30504 | 29110 | 30187 |
| (0.05,0.1] | 23983 | 23124 | 24086 |
| (0.1,0.2] | 28254 | 26914 | 28599 |
| (0.2,0.3] | 18631 | 17926 | 18898 |
| (0.3,0.4] | 13490 | 13109 | 13799 |
| (0.4,0.5] | 11630 | 11254 | 11989 |

Supplementary Table 2: With the UKB panel, genotype imputation accuracy ($r^2$) for 60 CEU samples with short Illumina reads on chromosome 20.

| MAF | #SNPs | QUILT2 | GLIMPSE2 | COV |
|---|---|---|---|---|
| [0,1e-05] | 944143 | 0.548 | 0.492 | 0.1x |
| (1e-05,2e-05] | 717636 | 0.531 | 0.448 | 0.1x |
| (2e-05,5e-05] | 619703 | 0.586 | 0.5 | 0.1x |
| (5e-05,0.0001] | 343705 | 0.601 | 0.526 | 0.1x |
| (0.0001,0.0002] | 243964 | 0.642 | 0.578 | 0.1x |
| (0.0002,0.0005] | 158021 | 0.688 | 0.635 | 0.1x |
| (0.0005,0.001] | 62682 | 0.728 | 0.674 | 0.1x |
| (0.001,0.002] | 44870 | 0.76 | 0.71 | 0.1x |
| (0.002,0.005] | 44852 | 0.803 | 0.757 | 0.1x |
| (0.005,0.01] | 27163 | 0.832 | 0.792 | 0.1x |
| (0.01,0.02] | 23822 | 0.858 | 0.822 | 0.1x |
| (0.02,0.05] | 30504 | 0.896 | 0.865 | 0.1x |
| (0.05,0.1] | 23983 | 0.917 | 0.89 | 0.1x |
| (0.1,0.2] | 28254 | 0.927 | 0.901 | 0.1x |
| (0.2,0.3] | 18631 | 0.931 | 0.908 | 0.1x |
| (0.3,0.4] | 13490 | 0.932 | 0.909 | 0.1x |
| (0.4,0.5] | 11630 | 0.932 | 0.907 | 0.1x |
| [0,1e-05] | 944143 | 0.557 | 0.548 | 0.25x |
| (1e-05,2e-05] | 717636 | 0.614 | 0.566 | 0.25x |
| (2e-05,5e-05] | 619703 | 0.685 | 0.637 | 0.25x |
| (5e-05,0.0001] | 343705 | 0.71 | 0.651 | 0.25x |
| (0.0001,0.0002] | 243964 | 0.764 | 0.721 | 0.25x |
| (0.0002,0.0005] | 158021 | 0.818 | 0.776 | 0.25x |
| (0.0005,0.001] | 62682 | 0.866 | 0.829 | 0.25x |
| (0.001,0.002] | 44870 | 0.887 | 0.856 | 0.25x |
| (0.002,0.005] | 44852 | 0.917 | 0.895 | 0.25x |
| (0.005,0.01] | 27163 | 0.935 | 0.916 | 0.25x |
| (0.01,0.02] | 23822 | 0.951 | 0.937 | 0.25x |
| (0.02,0.05] | 30504 | 0.968 | 0.959 | 0.25x |
| (0.05,0.1] | 23983 | 0.979 | 0.972 | 0.25x |
| (0.1,0.2] | 28254 | 0.983 | 0.978 | 0.25x |
| (0.2,0.3] | 18631 | 0.984 | 0.979 | 0.25x |
| (0.3,0.4] | 13490 | 0.984 | 0.979 | 0.25x |
| (0.4,0.5] | 11630 | 0.984 | 0.979 | 0.25x |
| [0,1e-05] | 944143 | 0.561 | 0.547 | 0.5x |
| (1e-05,2e-05] | 717636 | 0.642 | 0.619 | 0.5x |
| (2e-05,5e-05] | 619703 | 0.719 | 0.688 | 0.5x |
| (5e-05,0.0001] | 343705 | 0.74 | 0.71 | 0.5x |
| (0.0001,0.0002] | 243964 | 0.803 | 0.78 | 0.5x |
| (0.0002,0.0005] | 158021 | 0.855 | 0.835 | 0.5x |
| (0.0005,0.001] | 62682 | 0.907 | 0.889 | 0.5x |
| (0.001,0.002] | 44870 | 0.924 | 0.912 | 0.5x |
| (0.002,0.005] | 44852 | 0.952 | 0.942 | 0.5x |
| (0.005,0.01] | 27163 | 0.963 | 0.956 | 0.5x |
| (0.01,0.02] | 23822 | 0.974 | 0.969 | 0.5x |

Continued on next page

Continued from previous page

| MAF | #SNPs | QUILT2 | GLIMPSE2 | COV |
|---|---|---|---|---|
| (0.02,0.05] | 30504 | 0.984 | 0.98 | 0.5x |
| (0.05,0.1] | 23983 | 0.99 | 0.988 | 0.5x |
| (0.1,0.2] | 28254 | 0.992 | 0.991 | 0.5x |
| (0.2,0.3] | 18631 | 0.993 | 0.991 | 0.5x |
| (0.3,0.4] | 13490 | 0.993 | 0.992 | 0.5x |
| (0.4,0.5] | 11630 | 0.992 | 0.991 | 0.5x |
| [0,1e-05] | 944143 | 0.566 | 0.544 | 1x |
| (1e-05,2e-05] | 717636 | 0.661 | 0.656 | 1x |
| (2e-05,5e-05] | 619703 | 0.742 | 0.725 | 1x |
| (5e-05,0.0001] | 343705 | 0.767 | 0.752 | 1x |
| (0.0001,0.0002] | 243964 | 0.833 | 0.832 | 1x |
| (0.0002,0.0005] | 158021 | 0.877 | 0.875 | 1x |
| (0.0005,0.001] | 62682 | 0.921 | 0.922 | 1x |
| (0.001,0.002] | 44870 | 0.94 | 0.938 | 1x |
| (0.002,0.005] | 44852 | 0.966 | 0.966 | 1x |
| (0.005,0.01] | 27163 | 0.977 | 0.974 | 1x |
| (0.01,0.02] | 23822 | 0.984 | 0.982 | 1x |
| (0.02,0.05] | 30504 | 0.99 | 0.99 | 1x |
| (0.05,0.1] | 23983 | 0.994 | 0.994 | 1x |
| (0.1,0.2] | 28254 | 0.995 | 0.995 | 1x |
| (0.2,0.3] | 18631 | 0.995 | 0.995 | 1x |
| (0.3,0.4] | 13490 | 0.996 | 0.995 | 1x |
| (0.4,0.5] | 11630 | 0.996 | 0.995 | 1x |
| [0,1e-05] | 944143 | 0.58 | 0.549 | 2x |
| (1e-05,2e-05] | 717636 | 0.689 | 0.687 | 2x |
| (2e-05,5e-05] | 619703 | 0.761 | 0.754 | 2x |
| (5e-05,0.0001] | 343705 | 0.788 | 0.786 | 2x |
| (0.0001,0.0002] | 243964 | 0.85 | 0.864 | 2x |
| (0.0002,0.0005] | 158021 | 0.89 | 0.899 | 2x |
| (0.0005,0.001] | 62682 | 0.937 | 0.941 | 2x |
| (0.001,0.002] | 44870 | 0.952 | 0.953 | 2x |
| (0.002,0.005] | 44852 | 0.976 | 0.979 | 2x |
| (0.005,0.01] | 27163 | 0.984 | 0.983 | 2x |
| (0.01,0.02] | 23822 | 0.989 | 0.989 | 2x |
| (0.02,0.05] | 30504 | 0.994 | 0.993 | 2x |
| (0.05,0.1] | 23983 | 0.996 | 0.996 | 2x |
| (0.1,0.2] | 28254 | 0.996 | 0.996 | 2x |
| (0.2,0.3] | 18631 | 0.996 | 0.996 | 2x |
| (0.3,0.4] | 13490 | 0.997 | 0.997 | 2x |
| (0.4,0.5] | 11630 | 0.997 | 0.997 | 2x |

Supplementary Table 3: Imputation accuracy (F1-score) across 3 reference panels for 11 EUR samples with long ONT reads at common variants (MAF $\in (0.1, 0.2]$) on chromosome 20.

| Sample | COV | QUILT2 UKB | GLIMPSE2 UKB | QUILT2 HRC | GLIMPSE2 HRC | QUILT2 KGP | GLIMPSE2 KGP |
|---|---|---|---|---|---|---|---|
| HG00105 | 0.1x | 0.592 | 0.472 | 0.525 | 0.37 | 0.525 | 0.352 |
| HG00110 | 0.1x | 0.435 | 0.381 | 0.446 | 0.399 | 0.406 | 0.336 |
| HG00121 | 0.1x | 0.422 | 0.401 | 0.384 | 0.313 | 0.338 | 0.253 |
| HG00127 | 0.1x | 0.603 | 0.358 | 0.545 | 0.324 | 0.524 | 0.363 |
| HG00136 | 0.1x | 0.541 | 0.406 | 0.54 | 0.394 | 0.519 | 0.377 |
| HG00151 | 0.1x | 0.579 | 0.412 | 0.542 | 0.334 | 0.524 | 0.358 |
| HG00331 | 0.1x | 0.511 | 0.369 | 0.595 | 0.432 | 0.504 | 0.346 |
| HG01501 | 0.1x | 0.484 | 0.336 | 0.462 | 0.28 | 0.452 | 0.321 |
| HG01615 | 0.1x | 0.512 | 0.378 | 0.502 | 0.377 | 0.495 | 0.378 |
| HG01695 | 0.1x | 0.491 | 0.407 | 0.468 | 0.358 | 0.444 | 0.342 |
| HG01790 | 0.1x | 0.398 | 0.343 | 0.341 | 0.269 | 0.398 | 0.272 |
| HG00105 | 0.5x | 0.963 | 0.697 | 0.868 | 0.613 | 0.806 | 0.542 |
| HG00110 | 0.5x | 0.886 | 0.524 | 0.871 | 0.491 | 0.788 | 0.476 |
| HG00121 | 0.5x | 0.891 | 0.562 | 0.878 | 0.515 | 0.748 | 0.469 |
| HG00127 | 0.5x | 0.89 | 0.537 | 0.841 | 0.481 | 0.804 | 0.441 |
| HG00136 | 0.5x | 0.905 | 0.632 | 0.865 | 0.585 | 0.832 | 0.548 |
| HG00151 | 0.5x | 0.898 | 0.585 | 0.853 | 0.522 | 0.797 | 0.471 |
| HG00331 | 0.5x | 0.891 | 0.709 | 0.963 | 0.722 | 0.871 | 0.66 |
| HG01501 | 0.5x | 0.825 | 0.506 | 0.81 | 0.521 | 0.787 | 0.496 |
| HG01615 | 0.5x | 0.847 | 0.592 | 0.816 | 0.563 | 0.797 | 0.521 |
| HG01695 | 0.5x | 0.848 | 0.602 | 0.827 | 0.571 | 0.798 | 0.536 |
| HG01790 | 0.5x | 0.858 | 0.474 | 0.793 | 0.45 | 0.72 | 0.437 |
| HG00105 | 1x | 0.994 | 0.735 | 0.95 | 0.67 | 0.906 | 0.622 |
| HG00110 | 1x | 0.956 | 0.626 | 0.946 | 0.599 | 0.906 | 0.56 |
| HG00121 | 1x | 0.96 | 0.668 | 0.952 | 0.618 | 0.868 | 0.537 |
| HG00127 | 1x | 0.967 | 0.639 | 0.926 | 0.633 | 0.889 | 0.596 |
| HG00136 | 1x | 0.973 | 0.694 | 0.947 | 0.641 | 0.893 | 0.622 |
| HG00151 | 1x | 0.97 | 0.731 | 0.926 | 0.675 | 0.903 | 0.646 |
| HG00331 | 1x | 0.966 | 0.779 | 0.976 | 0.754 | 0.941 | 0.724 |
| HG01501 | 1x | 0.872 | 0.594 | 0.861 | 0.593 | 0.845 | 0.553 |
| HG01615 | 1x | 0.935 | 0.706 | 0.9 | 0.685 | 0.885 | 0.642 |
| HG01695 | 1x | 0.945 | 0.7 | 0.923 | 0.668 | 0.898 | 0.631 |
| HG01790 | 1x | 0.958 | 0.546 | 0.916 | 0.536 | 0.859 | 0.49 |
| HG00105 | 2x | 0.994 | 0.832 | 0.975 | 0.8 | 0.957 | 0.762 |
| HG00110 | 2x | 0.988 | 0.773 | 0.967 | 0.746 | 0.956 | 0.711 |
| HG00121 | 2x | 0.977 | 0.775 | 0.977 | 0.744 | 0.931 | 0.714 |
| HG00127 | 2x | 0.987 | 0.782 | 0.967 | 0.753 | 0.948 | 0.723 |
| HG00136 | 2x | 0.99 | 0.863 | 0.981 | 0.842 | 0.955 | 0.806 |
| HG00151 | 2x | 0.985 | 0.793 | 0.978 | 0.76 | 0.943 | 0.726 |
| HG00331 | 2x | 0.988 | 0.869 | 0.991 | 0.851 | 0.981 | 0.827 |
| HG01501 | 2x | 0.926 | 0.743 | 0.911 | 0.735 | 0.907 | 0.711 |
| HG01615 | 2x | 0.981 | 0.846 | 0.968 | 0.821 | 0.958 | 0.796 |
| HG01695 | 2x | 0.978 | 0.811 | 0.966 | 0.786 | 0.95 | 0.767 |
| HG01790 | 2x | 0.976 | 0.682 | 0.938 | 0.655 | 0.92 | 0.648 |

Supplementary Table 4: With the UKB panel, maternal genotype imputation accuracy ($r^2$) with QUILT2-nipt and QUILT2-diploid methods for 30 NIPT (CEU) samples across various sequencing coverages and fetal fractions.

| MAF | #SNPs | QUILT2-nipt | QUILT2-diploid | FF | COV |
|---|---|---|---|---|---|
| (1e-05,2e-05] | 717636 | 0.616 | 0.612 | 0.05 | 0.25x |
| (2e-05,5e-05] | 619703 | 0.7 | 0.697 | 0.05 | 0.25x |
| (5e-05,0.0001] | 343705 | 0.712 | 0.704 | 0.05 | 0.25x |
| (0.0001,0.0002] | 243964 | 0.744 | 0.739 | 0.05 | 0.25x |
| (0.0002,0.0005] | 158021 | 0.806 | 0.809 | 0.05 | 0.25x |
| (0.0005,0.001] | 62682 | 0.854 | 0.85 | 0.05 | 0.25x |
| (0.001,0.002] | 44870 | 0.862 | 0.862 | 0.05 | 0.25x |
| (0.002,0.005] | 44852 | 0.901 | 0.9 | 0.05 | 0.25x |
| (0.005,0.01] | 27163 | 0.916 | 0.909 | 0.05 | 0.25x |
| (0.01,0.02] | 23822 | 0.939 | 0.935 | 0.05 | 0.25x |
| (0.02,0.05] | 30504 | 0.957 | 0.953 | 0.05 | 0.25x |
| (0.05,0.1] | 23983 | 0.971 | 0.963 | 0.05 | 0.25x |
| (0.1,0.2] | 28254 | 0.976 | 0.969 | 0.05 | 0.25x |
| (0.2,0.3] | 18631 | 0.976 | 0.968 | 0.05 | 0.25x |
| (0.3,0.4] | 13490 | 0.978 | 0.968 | 0.05 | 0.25x |
| (0.4,0.5] | 11630 | 0.976 | 0.967 | 0.05 | 0.25x |
| (1e-05,2e-05] | 717636 | 0.606 | 0.564 | 0.1 | 0.25x |
| (2e-05,5e-05] | 619703 | 0.686 | 0.639 | 0.1 | 0.25x |
| (5e-05,0.0001] | 343705 | 0.685 | 0.647 | 0.1 | 0.25x |
| (0.0001,0.0002] | 243964 | 0.724 | 0.695 | 0.1 | 0.25x |
| (0.0002,0.0005] | 158021 | 0.788 | 0.754 | 0.1 | 0.25x |
| (0.0005,0.001] | 62682 | 0.827 | 0.8 | 0.1 | 0.25x |
| (0.001,0.002] | 44870 | 0.841 | 0.805 | 0.1 | 0.25x |
| (0.002,0.005] | 44852 | 0.885 | 0.862 | 0.1 | 0.25x |
| (0.005,0.01] | 27163 | 0.9 | 0.869 | 0.1 | 0.25x |
| (0.01,0.02] | 23822 | 0.921 | 0.888 | 0.1 | 0.25x |
| (0.02,0.05] | 30504 | 0.949 | 0.922 | 0.1 | 0.25x |
| (0.05,0.1] | 23983 | 0.961 | 0.932 | 0.1 | 0.25x |
| (0.1,0.2] | 28254 | 0.966 | 0.939 | 0.1 | 0.25x |
| (0.2,0.3] | 18631 | 0.965 | 0.933 | 0.1 | 0.25x |
| (0.3,0.4] | 13490 | 0.966 | 0.93 | 0.1 | 0.25x |
| (0.4,0.5] | 11630 | 0.966 | 0.93 | 0.1 | 0.25x |
| (1e-05,2e-05] | 717636 | 0.592 | 0.509 | 0.15 | 0.25x |
| (2e-05,5e-05] | 619703 | 0.67 | 0.59 | 0.15 | 0.25x |
| (5e-05,0.0001] | 343705 | 0.666 | 0.597 | 0.15 | 0.25x |
| (0.0001,0.0002] | 243964 | 0.709 | 0.635 | 0.15 | 0.25x |
| (0.0002,0.0005] | 158021 | 0.773 | 0.712 | 0.15 | 0.25x |
| (0.0005,0.001] | 62682 | 0.808 | 0.753 | 0.15 | 0.25x |
| (0.001,0.002] | 44870 | 0.828 | 0.762 | 0.15 | 0.25x |
| (0.002,0.005] | 44852 | 0.869 | 0.812 | 0.15 | 0.25x |
| (0.005,0.01] | 27163 | 0.883 | 0.819 | 0.15 | 0.25x |
| (0.01,0.02] | 23822 | 0.912 | 0.842 | 0.15 | 0.25x |
| (0.02,0.05] | 30504 | 0.938 | 0.883 | 0.15 | 0.25x |
| (0.05,0.1] | 23983 | 0.95 | 0.891 | 0.15 | 0.25x |

Continued on next page

Continued from previous page

| MAF | #SNPs | QUILT2-nipt | QUILT2-diploid | FF | COV |
|---|---|---|---|---|---|
| (0.1,0.2] | 28254 | 0.956 | 0.896 | 0.15 | 0.25x |
| (0.2,0.3] | 18631 | 0.955 | 0.889 | 0.15 | 0.25x |
| (0.3,0.4] | 13490 | 0.956 | 0.884 | 0.15 | 0.25x |
| (0.4,0.5] | 11630 | 0.955 | 0.884 | 0.15 | 0.25x |
| (1e-05,2e-05] | 717636 | 0.561 | 0.441 | 0.2 | 0.25x |
| (2e-05,5e-05] | 619703 | 0.646 | 0.528 | 0.2 | 0.25x |
| (5e-05,0.0001] | 343705 | 0.643 | 0.533 | 0.2 | 0.25x |
| (0.0001,0.0002] | 243964 | 0.691 | 0.578 | 0.2 | 0.25x |
| (0.0002,0.0005] | 158021 | 0.751 | 0.651 | 0.2 | 0.25x |
| (0.0005,0.001] | 62682 | 0.789 | 0.674 | 0.2 | 0.25x |
| (0.001,0.002] | 44870 | 0.801 | 0.683 | 0.2 | 0.25x |
| (0.002,0.005] | 44852 | 0.849 | 0.749 | 0.2 | 0.25x |
| (0.005,0.01] | 27163 | 0.86 | 0.762 | 0.2 | 0.25x |
| (0.01,0.02] | 23822 | 0.887 | 0.787 | 0.2 | 0.25x |
| (0.02,0.05] | 30504 | 0.923 | 0.833 | 0.2 | 0.25x |
| (0.05,0.1] | 23983 | 0.935 | 0.832 | 0.2 | 0.25x |
| (0.1,0.2] | 28254 | 0.941 | 0.842 | 0.2 | 0.25x |
| (0.2,0.3] | 18631 | 0.94 | 0.836 | 0.2 | 0.25x |
| (0.3,0.4] | 13490 | 0.943 | 0.826 | 0.2 | 0.25x |
| (0.4,0.5] | 11630 | 0.941 | 0.827 | 0.2 | 0.25x |
| (1e-05,2e-05] | 717636 | 0.533 | 0.336 | 0.3 | 0.25x |
| (2e-05,5e-05] | 619703 | 0.589 | 0.387 | 0.3 | 0.25x |
| (5e-05,0.0001] | 343705 | 0.594 | 0.414 | 0.3 | 0.25x |
| (0.0001,0.0002] | 243964 | 0.643 | 0.457 | 0.3 | 0.25x |
| (0.0002,0.0005] | 158021 | 0.704 | 0.525 | 0.3 | 0.25x |
| (0.0005,0.001] | 62682 | 0.737 | 0.552 | 0.3 | 0.25x |
| (0.001,0.002] | 44870 | 0.756 | 0.541 | 0.3 | 0.25x |
| (0.002,0.005] | 44852 | 0.794 | 0.626 | 0.3 | 0.25x |
| (0.005,0.01] | 27163 | 0.812 | 0.637 | 0.3 | 0.25x |
| (0.01,0.02] | 23822 | 0.843 | 0.677 | 0.3 | 0.25x |
| (0.02,0.05] | 30504 | 0.881 | 0.732 | 0.3 | 0.25x |
| (0.05,0.1] | 23983 | 0.893 | 0.721 | 0.3 | 0.25x |
| (0.1,0.2] | 28254 | 0.9 | 0.729 | 0.3 | 0.25x |
| (0.2,0.3] | 18631 | 0.898 | 0.724 | 0.3 | 0.25x |
| (0.3,0.4] | 13490 | 0.901 | 0.709 | 0.3 | 0.25x |
| (0.4,0.5] | 11630 | 0.899 | 0.71 | 0.3 | 0.25x |
| (1e-05,2e-05] | 717636 | 0.676 | 0.677 | 0.05 | 1x |
| (2e-05,5e-05] | 619703 | 0.769 | 0.768 | 0.05 | 1x |
| (5e-05,0.0001] | 343705 | 0.796 | 0.785 | 0.05 | 1x |
| (0.0001,0.0002] | 243964 | 0.833 | 0.826 | 0.05 | 1x |
| (0.0002,0.0005] | 158021 | 0.884 | 0.884 | 0.05 | 1x |
| (0.0005,0.001] | 62682 | 0.926 | 0.915 | 0.05 | 1x |
| (0.001,0.002] | 44870 | 0.939 | 0.935 | 0.05 | 1x |
| (0.002,0.005] | 44852 | 0.966 | 0.96 | 0.05 | 1x |
| (0.005,0.01] | 27163 | 0.974 | 0.967 | 0.05 | 1x |
| (0.01,0.02] | 23822 | 0.983 | 0.979 | 0.05 | 1x |
| (0.02,0.05] | 30504 | 0.989 | 0.984 | 0.05 | 1x |
| (0.05,0.1] | 23983 | 0.993 | 0.987 | 0.05 | 1x |

Continued on next page

Continued from previous page

| MAF | #SNPs | QUILT2-nipt | QUILT2-diploid | FF | COV |
|---|---|---|---|---|---|
| (0.1,0.2] | 28254 | 0.995 | 0.988 | 0.05 | 1x |
| (0.2,0.3] | 18631 | 0.995 | 0.986 | 0.05 | 1x |
| (0.3,0.4] | 13490 | 0.996 | 0.985 | 0.05 | 1x |
| (0.4,0.5] | 11630 | 0.995 | 0.986 | 0.05 | 1x |
| (1e-05,2e-05] | 717636 | 0.677 | 0.639 | 0.1 | 1x |
| (2e-05,5e-05] | 619703 | 0.769 | 0.741 | 0.1 | 1x |
| (5e-05,0.0001] | 343705 | 0.792 | 0.755 | 0.1 | 1x |
| (0.0001,0.0002] | 243964 | 0.831 | 0.799 | 0.1 | 1x |
| (0.0002,0.0005] | 158021 | 0.881 | 0.858 | 0.1 | 1x |
| (0.0005,0.001] | 62682 | 0.924 | 0.895 | 0.1 | 1x |
| (0.001,0.002] | 44870 | 0.935 | 0.907 | 0.1 | 1x |
| (0.002,0.005] | 44852 | 0.963 | 0.941 | 0.1 | 1x |
| (0.005,0.01] | 27163 | 0.972 | 0.945 | 0.1 | 1x |
| (0.01,0.02] | 23822 | 0.981 | 0.958 | 0.1 | 1x |
| (0.02,0.05] | 30504 | 0.988 | 0.966 | 0.1 | 1x |
| (0.05,0.1] | 23983 | 0.992 | 0.962 | 0.1 | 1x |
| (0.1,0.2] | 28254 | 0.993 | 0.962 | 0.1 | 1x |
| (0.2,0.3] | 18631 | 0.994 | 0.956 | 0.1 | 1x |
| (0.3,0.4] | 13490 | 0.994 | 0.954 | 0.1 | 1x |
| (0.4,0.5] | 11630 | 0.994 | 0.953 | 0.1 | 1x |
| (1e-05,2e-05] | 717636 | 0.673 | 0.609 | 0.15 | 1x |
| (2e-05,5e-05] | 619703 | 0.761 | 0.701 | 0.15 | 1x |
| (5e-05,0.0001] | 343705 | 0.786 | 0.706 | 0.15 | 1x |
| (0.0001,0.0002] | 243964 | 0.825 | 0.76 | 0.15 | 1x |
| (0.0002,0.0005] | 158021 | 0.881 | 0.821 | 0.15 | 1x |
| (0.0005,0.001] | 62682 | 0.923 | 0.863 | 0.15 | 1x |
| (0.001,0.002] | 44870 | 0.931 | 0.87 | 0.15 | 1x |
| (0.002,0.005] | 44852 | 0.959 | 0.91 | 0.15 | 1x |
| (0.005,0.01] | 27163 | 0.969 | 0.909 | 0.15 | 1x |
| (0.01,0.02] | 23822 | 0.979 | 0.915 | 0.15 | 1x |
| (0.02,0.05] | 30504 | 0.986 | 0.93 | 0.15 | 1x |
| (0.05,0.1] | 23983 | 0.991 | 0.916 | 0.15 | 1x |
| (0.1,0.2] | 28254 | 0.992 | 0.916 | 0.15 | 1x |
| (0.2,0.3] | 18631 | 0.992 | 0.908 | 0.15 | 1x |
| (0.3,0.4] | 13490 | 0.993 | 0.901 | 0.15 | 1x |
| (0.4,0.5] | 11630 | 0.992 | 0.899 | 0.15 | 1x |
| (1e-05,2e-05] | 717636 | 0.671 | 0.56 | 0.2 | 1x |
| (2e-05,5e-05] | 619703 | 0.761 | 0.649 | 0.2 | 1x |
| (5e-05,0.0001] | 343705 | 0.781 | 0.662 | 0.2 | 1x |
| (0.0001,0.0002] | 243964 | 0.815 | 0.705 | 0.2 | 1x |
| (0.0002,0.0005] | 158021 | 0.874 | 0.778 | 0.2 | 1x |
| (0.0005,0.001] | 62682 | 0.915 | 0.817 | 0.2 | 1x |
| (0.001,0.002] | 44870 | 0.926 | 0.82 | 0.2 | 1x |
| (0.002,0.005] | 44852 | 0.955 | 0.867 | 0.2 | 1x |
| (0.005,0.01] | 27163 | 0.965 | 0.867 | 0.2 | 1x |
| (0.01,0.02] | 23822 | 0.976 | 0.872 | 0.2 | 1x |
| (0.02,0.05] | 30504 | 0.984 | 0.883 | 0.2 | 1x |
| (0.05,0.1] | 23983 | 0.989 | 0.863 | 0.2 | 1x |

Continued on next page

Continued from previous page

| MAF | #SNPs | QUILT2-nipt | QUILT2-diploid | FF | COV |
|---|---|---|---|---|---|
| (0.1,0.2] | 28254 | 0.99 | 0.857 | 0.2 | 1x |
| (0.2,0.3] | 18631 | 0.99 | 0.846 | 0.2 | 1x |
| (0.3,0.4] | 13490 | 0.99 | 0.837 | 0.2 | 1x |
| (0.4,0.5] | 11630 | 0.989 | 0.831 | 0.2 | 1x |
| (1e-05,2e-05] | 717636 | 0.654 | 0.467 | 0.3 | 1x |
| (2e-05,5e-05] | 619703 | 0.741 | 0.547 | 0.3 | 1x |
| (5e-05,0.0001] | 343705 | 0.76 | 0.551 | 0.3 | 1x |
| (0.0001,0.0002] | 243964 | 0.804 | 0.607 | 0.3 | 1x |
| (0.0002,0.0005] | 158021 | 0.86 | 0.66 | 0.3 | 1x |
| (0.0005,0.001] | 62682 | 0.895 | 0.696 | 0.3 | 1x |
| (0.001,0.002] | 44870 | 0.912 | 0.711 | 0.3 | 1x |
| (0.002,0.005] | 44852 | 0.941 | 0.758 | 0.3 | 1x |
| (0.005,0.01] | 27163 | 0.952 | 0.747 | 0.3 | 1x |
| (0.01,0.02] | 23822 | 0.965 | 0.759 | 0.3 | 1x |
| (0.02,0.05] | 30504 | 0.974 | 0.773 | 0.3 | 1x |
| (0.05,0.1] | 23983 | 0.981 | 0.751 | 0.3 | 1x |
| (0.1,0.2] | 28254 | 0.981 | 0.739 | 0.3 | 1x |
| (0.2,0.3] | 18631 | 0.981 | 0.726 | 0.3 | 1x |
| (0.3,0.4] | 13490 | 0.982 | 0.706 | 0.3 | 1x |
| (0.4,0.5] | 11630 | 0.98 | 0.697 | 0.3 | 1x |
| (1e-05,2e-05] | 717636 | 0.705 | 0.697 | 0.05 | 2x |
| (2e-05,5e-05] | 619703 | 0.791 | 0.786 | 0.05 | 2x |
| (5e-05,0.0001] | 343705 | 0.812 | 0.817 | 0.05 | 2x |
| (0.0001,0.0002] | 243964 | 0.857 | 0.858 | 0.05 | 2x |
| (0.0002,0.0005] | 158021 | 0.902 | 0.9 | 0.05 | 2x |
| (0.0005,0.001] | 62682 | 0.942 | 0.939 | 0.05 | 2x |
| (0.001,0.002] | 44870 | 0.952 | 0.951 | 0.05 | 2x |
| (0.002,0.005] | 44852 | 0.976 | 0.973 | 0.05 | 2x |
| (0.005,0.01] | 27163 | 0.983 | 0.979 | 0.05 | 2x |
| (0.01,0.02] | 23822 | 0.989 | 0.986 | 0.05 | 2x |
| (0.02,0.05] | 30504 | 0.993 | 0.99 | 0.05 | 2x |
| (0.05,0.1] | 23983 | 0.996 | 0.989 | 0.05 | 2x |
| (0.1,0.2] | 28254 | 0.996 | 0.99 | 0.05 | 2x |
| (0.2,0.3] | 18631 | 0.996 | 0.988 | 0.05 | 2x |
| (0.3,0.4] | 13490 | 0.997 | 0.988 | 0.05 | 2x |
| (0.4,0.5] | 11630 | 0.996 | 0.988 | 0.05 | 2x |
| (1e-05,2e-05] | 717636 | 0.705 | 0.685 | 0.1 | 2x |
| (2e-05,5e-05] | 619703 | 0.794 | 0.773 | 0.1 | 2x |
| (5e-05,0.0001] | 343705 | 0.814 | 0.791 | 0.1 | 2x |
| (0.0001,0.0002] | 243964 | 0.851 | 0.839 | 0.1 | 2x |
| (0.0002,0.0005] | 158021 | 0.899 | 0.88 | 0.1 | 2x |
| (0.0005,0.001] | 62682 | 0.941 | 0.92 | 0.1 | 2x |
| (0.001,0.002] | 44870 | 0.951 | 0.932 | 0.1 | 2x |
| (0.002,0.005] | 44852 | 0.975 | 0.96 | 0.1 | 2x |
| (0.005,0.01] | 27163 | 0.982 | 0.964 | 0.1 | 2x |
| (0.01,0.02] | 23822 | 0.988 | 0.961 | 0.1 | 2x |
| (0.02,0.05] | 30504 | 0.992 | 0.973 | 0.1 | 2x |
| (0.05,0.1] | 23983 | 0.995 | 0.966 | 0.1 | 2x |

Continued on next page

Continued from previous page

| MAF | #SNPs | QUILT2-nipt | QUILT2-diploid | FF | COV |
|---|---|---|---|---|---|
| (0.1,0.2] | 28254 | 0.996 | 0.963 | 0.1 | 2x |
| (0.2,0.3] | 18631 | 0.995 | 0.959 | 0.1 | 2x |
| (0.3,0.4] | 13490 | 0.996 | 0.957 | 0.1 | 2x |
| (0.4,0.5] | 11630 | 0.996 | 0.955 | 0.1 | 2x |
| (1e-05,2e-05] | 717636 | 0.7 | 0.652 | 0.15 | 2x |
| (2e-05,5e-05] | 619703 | 0.787 | 0.74 | 0.15 | 2x |
| (5e-05,0.0001] | 343705 | 0.807 | 0.752 | 0.15 | 2x |
| (0.0001,0.0002] | 243964 | 0.851 | 0.811 | 0.15 | 2x |
| (0.0002,0.0005] | 158021 | 0.9 | 0.849 | 0.15 | 2x |
| (0.0005,0.001] | 62682 | 0.937 | 0.899 | 0.15 | 2x |
| (0.001,0.002] | 44870 | 0.948 | 0.905 | 0.15 | 2x |
| (0.002,0.005] | 44852 | 0.973 | 0.937 | 0.15 | 2x |
| (0.005,0.01] | 27163 | 0.981 | 0.936 | 0.15 | 2x |
| (0.01,0.02] | 23822 | 0.988 | 0.934 | 0.15 | 2x |
| (0.02,0.05] | 30504 | 0.991 | 0.934 | 0.15 | 2x |
| (0.05,0.1] | 23983 | 0.994 | 0.922 | 0.15 | 2x |
| (0.1,0.2] | 28254 | 0.995 | 0.914 | 0.15 | 2x |
| (0.2,0.3] | 18631 | 0.995 | 0.906 | 0.15 | 2x |
| (0.3,0.4] | 13490 | 0.995 | 0.898 | 0.15 | 2x |
| (0.4,0.5] | 11630 | 0.995 | 0.893 | 0.15 | 2x |
| (1e-05,2e-05] | 717636 | 0.698 | 0.616 | 0.2 | 2x |
| (2e-05,5e-05] | 619703 | 0.784 | 0.71 | 0.2 | 2x |
| (5e-05,0.0001] | 343705 | 0.805 | 0.708 | 0.2 | 2x |
| (0.0001,0.0002] | 243964 | 0.847 | 0.77 | 0.2 | 2x |
| (0.0002,0.0005] | 158021 | 0.896 | 0.82 | 0.2 | 2x |
| (0.0005,0.001] | 62682 | 0.934 | 0.858 | 0.2 | 2x |
| (0.001,0.002] | 44870 | 0.944 | 0.877 | 0.2 | 2x |
| (0.002,0.005] | 44852 | 0.971 | 0.904 | 0.2 | 2x |
| (0.005,0.01] | 27163 | 0.978 | 0.892 | 0.2 | 2x |
| (0.01,0.02] | 23822 | 0.986 | 0.886 | 0.2 | 2x |
| (0.02,0.05] | 30504 | 0.99 | 0.891 | 0.2 | 2x |
| (0.05,0.1] | 23983 | 0.993 | 0.868 | 0.2 | 2x |
| (0.1,0.2] | 28254 | 0.994 | 0.856 | 0.2 | 2x |
| (0.2,0.3] | 18631 | 0.993 | 0.843 | 0.2 | 2x |
| (0.3,0.4] | 13490 | 0.994 | 0.826 | 0.2 | 2x |
| (0.4,0.5] | 11630 | 0.993 | 0.821 | 0.2 | 2x |
| (1e-05,2e-05] | 717636 | 0.689 | 0.54 | 0.3 | 2x |
| (2e-05,5e-05] | 619703 | 0.774 | 0.621 | 0.3 | 2x |
| (5e-05,0.0001] | 343705 | 0.79 | 0.624 | 0.3 | 2x |
| (0.0001,0.0002] | 243964 | 0.835 | 0.688 | 0.3 | 2x |
| (0.0002,0.0005] | 158021 | 0.884 | 0.735 | 0.3 | 2x |
| (0.0005,0.001] | 62682 | 0.925 | 0.76 | 0.3 | 2x |
| (0.001,0.002] | 44870 | 0.934 | 0.777 | 0.3 | 2x |
| (0.002,0.005] | 44852 | 0.962 | 0.809 | 0.3 | 2x |
| (0.005,0.01] | 27163 | 0.97 | 0.784 | 0.3 | 2x |
| (0.01,0.02] | 23822 | 0.98 | 0.785 | 0.3 | 2x |
| (0.02,0.05] | 30504 | 0.985 | 0.787 | 0.3 | 2x |
| (0.05,0.1] | 23983 | 0.988 | 0.758 | 0.3 | 2x |

Continued on next page

Continued from previous page

| MAF | #SNPs | QUILT2-nipt | QUILT2-diploid | FF | COV |
|---|---|---|---|---|---|
| (0.1,0.2] | 28254 | 0.988 | 0.739 | 0.3 | 2x |
| (0.2,0.3] | 18631 | 0.988 | 0.718 | 0.3 | 2x |
| (0.3,0.4] | 13490 | 0.988 | 0.694 | 0.3 | 2x |
| (0.4,0.5] | 11630 | 0.987 | 0.689 | 0.3 | 2x |
| (1e-05,2e-05] | 717636 | 0.725 | 0.731 | 0.05 | 4x |
| (2e-05,5e-05] | 619703 | 0.811 | 0.814 | 0.05 | 4x |
| (5e-05,0.0001] | 343705 | 0.83 | 0.836 | 0.05 | 4x |
| (0.0001,0.0002] | 243964 | 0.875 | 0.883 | 0.05 | 4x |
| (0.0002,0.0005] | 158021 | 0.915 | 0.916 | 0.05 | 4x |
| (0.0005,0.001] | 62682 | 0.952 | 0.952 | 0.05 | 4x |
| (0.001,0.002] | 44870 | 0.96 | 0.959 | 0.05 | 4x |
| (0.002,0.005] | 44852 | 0.984 | 0.983 | 0.05 | 4x |
| (0.005,0.01] | 27163 | 0.988 | 0.985 | 0.05 | 4x |
| (0.01,0.02] | 23822 | 0.992 | 0.991 | 0.05 | 4x |
| (0.02,0.05] | 30504 | 0.995 | 0.992 | 0.05 | 4x |
| (0.05,0.1] | 23983 | 0.997 | 0.992 | 0.05 | 4x |
| (0.1,0.2] | 28254 | 0.997 | 0.991 | 0.05 | 4x |
| (0.2,0.3] | 18631 | 0.997 | 0.99 | 0.05 | 4x |
| (0.3,0.4] | 13490 | 0.997 | 0.99 | 0.05 | 4x |
| (0.4,0.5] | 11630 | 0.997 | 0.989 | 0.05 | 4x |
| (1e-05,2e-05] | 717636 | 0.725 | 0.716 | 0.1 | 4x |
| (2e-05,5e-05] | 619703 | 0.814 | 0.801 | 0.1 | 4x |
| (5e-05,0.0001] | 343705 | 0.832 | 0.818 | 0.1 | 4x |
| (0.0001,0.0002] | 243964 | 0.875 | 0.874 | 0.1 | 4x |
| (0.0002,0.0005] | 158021 | 0.914 | 0.904 | 0.1 | 4x |
| (0.0005,0.001] | 62682 | 0.952 | 0.941 | 0.1 | 4x |
| (0.001,0.002] | 44870 | 0.958 | 0.945 | 0.1 | 4x |
| (0.002,0.005] | 44852 | 0.982 | 0.972 | 0.1 | 4x |
| (0.005,0.01] | 27163 | 0.988 | 0.975 | 0.1 | 4x |
| (0.01,0.02] | 23822 | 0.992 | 0.973 | 0.1 | 4x |
| (0.02,0.05] | 30504 | 0.995 | 0.974 | 0.1 | 4x |
| (0.05,0.1] | 23983 | 0.997 | 0.969 | 0.1 | 4x |
| (0.1,0.2] | 28254 | 0.996 | 0.964 | 0.1 | 4x |
| (0.2,0.3] | 18631 | 0.996 | 0.958 | 0.1 | 4x |
| (0.3,0.4] | 13490 | 0.997 | 0.955 | 0.1 | 4x |
| (0.4,0.5] | 11630 | 0.996 | 0.954 | 0.1 | 4x |
| (1e-05,2e-05] | 717636 | 0.723 | 0.693 | 0.15 | 4x |
| (2e-05,5e-05] | 619703 | 0.809 | 0.79 | 0.15 | 4x |
| (5e-05,0.0001] | 343705 | 0.823 | 0.795 | 0.15 | 4x |
| (0.0001,0.0002] | 243964 | 0.871 | 0.85 | 0.15 | 4x |
| (0.0002,0.0005] | 158021 | 0.911 | 0.881 | 0.15 | 4x |
| (0.0005,0.001] | 62682 | 0.949 | 0.92 | 0.15 | 4x |
| (0.001,0.002] | 44870 | 0.957 | 0.929 | 0.15 | 4x |
| (0.002,0.005] | 44852 | 0.982 | 0.957 | 0.15 | 4x |
| (0.005,0.01] | 27163 | 0.987 | 0.951 | 0.15 | 4x |
| (0.01,0.02] | 23822 | 0.991 | 0.946 | 0.15 | 4x |
| (0.02,0.05] | 30504 | 0.994 | 0.945 | 0.15 | 4x |
| (0.05,0.1] | 23983 | 0.996 | 0.926 | 0.15 | 4x |

Continued on next page

Continued from previous page

| MAF | #SNPs | QUILT2-nipt | QUILT2-diploid | FF | COV |
|---|---:|---:|---:|---:|---:|
| (0.1,0.2] | 28254 | 0.995 | 0.915 | 0.15 | 4x |
| (0.2,0.3] | 18631 | 0.995 | 0.903 | 0.15 | 4x |
| (0.3,0.4] | 13490 | 0.995 | 0.895 | 0.15 | 4x |
| (0.4,0.5] | 11630 | 0.995 | 0.89 | 0.15 | 4x |
| (1e-05,2e-05] | 717636 | 0.72 | 0.665 | 0.2 | 4x |
| (2e-05,5e-05] | 619703 | 0.807 | 0.754 | 0.2 | 4x |
| (5e-05,0.0001] | 343705 | 0.819 | 0.765 | 0.2 | 4x |
| (0.0001,0.0002] | 243964 | 0.868 | 0.824 | 0.2 | 4x |
| (0.0002,0.0005] | 158021 | 0.909 | 0.857 | 0.2 | 4x |
| (0.0005,0.001] | 62682 | 0.945 | 0.894 | 0.2 | 4x |
| (0.001,0.002] | 44870 | 0.952 | 0.897 | 0.2 | 4x |
| (0.002,0.005] | 44852 | 0.979 | 0.931 | 0.2 | 4x |
| (0.005,0.01] | 27163 | 0.986 | 0.911 | 0.2 | 4x |
| (0.01,0.02] | 23822 | 0.989 | 0.908 | 0.2 | 4x |
| (0.02,0.05] | 30504 | 0.992 | 0.901 | 0.2 | 4x |
| (0.05,0.1] | 23983 | 0.994 | 0.874 | 0.2 | 4x |
| (0.1,0.2] | 28254 | 0.994 | 0.853 | 0.2 | 4x |
| (0.2,0.3] | 18631 | 0.993 | 0.838 | 0.2 | 4x |
| (0.3,0.4] | 13490 | 0.994 | 0.822 | 0.2 | 4x |
| (0.4,0.5] | 11630 | 0.993 | 0.814 | 0.2 | 4x |
| (1e-05,2e-05] | 717636 | 0.7 | 0.598 | 0.3 | 4x |
| (2e-05,5e-05] | 619703 | 0.793 | 0.681 | 0.3 | 4x |
| (5e-05,0.0001] | 343705 | 0.807 | 0.672 | 0.3 | 4x |
| (0.0001,0.0002] | 243964 | 0.859 | 0.751 | 0.3 | 4x |
| (0.0002,0.0005] | 158021 | 0.898 | 0.789 | 0.3 | 4x |
| (0.0005,0.001] | 62682 | 0.935 | 0.809 | 0.3 | 4x |
| (0.001,0.002] | 44870 | 0.946 | 0.811 | 0.3 | 4x |
| (0.002,0.005] | 44852 | 0.97 | 0.841 | 0.3 | 4x |
| (0.005,0.01] | 27163 | 0.977 | 0.807 | 0.3 | 4x |
| (0.01,0.02] | 23822 | 0.984 | 0.81 | 0.3 | 4x |
| (0.02,0.05] | 30504 | 0.987 | 0.797 | 0.3 | 4x |
| (0.05,0.1] | 23983 | 0.989 | 0.761 | 0.3 | 4x |
| (0.1,0.2] | 28254 | 0.988 | 0.736 | 0.3 | 4x |
| (0.2,0.3] | 18631 | 0.988 | 0.712 | 0.3 | 4x |
| (0.3,0.4] | 13490 | 0.988 | 0.685 | 0.3 | 4x |
| (0.4,0.5] | 11630 | 0.987 | 0.678 | 0.3 | 4x |

Supplementary Table 5: With the UKB panel, maternal and fetal genotype imputation accuracy ($r^2$) with QUILT2-nipt method for 30 NIPT (CEU) samples across various sequencing coverages and fetal fractions.

| MAF | #SNPs | Mother | Fetus | FF | COV |
|---|---|---|---|---|---|
| (1e-05,2e-05] | 717636 | 0.499 | 0.096 | 0.05 | 0.1x |
| (2e-05,5e-05] | 619703 | 0.554 | 0.085 | 0.05 | 0.1x |
| (5e-05,0.0001] | 343705 | 0.564 | 0.078 | 0.05 | 0.1x |
| (0.0001,0.0002] | 243964 | 0.602 | 0.082 | 0.05 | 0.1x |
| (0.0002,0.0005] | 158021 | 0.648 | 0.076 | 0.05 | 0.1x |
| (0.0005,0.001] | 62682 | 0.699 | 0.094 | 0.05 | 0.1x |
| (0.001,0.002] | 44870 | 0.707 | 0.144 | 0.05 | 0.1x |
| (0.002,0.005] | 44852 | 0.758 | 0.188 | 0.05 | 0.1x |
| (0.005,0.01] | 27163 | 0.791 | 0.168 | 0.05 | 0.1x |
| (0.01,0.02] | 23822 | 0.826 | 0.187 | 0.05 | 0.1x |
| (0.02,0.05] | 30504 | 0.873 | 0.204 | 0.05 | 0.1x |
| (0.05,0.1] | 23983 | 0.89 | 0.217 | 0.05 | 0.1x |
| (0.1,0.2] | 28254 | 0.903 | 0.234 | 0.05 | 0.1x |
| (0.2,0.3] | 18631 | 0.904 | 0.225 | 0.05 | 0.1x |
| (0.3,0.4] | 13490 | 0.91 | 0.214 | 0.05 | 0.1x |
| (0.4,0.5] | 11630 | 0.908 | 0.223 | 0.05 | 0.1x |
| (1e-05,2e-05] | 717636 | 0.453 | 0.102 | 0.1 | 0.1x |
| (2e-05,5e-05] | 619703 | 0.512 | 0.107 | 0.1 | 0.1x |
| (5e-05,0.0001] | 343705 | 0.516 | 0.098 | 0.1 | 0.1x |
| (0.0001,0.0002] | 243964 | 0.563 | 0.102 | 0.1 | 0.1x |
| (0.0002,0.0005] | 158021 | 0.611 | 0.101 | 0.1 | 0.1x |
| (0.0005,0.001] | 62682 | 0.652 | 0.133 | 0.1 | 0.1x |
| (0.001,0.002] | 44870 | 0.668 | 0.183 | 0.1 | 0.1x |
| (0.002,0.005] | 44852 | 0.718 | 0.179 | 0.1 | 0.1x |
| (0.005,0.01] | 27163 | 0.75 | 0.19 | 0.1 | 0.1x |
| (0.01,0.02] | 23822 | 0.792 | 0.244 | 0.1 | 0.1x |
| (0.02,0.05] | 30504 | 0.839 | 0.257 | 0.1 | 0.1x |
| (0.05,0.1] | 23983 | 0.858 | 0.283 | 0.1 | 0.1x |
| (0.1,0.2] | 28254 | 0.87 | 0.305 | 0.1 | 0.1x |
| (0.2,0.3] | 18631 | 0.872 | 0.285 | 0.1 | 0.1x |
| (0.3,0.4] | 13490 | 0.88 | 0.274 | 0.1 | 0.1x |
| (0.4,0.5] | 11630 | 0.873 | 0.285 | 0.1 | 0.1x |
| (1e-05,2e-05] | 717636 | 0.406 | 0.1 | 0.15 | 0.1x |
| (2e-05,5e-05] | 619703 | 0.446 | 0.095 | 0.15 | 0.1x |
| (5e-05,0.0001] | 343705 | 0.468 | 0.094 | 0.15 | 0.1x |
| (0.0001,0.0002] | 243964 | 0.509 | 0.083 | 0.15 | 0.1x |
| (0.0002,0.0005] | 158021 | 0.567 | 0.11 | 0.15 | 0.1x |
| (0.0005,0.001] | 62682 | 0.602 | 0.142 | 0.15 | 0.1x |
| (0.001,0.002] | 44870 | 0.612 | 0.191 | 0.15 | 0.1x |
| (0.002,0.005] | 44852 | 0.673 | 0.216 | 0.15 | 0.1x |
| (0.005,0.01] | 27163 | 0.717 | 0.213 | 0.15 | 0.1x |
| (0.01,0.02] | 23822 | 0.761 | 0.257 | 0.15 | 0.1x |
| (0.02,0.05] | 30504 | 0.813 | 0.292 | 0.15 | 0.1x |
| (0.05,0.1] | 23983 | 0.826 | 0.326 | 0.15 | 0.1x |

Continued on next page

Continued from previous page

| MAF | #SNPs | Mother | Fetus | FF | COV |
|---|---|---|---|---|---|
| (0.1,0.2] | 28254 | 0.842 | 0.344 | 0.15 | 0.1x |
| (0.2,0.3] | 18631 | 0.844 | 0.329 | 0.15 | 0.1x |
| (0.3,0.4] | 13490 | 0.851 | 0.319 | 0.15 | 0.1x |
| (0.4,0.5] | 11630 | 0.848 | 0.331 | 0.15 | 0.1x |
| (1e-05,2e-05] | 717636 | 0.366 | 0.103 | 0.2 | 0.1x |
| (2e-05,5e-05] | 619703 | 0.417 | 0.111 | 0.2 | 0.1x |
| (5e-05,0.0001] | 343705 | 0.428 | 0.1 | 0.2 | 0.1x |
| (0.0001,0.0002] | 243964 | 0.471 | 0.101 | 0.2 | 0.1x |
| (0.0002,0.0005] | 158021 | 0.517 | 0.122 | 0.2 | 0.1x |
| (0.0005,0.001] | 62682 | 0.556 | 0.175 | 0.2 | 0.1x |
| (0.001,0.002] | 44870 | 0.578 | 0.227 | 0.2 | 0.1x |
| (0.002,0.005] | 44852 | 0.638 | 0.264 | 0.2 | 0.1x |
| (0.005,0.01] | 27163 | 0.674 | 0.251 | 0.2 | 0.1x |
| (0.01,0.02] | 23822 | 0.721 | 0.314 | 0.2 | 0.1x |
| (0.02,0.05] | 30504 | 0.776 | 0.364 | 0.2 | 0.1x |
| (0.05,0.1] | 23983 | 0.791 | 0.384 | 0.2 | 0.1x |
| (0.1,0.2] | 28254 | 0.811 | 0.412 | 0.2 | 0.1x |
| (0.2,0.3] | 18631 | 0.811 | 0.389 | 0.2 | 0.1x |
| (0.3,0.4] | 13490 | 0.82 | 0.385 | 0.2 | 0.1x |
| (0.4,0.5] | 11630 | 0.818 | 0.395 | 0.2 | 0.1x |
| (1e-05,2e-05] | 717636 | 0.305 | 0.126 | 0.3 | 0.1x |
| (2e-05,5e-05] | 619703 | 0.354 | 0.129 | 0.3 | 0.1x |
| (5e-05,0.0001] | 343705 | 0.361 | 0.125 | 0.3 | 0.1x |
| (0.0001,0.0002] | 243964 | 0.41 | 0.134 | 0.3 | 0.1x |
| (0.0002,0.0005] | 158021 | 0.47 | 0.173 | 0.3 | 0.1x |
| (0.0005,0.001] | 62682 | 0.487 | 0.225 | 0.3 | 0.1x |
| (0.001,0.002] | 44870 | 0.493 | 0.296 | 0.3 | 0.1x |
| (0.002,0.005] | 44852 | 0.568 | 0.328 | 0.3 | 0.1x |
| (0.005,0.01] | 27163 | 0.59 | 0.334 | 0.3 | 0.1x |
| (0.01,0.02] | 23822 | 0.635 | 0.392 | 0.3 | 0.1x |
| (0.02,0.05] | 30504 | 0.702 | 0.433 | 0.3 | 0.1x |
| (0.05,0.1] | 23983 | 0.719 | 0.463 | 0.3 | 0.1x |
| (0.1,0.2] | 28254 | 0.739 | 0.491 | 0.3 | 0.1x |
| (0.2,0.3] | 18631 | 0.739 | 0.468 | 0.3 | 0.1x |
| (0.3,0.4] | 13490 | 0.745 | 0.468 | 0.3 | 0.1x |
| (0.4,0.5] | 11630 | 0.746 | 0.48 | 0.3 | 0.1x |
| (1e-05,2e-05] | 717636 | 0.616 | 0.148 | 0.05 | 0.25x |
| (2e-05,5e-05] | 619703 | 0.7 | 0.12 | 0.05 | 0.25x |
| (5e-05,0.0001] | 343705 | 0.712 | 0.122 | 0.05 | 0.25x |
| (0.0001,0.0002] | 243964 | 0.744 | 0.132 | 0.05 | 0.25x |
| (0.0002,0.0005] | 158021 | 0.806 | 0.138 | 0.05 | 0.25x |
| (0.0005,0.001] | 62682 | 0.854 | 0.149 | 0.05 | 0.25x |
| (0.001,0.002] | 44870 | 0.862 | 0.212 | 0.05 | 0.25x |
| (0.002,0.005] | 44852 | 0.901 | 0.228 | 0.05 | 0.25x |
| (0.005,0.01] | 27163 | 0.916 | 0.227 | 0.05 | 0.25x |
| (0.01,0.02] | 23822 | 0.939 | 0.288 | 0.05 | 0.25x |
| (0.02,0.05] | 30504 | 0.957 | 0.306 | 0.05 | 0.25x |
| (0.05,0.1] | 23983 | 0.971 | 0.306 | 0.05 | 0.25x |

Continued on next page

Continued from previous page

| MAF | #SNPs | Mother | Fetus | FF | COV |
|---|---|---|---|---|---|
| (0.1,0.2] | 28254 | 0.976 | 0.331 | 0.05 | 0.25x |
| (0.2,0.3] | 18631 | 0.976 | 0.317 | 0.05 | 0.25x |
| (0.3,0.4] | 13490 | 0.978 | 0.306 | 0.05 | 0.25x |
| (0.4,0.5] | 11630 | 0.976 | 0.306 | 0.05 | 0.25x |
| (1e-05,2e-05] | 717636 | 0.606 | 0.175 | 0.1 | 0.25x |
| (2e-05,5e-05] | 619703 | 0.686 | 0.175 | 0.1 | 0.25x |
| (5e-05,0.0001] | 343705 | 0.685 | 0.165 | 0.1 | 0.25x |
| (0.0001,0.0002] | 243964 | 0.724 | 0.18 | 0.1 | 0.25x |
| (0.0002,0.0005] | 158021 | 0.788 | 0.186 | 0.1 | 0.25x |
| (0.0005,0.001] | 62682 | 0.827 | 0.219 | 0.1 | 0.25x |
| (0.001,0.002] | 44870 | 0.841 | 0.283 | 0.1 | 0.25x |
| (0.002,0.005] | 44852 | 0.885 | 0.316 | 0.1 | 0.25x |
| (0.005,0.01] | 27163 | 0.9 | 0.325 | 0.1 | 0.25x |
| (0.01,0.02] | 23822 | 0.921 | 0.378 | 0.1 | 0.25x |
| (0.02,0.05] | 30504 | 0.949 | 0.415 | 0.1 | 0.25x |
| (0.05,0.1] | 23983 | 0.961 | 0.451 | 0.1 | 0.25x |
| (0.1,0.2] | 28254 | 0.966 | 0.465 | 0.1 | 0.25x |
| (0.2,0.3] | 18631 | 0.965 | 0.442 | 0.1 | 0.25x |
| (0.3,0.4] | 13490 | 0.966 | 0.433 | 0.1 | 0.25x |
| (0.4,0.5] | 11630 | 0.966 | 0.433 | 0.1 | 0.25x |
| (1e-05,2e-05] | 717636 | 0.592 | 0.226 | 0.15 | 0.25x |
| (2e-05,5e-05] | 619703 | 0.67 | 0.244 | 0.15 | 0.25x |
| (5e-05,0.0001] | 343705 | 0.666 | 0.232 | 0.15 | 0.25x |
| (0.0001,0.0002] | 243964 | 0.709 | 0.251 | 0.15 | 0.25x |
| (0.0002,0.0005] | 158021 | 0.773 | 0.266 | 0.15 | 0.25x |
| (0.0005,0.001] | 62682 | 0.808 | 0.297 | 0.15 | 0.25x |
| (0.001,0.002] | 44870 | 0.828 | 0.38 | 0.15 | 0.25x |
| (0.002,0.005] | 44852 | 0.869 | 0.392 | 0.15 | 0.25x |
| (0.005,0.01] | 27163 | 0.883 | 0.423 | 0.15 | 0.25x |
| (0.01,0.02] | 23822 | 0.912 | 0.493 | 0.15 | 0.25x |
| (0.02,0.05] | 30504 | 0.938 | 0.523 | 0.15 | 0.25x |
| (0.05,0.1] | 23983 | 0.95 | 0.55 | 0.15 | 0.25x |
| (0.1,0.2] | 28254 | 0.956 | 0.571 | 0.15 | 0.25x |
| (0.2,0.3] | 18631 | 0.955 | 0.543 | 0.15 | 0.25x |
| (0.3,0.4] | 13490 | 0.956 | 0.543 | 0.15 | 0.25x |
| (0.4,0.5] | 11630 | 0.955 | 0.539 | 0.15 | 0.25x |
| (1e-05,2e-05] | 717636 | 0.561 | 0.278 | 0.2 | 0.25x |
| (2e-05,5e-05] | 619703 | 0.646 | 0.308 | 0.2 | 0.25x |
| (5e-05,0.0001] | 343705 | 0.643 | 0.296 | 0.2 | 0.25x |
| (0.0001,0.0002] | 243964 | 0.691 | 0.307 | 0.2 | 0.25x |
| (0.0002,0.0005] | 158021 | 0.751 | 0.354 | 0.2 | 0.25x |
| (0.0005,0.001] | 62682 | 0.789 | 0.41 | 0.2 | 0.25x |
| (0.001,0.002] | 44870 | 0.801 | 0.48 | 0.2 | 0.25x |
| (0.002,0.005] | 44852 | 0.849 | 0.5 | 0.2 | 0.25x |
| (0.005,0.01] | 27163 | 0.86 | 0.523 | 0.2 | 0.25x |
| (0.01,0.02] | 23822 | 0.887 | 0.59 | 0.2 | 0.25x |
| (0.02,0.05] | 30504 | 0.923 | 0.624 | 0.2 | 0.25x |
| (0.05,0.1] | 23983 | 0.935 | 0.65 | 0.2 | 0.25x |

Continued on next page

Continued from previous page

| MAF | #SNPs | Mother | Fetus | FF | COV |
|---|---|---|---|---|---|
| (0.1,0.2] | 28254 | 0.941 | 0.657 | 0.2 | 0.25x |
| (0.2,0.3] | 18631 | 0.94 | 0.637 | 0.2 | 0.25x |
| (0.3,0.4] | 13490 | 0.943 | 0.639 | 0.2 | 0.25x |
| (0.4,0.5] | 11630 | 0.941 | 0.635 | 0.2 | 0.25x |
| (1e-05,2e-05] | 717636 | 0.533 | 0.319 | 0.3 | 0.25x |
| (2e-05,5e-05] | 619703 | 0.589 | 0.365 | 0.3 | 0.25x |
| (5e-05,0.0001] | 343705 | 0.594 | 0.375 | 0.3 | 0.25x |
| (0.0001,0.0002] | 243964 | 0.643 | 0.405 | 0.3 | 0.25x |
| (0.0002,0.0005] | 158021 | 0.704 | 0.431 | 0.3 | 0.25x |
| (0.0005,0.001] | 62682 | 0.737 | 0.488 | 0.3 | 0.25x |
| (0.001,0.002] | 44870 | 0.756 | 0.56 | 0.3 | 0.25x |
| (0.002,0.005] | 44852 | 0.794 | 0.591 | 0.3 | 0.25x |
| (0.005,0.01] | 27163 | 0.812 | 0.609 | 0.3 | 0.25x |
| (0.01,0.02] | 23822 | 0.843 | 0.659 | 0.3 | 0.25x |
| (0.02,0.05] | 30504 | 0.881 | 0.697 | 0.3 | 0.25x |
| (0.05,0.1] | 23983 | 0.893 | 0.731 | 0.3 | 0.25x |
| (0.1,0.2] | 28254 | 0.9 | 0.738 | 0.3 | 0.25x |
| (0.2,0.3] | 18631 | 0.898 | 0.723 | 0.3 | 0.25x |
| (0.3,0.4] | 13490 | 0.901 | 0.724 | 0.3 | 0.25x |
| (0.4,0.5] | 11630 | 0.899 | 0.721 | 0.3 | 0.25x |
| (1e-05,2e-05] | 717636 | 0.676 | 0.235 | 0.05 | 1x |
| (2e-05,5e-05] | 619703 | 0.769 | 0.254 | 0.05 | 1x |
| (5e-05,0.0001] | 343705 | 0.796 | 0.235 | 0.05 | 1x |
| (0.0001,0.0002] | 243964 | 0.833 | 0.244 | 0.05 | 1x |
| (0.0002,0.0005] | 158021 | 0.884 | 0.272 | 0.05 | 1x |
| (0.0005,0.001] | 62682 | 0.926 | 0.33 | 0.05 | 1x |
| (0.001,0.002] | 44870 | 0.939 | 0.389 | 0.05 | 1x |
| (0.002,0.005] | 44852 | 0.966 | 0.393 | 0.05 | 1x |
| (0.005,0.01] | 27163 | 0.974 | 0.41 | 0.05 | 1x |
| (0.01,0.02] | 23822 | 0.983 | 0.464 | 0.05 | 1x |
| (0.02,0.05] | 30504 | 0.989 | 0.508 | 0.05 | 1x |
| (0.05,0.1] | 23983 | 0.993 | 0.539 | 0.05 | 1x |
| (0.1,0.2] | 28254 | 0.995 | 0.552 | 0.05 | 1x |
| (0.2,0.3] | 18631 | 0.995 | 0.53 | 0.05 | 1x |
| (0.3,0.4] | 13490 | 0.996 | 0.528 | 0.05 | 1x |
| (0.4,0.5] | 11630 | 0.995 | 0.52 | 0.05 | 1x |
| (1e-05,2e-05] | 717636 | 0.677 | 0.379 | 0.1 | 1x |
| (2e-05,5e-05] | 619703 | 0.769 | 0.422 | 0.1 | 1x |
| (5e-05,0.0001] | 343705 | 0.792 | 0.434 | 0.1 | 1x |
| (0.0001,0.0002] | 243964 | 0.831 | 0.458 | 0.1 | 1x |
| (0.0002,0.0005] | 158021 | 0.881 | 0.509 | 0.1 | 1x |
| (0.0005,0.001] | 62682 | 0.924 | 0.566 | 0.1 | 1x |
| (0.001,0.002] | 44870 | 0.935 | 0.641 | 0.1 | 1x |
| (0.002,0.005] | 44852 | 0.963 | 0.65 | 0.1 | 1x |
| (0.005,0.01] | 27163 | 0.972 | 0.678 | 0.1 | 1x |
| (0.01,0.02] | 23822 | 0.981 | 0.713 | 0.1 | 1x |
| (0.02,0.05] | 30504 | 0.988 | 0.745 | 0.1 | 1x |
| (0.05,0.1] | 23983 | 0.992 | 0.769 | 0.1 | 1x |

Continued on next page

Continued from previous page

| MAF | #SNPs | Mother | Fetus | FF | COV |
|---|---|---|---|---|---|
| (0.1,0.2] | 28254 | 0.993 | 0.778 | 0.1 | 1x |
| (0.2,0.3] | 18631 | 0.994 | 0.76 | 0.1 | 1x |
| (0.3,0.4] | 13490 | 0.994 | 0.759 | 0.1 | 1x |
| (0.4,0.5] | 11630 | 0.994 | 0.748 | 0.1 | 1x |
| (1e-05,2e-05] | 717636 | 0.673 | 0.472 | 0.15 | 1x |
| (2e-05,5e-05] | 619703 | 0.761 | 0.538 | 0.15 | 1x |
| (5e-05,0.0001] | 343705 | 0.786 | 0.539 | 0.15 | 1x |
| (0.0001,0.0002] | 243964 | 0.825 | 0.583 | 0.15 | 1x |
| (0.0002,0.0005] | 158021 | 0.881 | 0.647 | 0.15 | 1x |
| (0.0005,0.001] | 62682 | 0.923 | 0.698 | 0.15 | 1x |
| (0.001,0.002] | 44870 | 0.931 | 0.748 | 0.15 | 1x |
| (0.002,0.005] | 44852 | 0.959 | 0.765 | 0.15 | 1x |
| (0.005,0.01] | 27163 | 0.969 | 0.793 | 0.15 | 1x |
| (0.01,0.02] | 23822 | 0.979 | 0.819 | 0.15 | 1x |
| (0.02,0.05] | 30504 | 0.986 | 0.849 | 0.15 | 1x |
| (0.05,0.1] | 23983 | 0.991 | 0.868 | 0.15 | 1x |
| (0.1,0.2] | 28254 | 0.992 | 0.872 | 0.15 | 1x |
| (0.2,0.3] | 18631 | 0.992 | 0.858 | 0.15 | 1x |
| (0.3,0.4] | 13490 | 0.993 | 0.855 | 0.15 | 1x |
| (0.4,0.5] | 11630 | 0.992 | 0.85 | 0.15 | 1x |
| (1e-05,2e-05] | 717636 | 0.671 | 0.521 | 0.2 | 1x |
| (2e-05,5e-05] | 619703 | 0.761 | 0.581 | 0.2 | 1x |
| (5e-05,0.0001] | 343705 | 0.781 | 0.609 | 0.2 | 1x |
| (0.0001,0.0002] | 243964 | 0.815 | 0.646 | 0.2 | 1x |
| (0.0002,0.0005] | 158021 | 0.874 | 0.701 | 0.2 | 1x |
| (0.0005,0.001] | 62682 | 0.915 | 0.758 | 0.2 | 1x |
| (0.001,0.002] | 44870 | 0.926 | 0.799 | 0.2 | 1x |
| (0.002,0.005] | 44852 | 0.955 | 0.822 | 0.2 | 1x |
| (0.005,0.01] | 27163 | 0.965 | 0.85 | 0.2 | 1x |
| (0.01,0.02] | 23822 | 0.976 | 0.869 | 0.2 | 1x |
| (0.02,0.05] | 30504 | 0.984 | 0.893 | 0.2 | 1x |
| (0.05,0.1] | 23983 | 0.989 | 0.909 | 0.2 | 1x |
| (0.1,0.2] | 28254 | 0.99 | 0.913 | 0.2 | 1x |
| (0.2,0.3] | 18631 | 0.99 | 0.904 | 0.2 | 1x |
| (0.3,0.4] | 13490 | 0.99 | 0.901 | 0.2 | 1x |
| (0.4,0.5] | 11630 | 0.989 | 0.899 | 0.2 | 1x |
| (1e-05,2e-05] | 717636 | 0.654 | 0.551 | 0.3 | 1x |
| (2e-05,5e-05] | 619703 | 0.741 | 0.607 | 0.3 | 1x |
| (5e-05,0.0001] | 343705 | 0.76 | 0.645 | 0.3 | 1x |
| (0.0001,0.0002] | 243964 | 0.804 | 0.682 | 0.3 | 1x |
| (0.0002,0.0005] | 158021 | 0.86 | 0.745 | 0.3 | 1x |
| (0.0005,0.001] | 62682 | 0.895 | 0.794 | 0.3 | 1x |
| (0.001,0.002] | 44870 | 0.912 | 0.838 | 0.3 | 1x |
| (0.002,0.005] | 44852 | 0.941 | 0.868 | 0.3 | 1x |
| (0.005,0.01] | 27163 | 0.952 | 0.887 | 0.3 | 1x |
| (0.01,0.02] | 23822 | 0.965 | 0.901 | 0.3 | 1x |
| (0.02,0.05] | 30504 | 0.974 | 0.923 | 0.3 | 1x |
| (0.05,0.1] | 23983 | 0.981 | 0.937 | 0.3 | 1x |

Continued on next page

Continued from previous page

| MAF | #SNPs | Mother | Fetus | FF | COV |
|---|---|---|---|---|---|
| (0.1,0.2] | 28254 | 0.981 | 0.934 | 0.3 | 1x |
| (0.2,0.3] | 18631 | 0.981 | 0.929 | 0.3 | 1x |
| (0.3,0.4] | 13490 | 0.982 | 0.927 | 0.3 | 1x |
| (0.4,0.5] | 11630 | 0.98 | 0.925 | 0.3 | 1x |
| (1e-05,2e-05] | 717636 | 0.705 | 0.281 | 0.05 | 2x |
| (2e-05,5e-05] | 619703 | 0.791 | 0.355 | 0.05 | 2x |
| (5e-05,0.0001] | 343705 | 0.812 | 0.326 | 0.05 | 2x |
| (0.0001,0.0002] | 243964 | 0.857 | 0.343 | 0.05 | 2x |
| (0.0002,0.0005] | 158021 | 0.902 | 0.377 | 0.05 | 2x |
| (0.0005,0.001] | 62682 | 0.942 | 0.43 | 0.05 | 2x |
| (0.001,0.002] | 44870 | 0.952 | 0.522 | 0.05 | 2x |
| (0.002,0.005] | 44852 | 0.976 | 0.538 | 0.05 | 2x |
| (0.005,0.01] | 27163 | 0.983 | 0.549 | 0.05 | 2x |
| (0.01,0.02] | 23822 | 0.989 | 0.604 | 0.05 | 2x |
| (0.02,0.05] | 30504 | 0.993 | 0.635 | 0.05 | 2x |
| (0.05,0.1] | 23983 | 0.996 | 0.66 | 0.05 | 2x |
| (0.1,0.2] | 28254 | 0.996 | 0.666 | 0.05 | 2x |
| (0.2,0.3] | 18631 | 0.996 | 0.653 | 0.05 | 2x |
| (0.3,0.4] | 13490 | 0.997 | 0.649 | 0.05 | 2x |
| (0.4,0.5] | 11630 | 0.996 | 0.643 | 0.05 | 2x |
| (1e-05,2e-05] | 717636 | 0.705 | 0.478 | 0.1 | 2x |
| (2e-05,5e-05] | 619703 | 0.794 | 0.528 | 0.1 | 2x |
| (5e-05,0.0001] | 343705 | 0.814 | 0.537 | 0.1 | 2x |
| (0.0001,0.0002] | 243964 | 0.851 | 0.574 | 0.1 | 2x |
| (0.0002,0.0005] | 158021 | 0.899 | 0.63 | 0.1 | 2x |
| (0.0005,0.001] | 62682 | 0.941 | 0.683 | 0.1 | 2x |
| (0.001,0.002] | 44870 | 0.951 | 0.724 | 0.1 | 2x |
| (0.002,0.005] | 44852 | 0.975 | 0.74 | 0.1 | 2x |
| (0.005,0.01] | 27163 | 0.982 | 0.778 | 0.1 | 2x |
| (0.01,0.02] | 23822 | 0.988 | 0.809 | 0.1 | 2x |
| (0.02,0.05] | 30504 | 0.992 | 0.832 | 0.1 | 2x |
| (0.05,0.1] | 23983 | 0.995 | 0.851 | 0.1 | 2x |
| (0.1,0.2] | 28254 | 0.996 | 0.854 | 0.1 | 2x |
| (0.2,0.3] | 18631 | 0.995 | 0.846 | 0.1 | 2x |
| (0.3,0.4] | 13490 | 0.996 | 0.84 | 0.1 | 2x |
| (0.4,0.5] | 11630 | 0.996 | 0.839 | 0.1 | 2x |
| (1e-05,2e-05] | 717636 | 0.7 | 0.537 | 0.15 | 2x |
| (2e-05,5e-05] | 619703 | 0.787 | 0.615 | 0.15 | 2x |
| (5e-05,0.0001] | 343705 | 0.807 | 0.623 | 0.15 | 2x |
| (0.0001,0.0002] | 243964 | 0.851 | 0.661 | 0.15 | 2x |
| (0.0002,0.0005] | 158021 | 0.9 | 0.728 | 0.15 | 2x |
| (0.0005,0.001] | 62682 | 0.937 | 0.785 | 0.15 | 2x |
| (0.001,0.002] | 44870 | 0.948 | 0.817 | 0.15 | 2x |
| (0.002,0.005] | 44852 | 0.973 | 0.841 | 0.15 | 2x |
| (0.005,0.01] | 27163 | 0.981 | 0.867 | 0.15 | 2x |
| (0.01,0.02] | 23822 | 0.988 | 0.889 | 0.15 | 2x |
| (0.02,0.05] | 30504 | 0.991 | 0.906 | 0.15 | 2x |
| (0.05,0.1] | 23983 | 0.994 | 0.919 | 0.15 | 2x |

Continued on next page

Continued from previous page

| MAF | #SNPs | Mother | Fetus | FF | COV |
|---|---|---|---|---|---|
| (0.1,0.2] | 28254 | 0.995 | 0.917 | 0.15 | 2x |
| (0.2,0.3] | 18631 | 0.995 | 0.909 | 0.15 | 2x |
| (0.3,0.4] | 13490 | 0.995 | 0.907 | 0.15 | 2x |
| (0.4,0.5] | 11630 | 0.995 | 0.903 | 0.15 | 2x |
| (1e-05,2e-05] | 717636 | 0.698 | 0.573 | 0.2 | 2x |
| (2e-05,5e-05] | 619703 | 0.784 | 0.633 | 0.2 | 2x |
| (5e-05,0.0001] | 343705 | 0.805 | 0.661 | 0.2 | 2x |
| (0.0001,0.0002] | 243964 | 0.847 | 0.703 | 0.2 | 2x |
| (0.0002,0.0005] | 158021 | 0.896 | 0.761 | 0.2 | 2x |
| (0.0005,0.001] | 62682 | 0.934 | 0.814 | 0.2 | 2x |
| (0.001,0.002] | 44870 | 0.944 | 0.854 | 0.2 | 2x |
| (0.002,0.005] | 44852 | 0.971 | 0.867 | 0.2 | 2x |
| (0.005,0.01] | 27163 | 0.978 | 0.895 | 0.2 | 2x |
| (0.01,0.02] | 23822 | 0.986 | 0.917 | 0.2 | 2x |
| (0.02,0.05] | 30504 | 0.99 | 0.934 | 0.2 | 2x |
| (0.05,0.1] | 23983 | 0.993 | 0.943 | 0.2 | 2x |
| (0.1,0.2] | 28254 | 0.994 | 0.938 | 0.2 | 2x |
| (0.2,0.3] | 18631 | 0.993 | 0.932 | 0.2 | 2x |
| (0.3,0.4] | 13490 | 0.994 | 0.928 | 0.2 | 2x |
| (0.4,0.5] | 11630 | 0.993 | 0.925 | 0.2 | 2x |
| (1e-05,2e-05] | 717636 | 0.689 | 0.591 | 0.3 | 2x |
| (2e-05,5e-05] | 619703 | 0.774 | 0.649 | 0.3 | 2x |
| (5e-05,0.0001] | 343705 | 0.79 | 0.681 | 0.3 | 2x |
| (0.0001,0.0002] | 243964 | 0.835 | 0.731 | 0.3 | 2x |
| (0.0002,0.0005] | 158021 | 0.884 | 0.788 | 0.3 | 2x |
| (0.0005,0.001] | 62682 | 0.925 | 0.843 | 0.3 | 2x |
| (0.001,0.002] | 44870 | 0.934 | 0.879 | 0.3 | 2x |
| (0.002,0.005] | 44852 | 0.962 | 0.901 | 0.3 | 2x |
| (0.005,0.01] | 27163 | 0.97 | 0.921 | 0.3 | 2x |
| (0.01,0.02] | 23822 | 0.98 | 0.935 | 0.3 | 2x |
| (0.02,0.05] | 30504 | 0.985 | 0.951 | 0.3 | 2x |
| (0.05,0.1] | 23983 | 0.988 | 0.958 | 0.3 | 2x |
| (0.1,0.2] | 28254 | 0.988 | 0.953 | 0.3 | 2x |
| (0.2,0.3] | 18631 | 0.988 | 0.948 | 0.3 | 2x |
| (0.3,0.4] | 13490 | 0.988 | 0.946 | 0.3 | 2x |
| (0.4,0.5] | 11630 | 0.987 | 0.942 | 0.3 | 2x |
| (1e-05,2e-05] | 717636 | 0.725 | 0.358 | 0.05 | 4x |
| (2e-05,5e-05] | 619703 | 0.811 | 0.4 | 0.05 | 4x |
| (5e-05,0.0001] | 343705 | 0.83 | 0.396 | 0.05 | 4x |
| (0.0001,0.0002] | 243964 | 0.875 | 0.428 | 0.05 | 4x |
| (0.0002,0.0005] | 158021 | 0.915 | 0.477 | 0.05 | 4x |
| (0.0005,0.001] | 62682 | 0.952 | 0.534 | 0.05 | 4x |
| (0.001,0.002] | 44870 | 0.96 | 0.589 | 0.05 | 4x |
| (0.002,0.005] | 44852 | 0.984 | 0.605 | 0.05 | 4x |
| (0.005,0.01] | 27163 | 0.988 | 0.64 | 0.05 | 4x |
| (0.01,0.02] | 23822 | 0.992 | 0.683 | 0.05 | 4x |
| (0.02,0.05] | 30504 | 0.995 | 0.707 | 0.05 | 4x |
| (0.05,0.1] | 23983 | 0.997 | 0.734 | 0.05 | 4x |

Continued on next page

Continued from previous page

| MAF | #SNPs | Mother | Fetus | FF | COV |
|---|---|---|---|---|---|
| (0.1,0.2] | 28254 | 0.997 | 0.741 | 0.05 | 4x |
| (0.2,0.3] | 18631 | 0.997 | 0.73 | 0.05 | 4x |
| (0.3,0.4] | 13490 | 0.997 | 0.719 | 0.05 | 4x |
| (0.4,0.5] | 11630 | 0.997 | 0.721 | 0.05 | 4x |
| (1e-05,2e-05] | 717636 | 0.725 | 0.547 | 0.1 | 4x |
| (2e-05,5e-05] | 619703 | 0.814 | 0.588 | 0.1 | 4x |
| (5e-05,0.0001] | 343705 | 0.832 | 0.608 | 0.1 | 4x |
| (0.0001,0.0002] | 243964 | 0.875 | 0.628 | 0.1 | 4x |
| (0.0002,0.0005] | 158021 | 0.914 | 0.696 | 0.1 | 4x |
| (0.0005,0.001] | 62682 | 0.952 | 0.746 | 0.1 | 4x |
| (0.001,0.002] | 44870 | 0.958 | 0.79 | 0.1 | 4x |
| (0.002,0.005] | 44852 | 0.982 | 0.816 | 0.1 | 4x |
| (0.005,0.01] | 27163 | 0.988 | 0.848 | 0.1 | 4x |
| (0.01,0.02] | 23822 | 0.992 | 0.866 | 0.1 | 4x |
| (0.02,0.05] | 30504 | 0.995 | 0.888 | 0.1 | 4x |
| (0.05,0.1] | 23983 | 0.997 | 0.901 | 0.1 | 4x |
| (0.1,0.2] | 28254 | 0.996 | 0.894 | 0.1 | 4x |
| (0.2,0.3] | 18631 | 0.996 | 0.887 | 0.1 | 4x |
| (0.3,0.4] | 13490 | 0.997 | 0.884 | 0.1 | 4x |
| (0.4,0.5] | 11630 | 0.996 | 0.878 | 0.1 | 4x |
| (1e-05,2e-05] | 717636 | 0.723 | 0.569 | 0.15 | 4x |
| (2e-05,5e-05] | 619703 | 0.809 | 0.628 | 0.15 | 4x |
| (5e-05,0.0001] | 343705 | 0.823 | 0.66 | 0.15 | 4x |
| (0.0001,0.0002] | 243964 | 0.871 | 0.694 | 0.15 | 4x |
| (0.0002,0.0005] | 158021 | 0.911 | 0.747 | 0.15 | 4x |
| (0.0005,0.001] | 62682 | 0.949 | 0.806 | 0.15 | 4x |
| (0.001,0.002] | 44870 | 0.957 | 0.837 | 0.15 | 4x |
| (0.002,0.005] | 44852 | 0.982 | 0.847 | 0.15 | 4x |
| (0.005,0.01] | 27163 | 0.987 | 0.892 | 0.15 | 4x |
| (0.01,0.02] | 23822 | 0.991 | 0.903 | 0.15 | 4x |
| (0.02,0.05] | 30504 | 0.994 | 0.927 | 0.15 | 4x |
| (0.05,0.1] | 23983 | 0.996 | 0.93 | 0.15 | 4x |
| (0.1,0.2] | 28254 | 0.995 | 0.929 | 0.15 | 4x |
| (0.2,0.3] | 18631 | 0.995 | 0.92 | 0.15 | 4x |
| (0.3,0.4] | 13490 | 0.995 | 0.918 | 0.15 | 4x |
| (0.4,0.5] | 11630 | 0.995 | 0.911 | 0.15 | 4x |
| (1e-05,2e-05] | 717636 | 0.72 | 0.604 | 0.2 | 4x |
| (2e-05,5e-05] | 619703 | 0.807 | 0.658 | 0.2 | 4x |
| (5e-05,0.0001] | 343705 | 0.819 | 0.685 | 0.2 | 4x |
| (0.0001,0.0002] | 243964 | 0.868 | 0.731 | 0.2 | 4x |
| (0.0002,0.0005] | 158021 | 0.909 | 0.779 | 0.2 | 4x |
| (0.0005,0.001] | 62682 | 0.945 | 0.836 | 0.2 | 4x |
| (0.001,0.002] | 44870 | 0.952 | 0.869 | 0.2 | 4x |
| (0.002,0.005] | 44852 | 0.979 | 0.887 | 0.2 | 4x |
| (0.005,0.01] | 27163 | 0.986 | 0.917 | 0.2 | 4x |
| (0.01,0.02] | 23822 | 0.989 | 0.934 | 0.2 | 4x |
| (0.02,0.05] | 30504 | 0.992 | 0.946 | 0.2 | 4x |
| (0.05,0.1] | 23983 | 0.994 | 0.952 | 0.2 | 4x |

Continued on next page

Continued from previous page

| MAF | #SNPs | Mother | Fetus | FF | COV |
|---|---|---|---|---|---|
| (0.1,0.2] | 28254 | 0.994 | 0.944 | 0.2 | 4x |
| (0.2,0.3] | 18631 | 0.993 | 0.936 | 0.2 | 4x |
| (0.3,0.4] | 13490 | 0.994 | 0.934 | 0.2 | 4x |
| (0.4,0.5] | 11630 | 0.993 | 0.927 | 0.2 | 4x |
| (1e-05,2e-05] | 717636 | 0.7 | 0.622 | 0.3 | 4x |
| (2e-05,5e-05] | 619703 | 0.793 | 0.676 | 0.3 | 4x |
| (5e-05,0.0001] | 343705 | 0.807 | 0.708 | 0.3 | 4x |
| (0.0001,0.0002] | 243964 | 0.859 | 0.755 | 0.3 | 4x |
| (0.0002,0.0005] | 158021 | 0.898 | 0.81 | 0.3 | 4x |
| (0.0005,0.001] | 62682 | 0.935 | 0.856 | 0.3 | 4x |
| (0.001,0.002] | 44870 | 0.946 | 0.896 | 0.3 | 4x |
| (0.002,0.005] | 44852 | 0.97 | 0.915 | 0.3 | 4x |
| (0.005,0.01] | 27163 | 0.977 | 0.933 | 0.3 | 4x |
| (0.01,0.02] | 23822 | 0.984 | 0.945 | 0.3 | 4x |
| (0.02,0.05] | 30504 | 0.987 | 0.958 | 0.3 | 4x |
| (0.05,0.1] | 23983 | 0.989 | 0.962 | 0.3 | 4x |
| (0.1,0.2] | 28254 | 0.988 | 0.955 | 0.3 | 4x |
| (0.2,0.3] | 18631 | 0.988 | 0.949 | 0.3 | 4x |
| (0.3,0.4] | 13490 | 0.988 | 0.946 | 0.3 | 4x |
| (0.4,0.5] | 11630 | 0.987 | 0.941 | 0.3 | 4x |
| (1e-05,2e-05] | 717636 | 0.759 | 0.431 | 0.05 | 10x |
| (2e-05,5e-05] | 619703 | 0.836 | 0.469 | 0.05 | 10x |
| (5e-05,0.0001] | 343705 | 0.851 | 0.48 | 0.05 | 10x |
| (0.0001,0.0002] | 243964 | 0.896 | 0.503 | 0.05 | 10x |
| (0.0002,0.0005] | 158021 | 0.93 | 0.556 | 0.05 | 10x |
| (0.0005,0.001] | 62682 | 0.959 | 0.603 | 0.05 | 10x |
| (0.001,0.002] | 44870 | 0.965 | 0.665 | 0.05 | 10x |
| (0.002,0.005] | 44852 | 0.987 | 0.682 | 0.05 | 10x |
| (0.005,0.01] | 27163 | 0.993 | 0.737 | 0.05 | 10x |
| (0.01,0.02] | 23822 | 0.993 | 0.766 | 0.05 | 10x |
| (0.02,0.05] | 30504 | 0.996 | 0.795 | 0.05 | 10x |
| (0.05,0.1] | 23983 | 0.998 | 0.809 | 0.05 | 10x |
| (0.1,0.2] | 28254 | 0.997 | 0.804 | 0.05 | 10x |
| (0.2,0.3] | 18631 | 0.997 | 0.794 | 0.05 | 10x |
| (0.3,0.4] | 13490 | 0.998 | 0.791 | 0.05 | 10x |
| (0.4,0.5] | 11630 | 0.997 | 0.782 | 0.05 | 10x |
| (1e-05,2e-05] | 717636 | 0.755 | 0.534 | 0.1 | 10x |
| (2e-05,5e-05] | 619703 | 0.832 | 0.581 | 0.1 | 10x |
| (5e-05,0.0001] | 343705 | 0.843 | 0.61 | 0.1 | 10x |
| (0.0001,0.0002] | 243964 | 0.889 | 0.629 | 0.1 | 10x |
| (0.0002,0.0005] | 158021 | 0.925 | 0.697 | 0.1 | 10x |
| (0.0005,0.001] | 62682 | 0.957 | 0.751 | 0.1 | 10x |
| (0.001,0.002] | 44870 | 0.963 | 0.803 | 0.1 | 10x |
| (0.002,0.005] | 44852 | 0.986 | 0.819 | 0.1 | 10x |
| (0.005,0.01] | 27163 | 0.991 | 0.864 | 0.1 | 10x |
| (0.01,0.02] | 23822 | 0.992 | 0.885 | 0.1 | 10x |
| (0.02,0.05] | 30504 | 0.995 | 0.902 | 0.1 | 10x |
| (0.05,0.1] | 23983 | 0.996 | 0.91 | 0.1 | 10x |

Continued on next page

Continued from previous page

| MAF | #SNPs | Mother | Fetus | FF | COV |
|---|---|---|---|---|---|
| (0.1,0.2] | 28254 | 0.996 | 0.902 | 0.1 | 10x |
| (0.2,0.3] | 18631 | 0.996 | 0.892 | 0.1 | 10x |
| (0.3,0.4] | 13490 | 0.996 | 0.885 | 0.1 | 10x |
| (0.4,0.5] | 11630 | 0.995 | 0.877 | 0.1 | 10x |
| (1e-05,2e-05] | 717636 | 0.748 | 0.581 | 0.15 | 10x |
| (2e-05,5e-05] | 619703 | 0.822 | 0.619 | 0.15 | 10x |
| (5e-05,0.0001] | 343705 | 0.836 | 0.656 | 0.15 | 10x |
| (0.0001,0.0002] | 243964 | 0.885 | 0.689 | 0.15 | 10x |
| (0.0002,0.0005] | 158021 | 0.921 | 0.74 | 0.15 | 10x |
| (0.0005,0.001] | 62682 | 0.952 | 0.801 | 0.15 | 10x |
| (0.001,0.002] | 44870 | 0.959 | 0.849 | 0.15 | 10x |
| (0.002,0.005] | 44852 | 0.983 | 0.857 | 0.15 | 10x |
| (0.005,0.01] | 27163 | 0.989 | 0.899 | 0.15 | 10x |
| (0.01,0.02] | 23822 | 0.991 | 0.913 | 0.15 | 10x |
| (0.02,0.05] | 30504 | 0.993 | 0.93 | 0.15 | 10x |
| (0.05,0.1] | 23983 | 0.994 | 0.934 | 0.15 | 10x |
| (0.1,0.2] | 28254 | 0.993 | 0.922 | 0.15 | 10x |
| (0.2,0.3] | 18631 | 0.993 | 0.913 | 0.15 | 10x |
| (0.3,0.4] | 13490 | 0.993 | 0.91 | 0.15 | 10x |
| (0.4,0.5] | 11630 | 0.992 | 0.899 | 0.15 | 10x |
| (1e-05,2e-05] | 717636 | 0.745 | 0.609 | 0.2 | 10x |
| (2e-05,5e-05] | 619703 | 0.817 | 0.641 | 0.2 | 10x |
| (5e-05,0.0001] | 343705 | 0.829 | 0.67 | 0.2 | 10x |
| (0.0001,0.0002] | 243964 | 0.879 | 0.704 | 0.2 | 10x |
| (0.0002,0.0005] | 158021 | 0.917 | 0.767 | 0.2 | 10x |
| (0.0005,0.001] | 62682 | 0.949 | 0.815 | 0.2 | 10x |
| (0.001,0.002] | 44870 | 0.954 | 0.862 | 0.2 | 10x |
| (0.002,0.005] | 44852 | 0.98 | 0.881 | 0.2 | 10x |
| (0.005,0.01] | 27163 | 0.985 | 0.91 | 0.2 | 10x |
| (0.01,0.02] | 23822 | 0.988 | 0.922 | 0.2 | 10x |
| (0.02,0.05] | 30504 | 0.99 | 0.937 | 0.2 | 10x |
| (0.05,0.1] | 23983 | 0.991 | 0.941 | 0.2 | 10x |
| (0.1,0.2] | 28254 | 0.99 | 0.929 | 0.2 | 10x |
| (0.2,0.3] | 18631 | 0.989 | 0.919 | 0.2 | 10x |
| (0.3,0.4] | 13490 | 0.989 | 0.915 | 0.2 | 10x |
| (0.4,0.5] | 11630 | 0.988 | 0.909 | 0.2 | 10x |
| (1e-05,2e-05] | 717636 | 0.726 | 0.633 | 0.3 | 10x |
| (2e-05,5e-05] | 619703 | 0.8 | 0.661 | 0.3 | 10x |
| (5e-05,0.0001] | 343705 | 0.811 | 0.687 | 0.3 | 10x |
| (0.0001,0.0002] | 243964 | 0.863 | 0.721 | 0.3 | 10x |
| (0.0002,0.0005] | 158021 | 0.901 | 0.777 | 0.3 | 10x |
| (0.0005,0.001] | 62682 | 0.935 | 0.835 | 0.3 | 10x |
| (0.001,0.002] | 44870 | 0.937 | 0.877 | 0.3 | 10x |
| (0.002,0.005] | 44852 | 0.966 | 0.897 | 0.3 | 10x |
| (0.005,0.01] | 27163 | 0.972 | 0.92 | 0.3 | 10x |
| (0.01,0.02] | 23822 | 0.977 | 0.928 | 0.3 | 10x |
| (0.02,0.05] | 30504 | 0.979 | 0.942 | 0.3 | 10x |
| (0.05,0.1] | 23983 | 0.981 | 0.944 | 0.3 | 10x |

Continued on next page

Continued from previous page

| MAF | #SNPs | Mother | Fetus | FF | COV |
|---|---|---|---|---|---|
| (0.1,0.2] | 28254 | 0.98 | 0.937 | 0.3 | 10x |
| (0.2,0.3] | 18631 | 0.979 | 0.928 | 0.3 | 10x |
| (0.3,0.4] | 13490 | 0.978 | 0.925 | 0.3 | 10x |
| (0.4,0.5] | 11630 | 0.977 | 0.921 | 0.3 | 10x |
